# The adaptive value of tandem communication in ants: insights from an agent-based model

**DOI:** 10.1101/2020.09.14.296426

**Authors:** Natascha Goy, Simone M. Glaser, Christoph Grüter

## Abstract

Social animals often share information about the location of resources, such as a food source or a new nest-site. One well-studied communication strategy in ants is tandem running, whereby a leader guides a recruit to a resource. Tandem running is considered an example of animal teaching because a leader adjusts her behaviour and invests time to help another ant to learn the location of a resource more efficiently. Tandem running also has costs, such as waiting inside the nest for a leader and a reduced walking speed. Whether and when these costs outweigh the benefits of tandem running is not well understood. We developed an agent-based simulation model to investigate the conditions that favour communication by tandem running during foraging. We predicted that the spatio-temporal distribution of food sources, colony size and the ratio of scouts and recruits affect colony foraging success. Our results suggest that communication is favoured when food sources are hard to find, of variable quality and long lasting. These results mirror the findings of simulations of honeybee communication. Scouts locate food sources faster than tandem followers in some environments, suggesting that tandem running may fulfil the criteria of teaching only in some situations. Furthermore, tandem running was only beneficial above a critical colony size threshold. Taken together, our model suggests that there is a considerable parameter range that favours colonies that do not use communication, which could explain why many social insects with small colony sizes forage solitarily.

## Introduction

Finding food is critical for survival and reproduction, but also energy- and time-consuming. Foraging for food can be done independently or by using information provided by other organisms (Sumpter 2010; Hoppitt and Laland 2013). In social insects, such as ants, social bees or social wasps, new food sources are usually discovered by scouts that explore the environment on their own (Hölldobler and Wilson 1990; Seeley 1995). After finding a profitable food source, they return to their nest and often communicate their discovery to nestmates. The communicated information depends on the species, but often includes the location of the resource, e.g. by means of laying a pheromone trail (Hölldobler and Wilson 1990; Jarau and Hrncir 2009; Czaczkes et al. 2015b). Recruitment communication allows colonies to exploit profitable feeding sites fast, e.g. before competitors have discovered and consumed the food source. Once the foragers have learned the location of the food source, they can use their route memory to return to the feeding site (e.g. von Frisch 1967; Collett et al. 2013).

In social insects, foraging strategies should not only take into account short-term individual success, but also how they affect colony foraging success. Thus, the value of communication should ultimately be studied at the colony level. So far, most theoretical and empirical studies that explored the value of communication for colony foraging success have focused on honeybees (but see also e.g. Sumpter and Pratt 2003; Dechaume-Moncharmont et al. 2005; Czaczkes et al. 2015a). These studies suggest that the value of communicating the location of food sources by means of waggle dances depends on how food sources are distributed (Sherman and Visscher 2002; Dornhaus and Chittka 2004; Dornhaus et al. 2006; Beekman and Lew 2008; Donaldson-Matasci and Dornhaus 2012; Schürch and Grüter 2014; I’Anson Price et al. 2019; reviewed in I’Anson Price and Grüter 2015). For example, Beekman & Lew (2008) found that the value of the “dance language” (the spatial information provided by the waggle dance) depends on the size and distance of the food patches. When patches were large and close to the hive, colonies that did not use dance communication and instead followed an individual foraging strategy were more successful. Dornhaus et al. (2006) concluded that dance communication does not help colonies collect more energy if there are many food sources that vary little in quality. Their models suggest that communication is beneficial if high-quality food sources are available, but are hard to find and that dance communication could be detrimental if food sources are easy to find (see also Dechaume-Moncharmont et al. 2005). In the latter case, foragers should search for new food sources through scouting (independent search) and return to known high-quality food sources using route memory (Schürch and Grüter 2014).

There is a well-known, but not yet fully understood link between colony size and the method of recruitment in ants (Beckers et al. 1989; Planqué et al. 2010; Dornhaus et al. 2012). While large colony size is associated with pheromone-based mass-recruitment, species with smaller colony sizes often forage solitarily or they use a recruitment method called tandem running (Beckers et al. 1989). In tandem running, an experienced ant (tandem leader) guides an inexperienced nestmate (tandem follower) to a new nest-site or a rewarding food source (Hingston 1929; Wilson 1959; Möglich et al. 1974; Franks and Richardson 2006; Pratt 2008; Kaur et al. 2017; reviewed in Franklin 2014). It has been argued that tandem followers locate resources quicker than scouts that search for resources by individual exploration and trial-and-error learning (Franks and Richardson 2006). Additionally, ants that are recruited by a tandem leader might find food sources of higher quality because foragers are more likely to perform tandem runs after finding a better food source (Shaffer et al. 2013). On the other hand, tandem running also has disadvantages. During a tandem run, both ants walk with reduced speed (Franks and Richardson 2006; Kaur et al. 2017) and a substantial proportion of tandem runs fail (e.g. Wilson 1959; Pratt 2008; Glaser and Grüter 2018; Grüter et al. 2018). Furthermore, recruits experience time and opportunity costs as they wait inside their nest for a leader, rather than search in the environment for food sources by themselves. These disadvantages could explain why some ant species do not seem to use tandem communication when foraging, even though tandem runs are used during colony migrations (Hölldobler 1984; Traniello and Hölldobler 1984; Fresneau 1985; Maschwitz et al. 1986). More generally, a sizeable group of ant species do not seem to use any form of communication during foraging (e.g. Beckers et al. 1989; Lanan 2014; Reeves and Moreau 2019). This raises the question whether, when and how a communication method that is relatively slow and small-scale-, like tandem running, improves colony foraging success and whether the ecological circumstances that favour tandem running match those that favour honeybee dance communication.

We developed an agent-based simulation model to investigate the importance of recruitment communication in the form of tandem running for the foraging success of virtual ant colonies. We compared colonies that could perform tandem runs with colonies that consisted only of scouts, *i.e*. without tandem running in an environment that varied in the number, quality, distance and longevity of food sources. Additionally, we tested whether colony size affects the importance of communication for colony foraging success. Finally, we explored the role of forager ratio (relative numbers of scouts and recruits) and tested if recruits indeed locate food sources faster than scouts. Based on studies that simulated honeybee foraging, we predicted that tandem running is beneficial when high quality food sources are hard to find (Dornhaus et al. 2006; Beekman and Lew 2008), but is detrimental to colony success when food sources are short-lived (Schürch and Grüter 2014). We also predicted that larger colonies benefit more from tandem running.

### The agent-based simulation model

An agent-based simulation model (ABM) was developed using the software Netlogo 6.1.1 (Wilensky 1999, Wilensky and Rand 2015) (the NetLogo file can be found in the online material). The model simulates the foragers of an artificial ant colony in an environment consisting of their nest and food sources. Some of the basic parameters, like the range of colony sizes, walking speeds or energy collected by foragers were derived from the ant species *Temnothorax nylanderi* (Glaser and Grüter 2018).

#### Purpose

The aim of our model was to explore the adaptive value of tandem running in ants by measuring the colony foraging success (as gained energy) and the time required by foragers to find a food source. We compared colonies that could perform tandem runs with colonies that consisted only of scouts, *i.e*. without tandem running. This latter situation is found in many ant species with small colony sizes, such as *Diacamma* or *Neoponera* (Hölldobler 1984, Traniello and Hölldobler 1984, Fresneau 1985, R. Kaur, pers. communication). In both situations, foragers could also use route memory (or private information) to return to food sources they visited in the past. We assessed the effects of communication depending on food source distribution (number, distance), their quality and stability as well as colony size and the scout-recruit ratio.

#### Entities, state variables and scale

Netlogo operates with patches that can be used to measure distances and ticks for time steps. In our model, 1 tick is equivalent to 1 second and 1 patch to 1 cm. The agents operate in a two-dimensional square grid of 140×140 patches (arena) with a nest and either 2 or 10 food sources (FS). This simulated environments with few or many food sources. The nest is located in the center (x=0, y=0), with a radius of 10 patches. The food patches were at a distance of either 40 (default) or 20 patches from the outer edge of the nest, simulating natural conditions as *T. nylanderi* mostly forages within 50 cm from their nest (Heinze et al. 1996). Each food source had a size of 1 patch, which could represent a dead insect or a drop of honey dew, and could either be of high or low quality (FS_high_ and FS_low_), simulating a sugar solution of either 1 molar or 0.1 molar concentration.

Since all agents are foragers, our default colony size of 100 would correspond to a natural colony consisting of ~300-400 workers, assuming that foragers account for about 20-30% of a *Temnothorax* colony (e.g. Shaffer et al. 2013). Simulated colonies consisted of varying ratios of scouts that search for resources independently and recruits that waited in the nest until they are recruited to a food source. The default scout-recruit ratio was 1:4 (*i.e*. 20 scouts + 80 recruits in the default situation), similar to what has been observed in honeybees where scouts represent about 5-35 % of the colony (von Frisch 1967; Seeley 1995). In colonies without tandem running, all foragers were scouts. In the default configuration, scouts and recruits can both assume any of the following seven states: (*1*) idle inside the nest, (*2*) feeding at food sources, (*3*) returning to the nest with food, (*4*) unloading food, (*5*) searching for a follower inside the nest, (*6*) leading a tandem run to the food source or (*7*) returning alone to the food source (*i.e*. use private information). Additionally, scouts search for food sources independently, while recruits wait inside the nest for a tandem leader. Recruits can then follow tandem runs to a food source (Fig. 1).

**Fig. 1.**
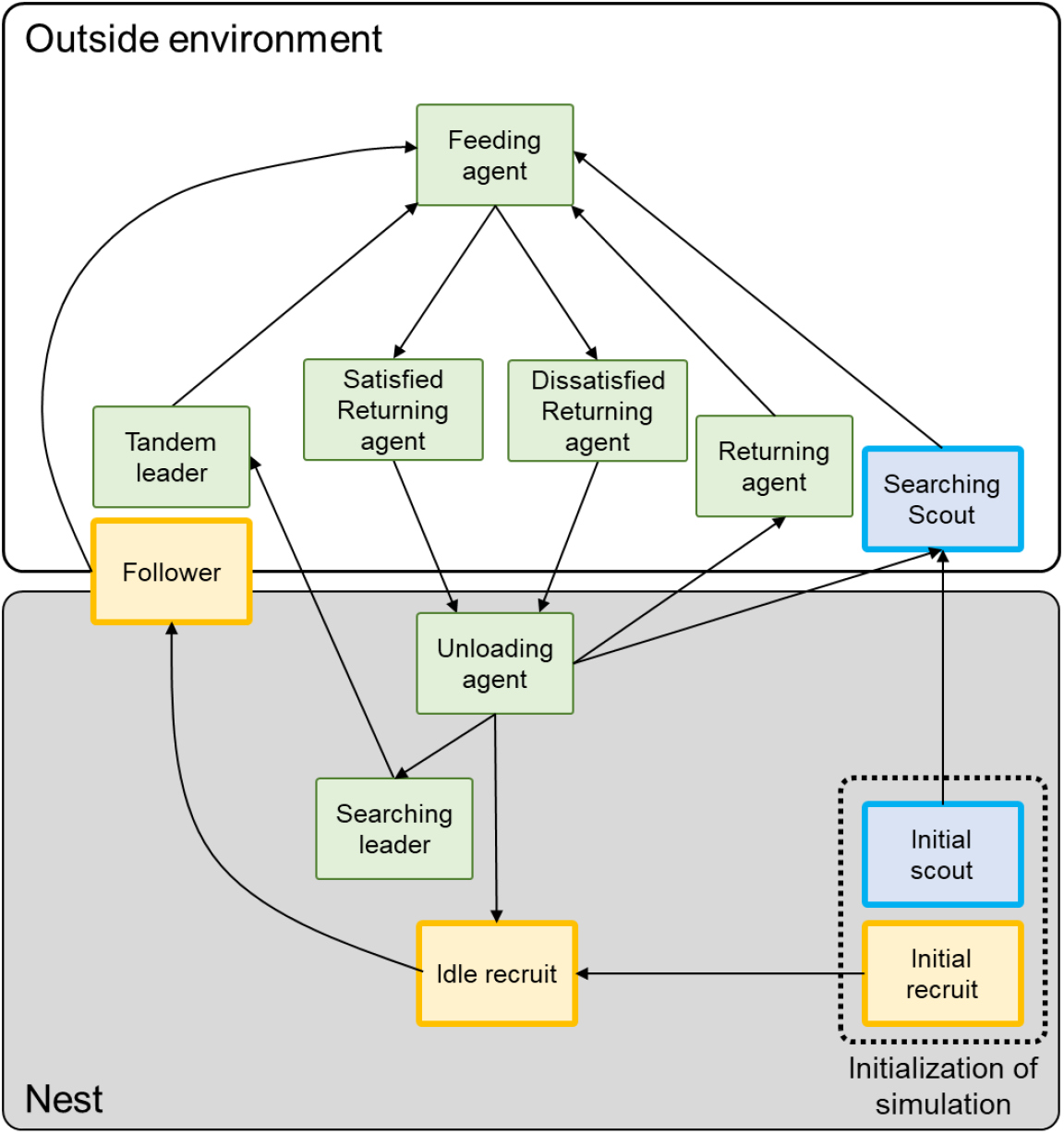
State diagram for the agent-based model for colonies with tandem runs. All foragers start inside the nest. Green boxes represent agent states that are possible for both scouts and recruits, blue boxes are states that are only possible for scouts and orange boxes represent states that are only possible for recruits.

Colonies gain nest energy (NE) when agents return to the nest and “unload” the energy gained at food sources. We estimated the energy content of a crop load of a *T. nylanderi* forager the following way: we measured foragers (N = 21, from 3 colonies) and estimated that full foragers carry ~0.15 mg of sugar solution. Given the energetic value of sucrose, we calculated that a forager feeding at a 1 molar sucrose solution collects ~0.75 Joule per foraging trip, whereas a forager feeding at a 0.1 molar solution would obtain ~0.075 J.

#### Process overview

The default simulation duration *t*_max_ was 5400 ticks (corresponding to 90 minutes), but we also tested a duration of 21600 ticks (corresponding to 6 h). Time and distance in the model were connected via the walking speeds (0.8 patches/tick outside the nest = *v*_outside_, 0.4 patches/tick in a t_andem_ run= *v*_tandem_), which were chosen to be similar to the walking velocity (in cm/sec) of *T. nylanderi* ants (Glaser and Grüter 2018).

When the model was initialized (*t*=0), the nest and either 2 or 10 food sources and the agents were created. In the situation without tandem running, only scouts were simulated. All agents started in the centre of the nest. Scouts immediately started to perform a random walk to search for food sources with the speed of *v*_outside_, whereas recruits patrolled inside the nest with speed *v*_nest_ (0.1 patches/tick) and waited to be recruited by another agent. All agents started with an energy of zero. When leaving the nest, this energy decreases every tick by a metabolic cost *M*_cost_ (see Table 1). *M*_cost_ was chosen so that the metabolic costs that accumulate during an average foraging trip correspond to ~0.1% of the value of energy obtained during a typical foraging trip (Fewell 1988). We estimated this by running several simulations and measuring foraging trip duration of our agents. We also ran simulations with metabolic rates that were 10-times higher or 10-times lower than our default value but found that this did not affect the general patterns (Fig. S1).

**Table 1:** Overview of the model parameters and the used values.

| Variables | Description | Default values | Other values tested |
| --- | --- | --- | --- |
| FS distance | Distance from nest centre to food source | 40 | 20 |
| FS number | Number of food sources | 2 or 10 |  |
| Colony size | Number of agents (foragers) in a colony | 100 | 20-200 |
| Scout-recruit ratio | The ratio of scouts and recruits in a colony | 1:4 (r = 0.2) or (all scouts) | 1:9 to 10:0 (r = 0.1 to 1.0) |
| FS<sub>High</sub> | Energy gained from the high-quality food source | 0.75 J |  |
| FS<sub>Low</sub> | Energy gained from the low-quality food source | 0.075 J |  |
| $t_{\max}$ | Duration of a simulation (1 tick ~ 1 second) | 5400 ticks | 21600 ticks |
| $v_{\text{outside}}$ | Walking velocity of ants outside the nest | 0.8 patch/ticks | |
| $v_{\text{nest}}$ | Walking velocity of ants inside the nest | 0.1 patch/ticks | |
| $v_{\text{tandem}}$ | Walking velocity of Tandem leader and Tandem follower towards the respective food source | 0.4 patch/ticks | |
| $M_{\text{cost}}$ | Metabolic or energy cost of walking outside | $2.446 \times e^{-7}$ J/tick | $2.446 \times e^{-6}$,<br>$2.446 \times e^{-8}$ |
| $t_{\text{scouts}}$ | Time a scout searches food before returning to nest | 600-900 ticks | |
| $t_{\text{nest-stay}}$ | Time a returned forager stays inside the nest | 60 ticks | |
| $t_{\text{tandemstarter}}$ | Time an active recruiter searches for a recruit inside the nest | 120 ticks | |
| $t_{\text{feeding}}$ | Feeding time of drinking agents | 120 ticks | |
| $p_{\text{break-up}}$ | Probability that tandems break up | 0/tick | 0.002/tick, |
|  |  |  | 0.005/tick |
| $p_{\text{recruitment}}$ | Probability to recruit when “satisfied” | 50% | |

When an agent finds a food source, it becomes a feeding agent and feeds for a duration of 120 ticks. It gains either 0.75 J or 0.075 J, depending on whether the food source is of high or low quality. If scouts do not find a food source within a certain time period (*t*_scouts_), they return to the nest. If they are at greater distances from the nest, unsuccessful scouts return quicker (600 ticks). Unsuccessful scouts that are closer to the nest (35 patches from the center) return if 900 ticks have passed. This was done to match observations that *T. nylanderi* scouts often return to their nest if they had been searching unsuccessfully for several minutes (S.M.G., personal observation). After their return, unsuccessful scouts wait idle inside the nest for 60 ticks (*t*_nest-stay_), before resuming to scout. At the end of the feeding time, agents return to the nest either as “satisfied” or “unsatisfied” foragers. Foragers that found a high-quality food source were always satisfied, whereas agents feeding at a low-quality food source had only a 10% probability to become satisfied. After unloading for the duration of *t*_nest-stay_, “satisfied” agents become prospective tandem leaders with a 50% probability (*p*recruitment), whereas unsatisfied agents would not recruit. This leads to a recruitment probability of 5-50% per trip, which is similar to what has been found in both *T. nylanderi* and *Pachycondyla harpax* (Glaser and Grüter 2018; Grüter et al. 2018). Satisfied agents return to the same food source they had visited before, either in a tandem run or alone. In other words, they use “route memory” to revisit a high-quality food source, but were unlikely to return to a low-quality food source (10% probability). Unsatisfied agents would not recruit and either wait inside the nest for a tandem leader (recruits) or they search for a new food source (scouts). Fig. S2 is a screenshot of a simulation showing the arrangement of the nest, food sources and some of the agent states.

Prospective tandem leaders stay inside the nest and search for a potential recruit for the duration of 120 ticks (*t*_tandemstarter_). A tandem run starts when a leader encounters a recruit on the same patch. By default, tandem runs do not break up but we also tested situations with a break-up probability of 0.002 and 0.005 per tick, which corresponds to tandem success rates of ~75 % and ~50% for the default distance (calculated based on an average tandem run duration of 127 ticks for the default food source distance). Lost tandem followers first perform a random walk for 180 ticks (*t*_search-time_) and – if they do not find a food source – have an equal probability to become either a scout or to return to the nest as an unsatisfied forager.

In the default settings, food sources were *ad libitum, i.e* they did not disappear during the simulations. Since this may not always be the case, we also simulated food sources that disappeared after they were visited by 10 agents to create a more dynamic foraging environment. If a food source disappears before ants return to it (either alone or in a tandem), agents reaching the old food source location search randomly for 180 ticks (*t*_search-time_), then they become unsatisfied foragers and return to the nest. If the food source vanishes during feeding, the agent becomes an unsatisfied forager. Food sources that have disappeared are replaced by an identical food source at the same position after 600 ticks have passed, which means that it has to be discovered again by scouts. For each simulation run, new inexperienced agents were created as described above.

#### Tested factors

- Spatio-temporal distribution and variability of food sources: we tested the effects of food source number (2 or 10), variability (only high-quality or variable quality. In the latter situation, food sources alternated in quality, *i.e*. FS 1 high-quality, FS2 low-quality, FS3, high-quality etc.), distance, foraging (simulation) duration and food source longevity (stable or short-lived).
- Scout-recruit ratio: next to our default ratio of 1:4, we tested several other ratios, including the extreme case with only scouts.
- Colony size: in addition to simulating a colony size of 100 agents, we tested a range of other colony sizes (Table 1).
- Food discovery time of scouts and recruits: For each simulation run, we quantified the time scouts needed to discover their first food source. In recruits, we measured both their waiting time inside the nest and the duration of the tandem run. These durations were averaged per forager type and per simulation run. Agents that did not discover a food source during an entire simulation were given the maximum value of 5400 ticks. Only conditions with successful tandem runs were considered.

The total nest energy NE (total J of all individual collection trips minus the total J of the metabolic costs) were measured for each simulation run.

#### Sensitivity of outcomes

Due to the stochasticity of simulations we performed 30 simulation runs for each tested combination of parameters. To assess how sensitive our model is to changes in the default parameters, we varied many of them and explored their effect, as mentioned above (see Table 1). We also simulated environments with only low-quality food sources. A pure scouting strategy was always better under these circumstances (Fig. S3). This is because tandem runs are very rare when all food sources are of low quality and recruits spend most of their time inside the nest.

#### Statistical analyses

All statistical analyses were performed using the software R 3.6.3 (www.r-project.org). Since different treatments occasionally had unequal variance (heteroscedasticity) or contained zeros and in order to provide a consistent statistical approach we used non-parametric statistical tests throughout. It should be noted, however, that when we compared parametric and non-parametric methods (Anova’s), they yielded very similar results. We used Mann-Whitney U tests to compare two independent samples and Wilcoxon signed-rank tests for paired data. In addition to the p-values, the R software provides the test statistic value W, which is a linear transformation of the usual rank sum statistic U. When three groups were compared, we used Kruskal-Wallis tests and Dunn tests with sequential Bonferroni corrections for post-hoc pair-wise comparisons (“FSA” package, Ogle et al. 2020) (Sokal and Rohlf 1995).

## Results

### Distribution of food sources

We first tested if the number of food sources and their distance from the nest affect the value of communication. When colonies had access to few food sources, they were more successful with tandem recruitment (scout-recruit ratio of 1:4) than colonies consisting only of scouts, irrespective of whether food sources were of high-quality (Fig. 2) (Mann-Whitney U Test, W = 215, p = 0.0004) or of variable quality (W = 307, p = 0.034). In a rich environment, with 10 food sources, colonies collected overall more energy (Fig. 2). Tandem communication was beneficial when food source quality was variable (W = 112, p < 0.0001), whereas colonies consisting only of scouts performed better when all 10 food sources were of high quality (W = 773, p < 0.0001). This general pattern did not change when food sources were closer to the nest (20 patches instead of 40 patches) (2 food sources, high-quality: p < 0.0001; variable-quality: W = 210, p = 0.0003; 10 food sources, high-quality: W = 827, p < 0.0001; variable-quality: W = 54, p < 0.0001), but colonies gained overall more energy when all food sources were close to the nest (Fig. 2). Fig. 2e and 2f illustrate the temporal development of nest energy during exemplary simulation runs that correspond to the conditions shown in Fig. 2a and 2b with high-quality food sources.

**Fig. 2.**
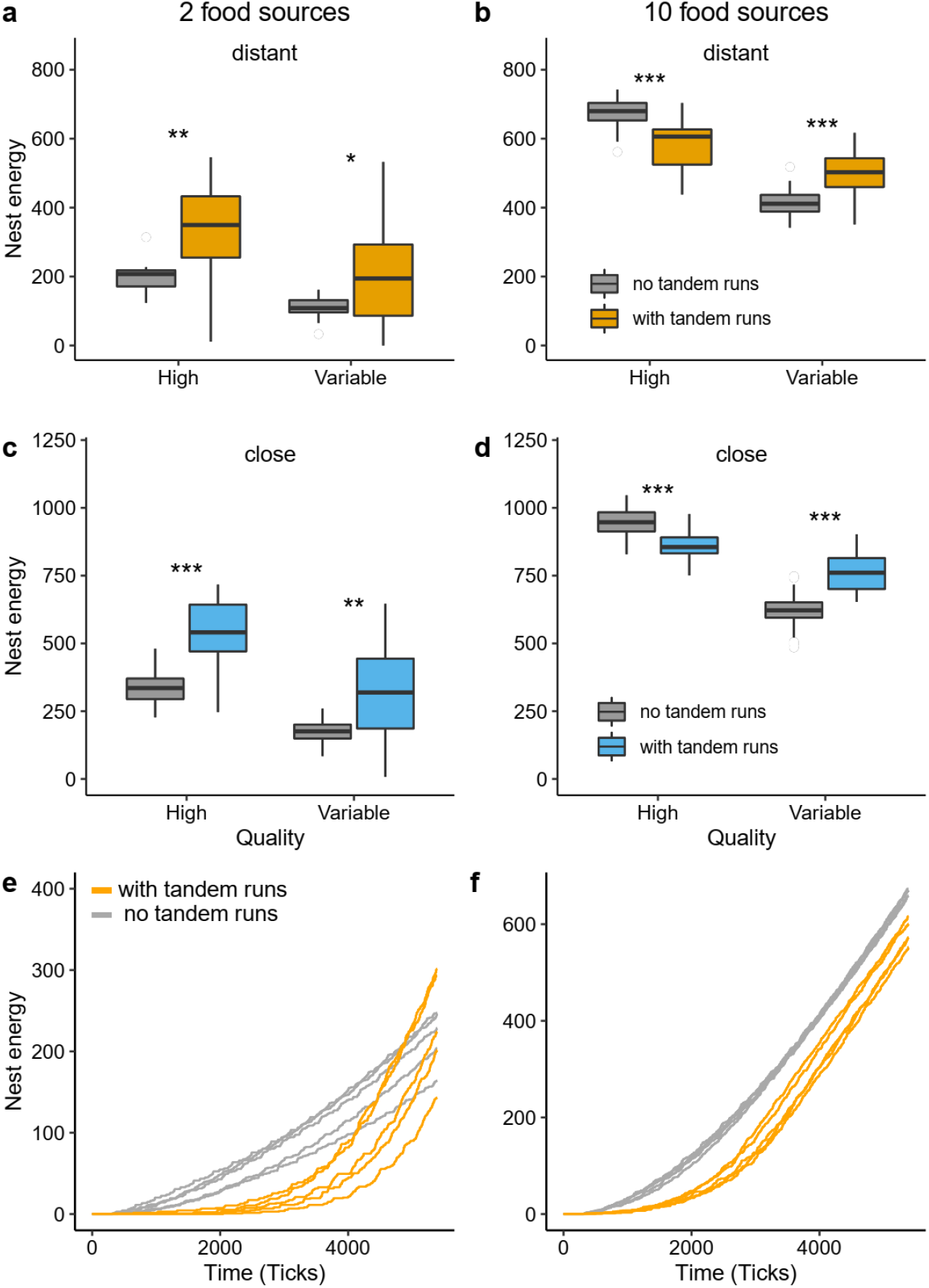
Nest energy with 2 or 10 food sources. In (a) and (b) food sources were at a distance of 40 patches, whereas in (c) and (d) food sources were at a distance of 20 patches. *p<0.05, **p<0.001, ***p<0.0001. In (e) and (f), nest energy is plotted over time for conditions as shown in (a) and (b) when all food sources were of high quality (5 simulation runs per treatment for visualisation of the trajectory).

### Foraging duration and food source longevity

When we increased the foraging duration (*i.e*. the simulation duration) from 5400 to 21600 ticks, we found a similar pattern. Tandem running was highly beneficial when there were few food sources (high-quality: W = 0, p < 0.0001; variable-quality: W = 0, p < 0.0001). Tandem runs were also beneficial when there were many food sources of variable quality (W = 56, p < 0.0001). In the case of many high-quality food sources, pure scout colonies performed better (W = 900, p < 0.0001). It is noteworthy that colonies with tandem communication were almost as successful in an environment with 2 food sources as in an environment with 10 food sources (Fig. 3a,b).

**Fig. 3.**
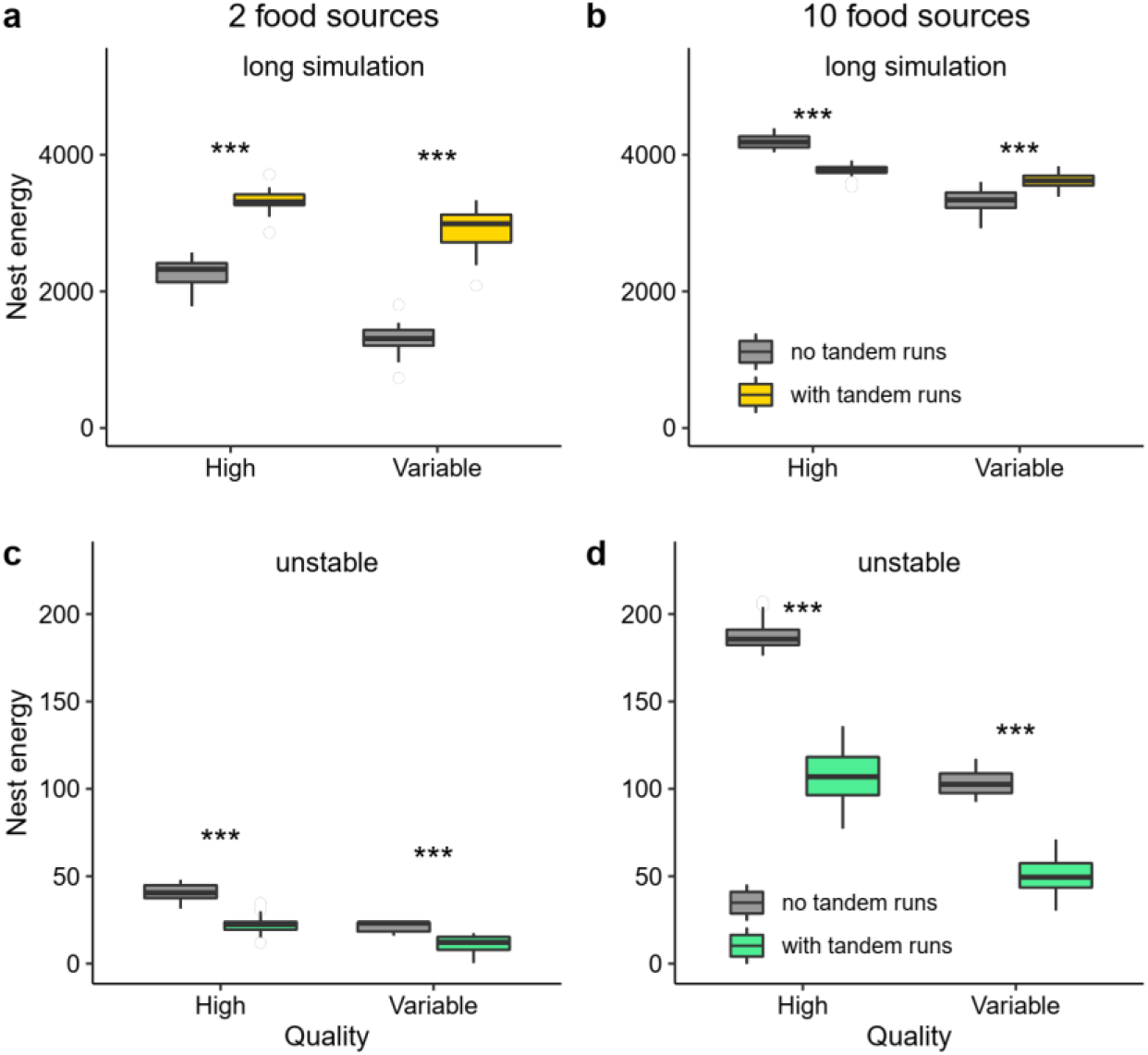
Nest energy with 2 or 10 food sources. In (a) and (b), food sources simulations were 4-times longer (21600 ticks instead of 5400). In (c) and (d), simulations lasted 5400 ticks and food sources disappeared if they were visited by 10 ants. A new food source appeared after a delay.

So far, we assumed that food sources offered food during the entire simulation. Next, we tested the effects of short-lived food sources. If food sources were unstable (sometimes called non-renewable), a scouting strategy was more successful, irrespective of the number of food sources and their variability (Fig. 3c,d) (2 food sources, high-quality: W = 897, p-value < 0.0001; variable-quality: W = 896, p-value < 0.0001; 10 food sources, high-quality: W = 900, p-value < 0.0001; variable-quality: W = 900, p-value < 0.0001). Differences were particularly pronounced when colonies were offered many food sources. Scouting remained the better strategy when we increased the foraging duration to 21600 ticks (e.g. 2 food sources, high-quality: W = 900, p-value < 0.0001; variable-quality: W = 900, p-value < 0.0001).

### Tandem success rate

Tandems do occasionally break up and we tested how this affects the energy collected by colonies. We compared colonies with 100% (default), ~75% and ~50% successful tandem runs and colonies with only scouts in an environment with few food sources, *i.e*. under conditions where tandem runs are beneficial (Fig. 2a). Our simulations show that a reduction in tandem success rate has a negative impact on the energy intake that is collected by colonies (Fig. 4). If only about 50% of the tandem runs are successful, colonies without any tandem running collect more energy in an environment with few, stable food sources (Fig. 4) (high-quality, 50% success rate vs. no tandems: W = 246, p = 0.002, variable-quality, 50% success rate vs. no tandems: W = 257, p = 0.004).

**Fig. 4.**
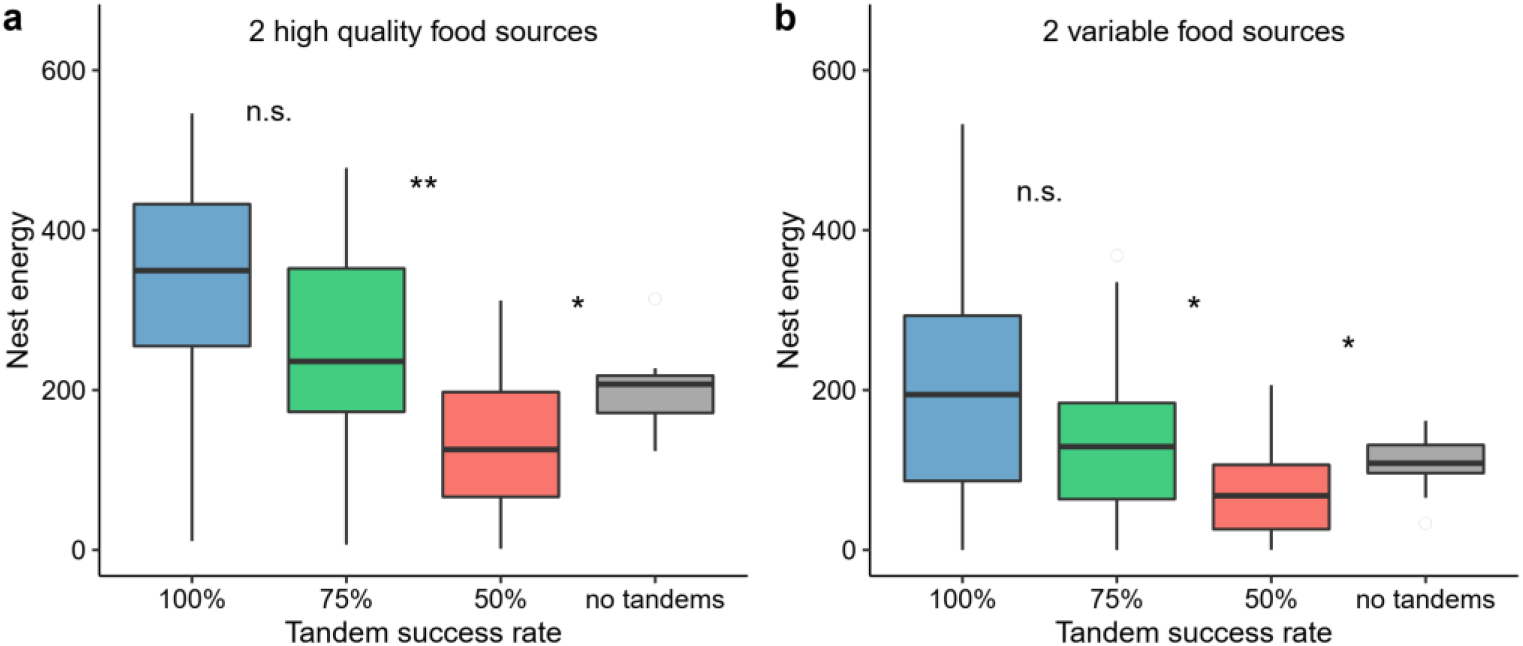
Nest energy with 2 food sources of high (a) and mixed (b) quality in relation to the tandem success rate. Adjacent treatment groups were compared, as indicated by asterisks or “n.s.”. No tandems = only scouts. Default settings were used for the other parameters.

However, tandem runs with a high rate of failure (50%) are not always a disadvantage compared to having no communication. When colonies can forage for longer (simulations of 21600 ticks), colonies that perform tandem runs with a ~50% break-up rate are more successful than colonies consisting of only scouts (Fig. 5) (high-quality, 50% success rate vs. no tandems: W = 866 p < 0.0001, variable-quality, 50% success rate vs. no tandems: W = 689, p = 0.0003), highlighting the benefits of imperfect communication over longer time periods.

**Fig. 5.**
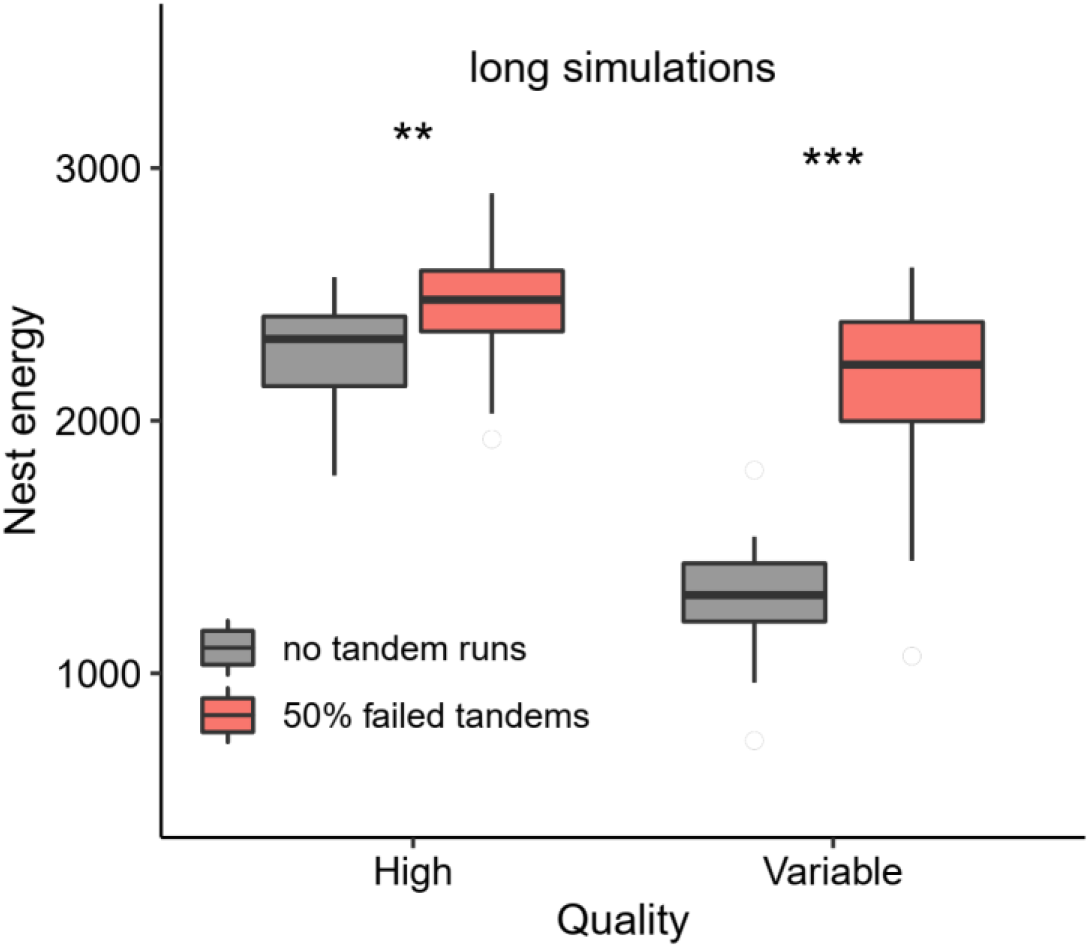
Nest energy with 2 food sources of high or variable quality and a long foraging duration. Colonies were scouting or could recruit with tandem runs that had a ~50% failure rate. Default settings were used for the other parameters.

### Colony size and scout-recruit ratio

We tested various colony sizes ranging from 20 to 200 agents in an environment with few, variable food sources, *i.e*. an environment that favours tandem running under default conditions (see Fig. 2a).

Colony size had a strong effect on the total collected energy that was collected (Fig. 6). If colonies were very small (20 foragers), they were least successful if they performed tandem runs and had a default scout-recruit ratio of 0.2 (Table 2). There was no difference in foraging success when colony size ranged from 30 to 50 foragers. However, colonies with tandem recruitment were more successful if they had at least 60 agents (Table 2). The most successful colonies contained 40% scouts, suggesting that the scout-recruit ratio has a considerable impact on colony success. Fig. 6b shows the nest energy collected per agent (nest energy/colony size). In colonies with only scouts, individual agents collected a relatively constant amount of energy irrespective of colony size (Spearman rank correlation: rho = 0.1, p = 0.09). In colonies with tandem running, on the other hand, individual agents collected more energy on average as colony size increased from 20 to 100 agents (r = 0.2, rho = 0.34, p < 0.0001; r = 0.4, rho = 0.35, p < 0.0001).

**Fig. 6.**
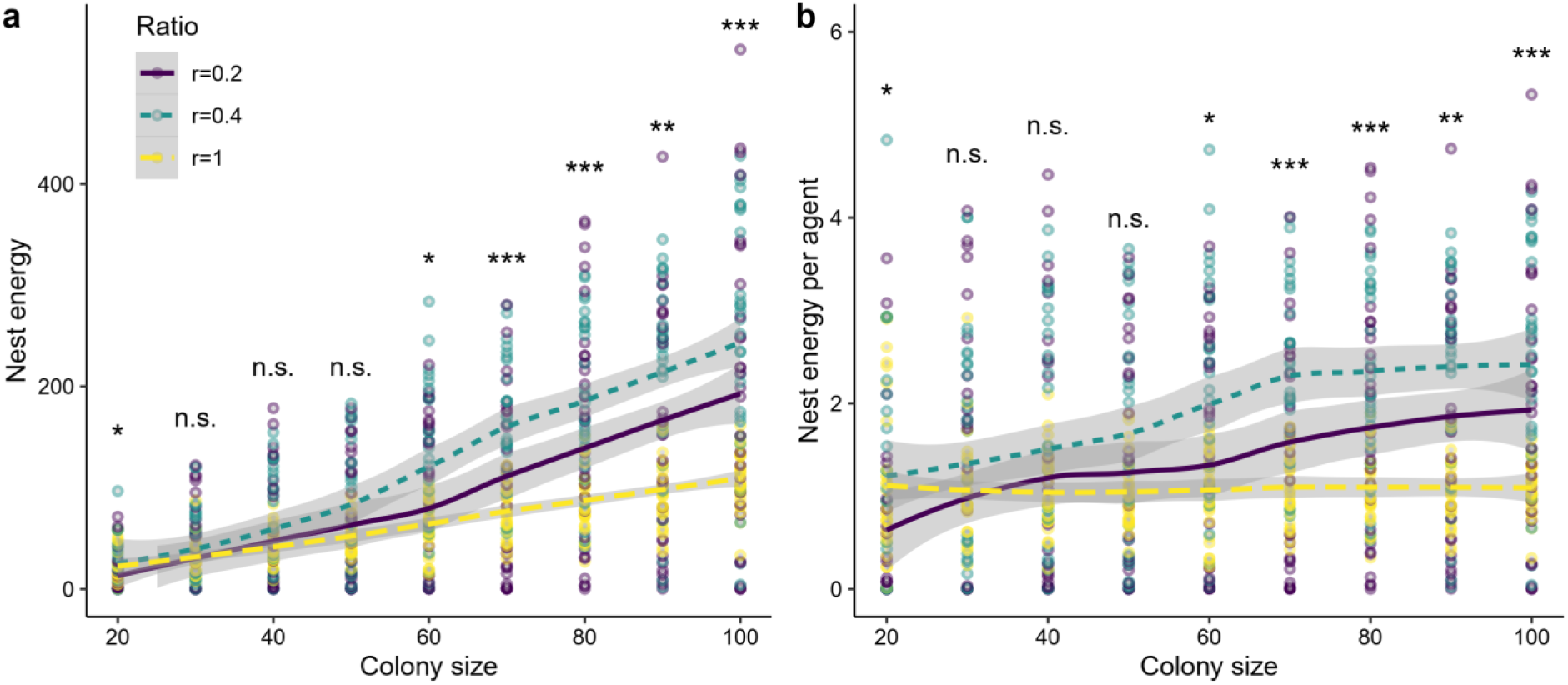
The relationship between colony size and nest energy (a) and nest energy per agent (b) in colonies with and without tandem runs in environments with two food sources of variable quality. Three scout-recruit ratios were simulated, r = 0.2 and r = 0.4 and colonies consisting only of scouts, r = 1.0. Grey area indicates confidence intervals. Significance tests refer to comparisons among ratios, separately for each colony size. P-values for total nest energy (a) or energy per agent (b) are identical. Default settings were used for the other parameters.

**Table 2:** Effect of colony size on nest energy. Three conditions were tested: in two conditions, colonies performed tandem runs and had a scout-recruit ratio of 0.2 or 0.4. In the third condition, colonies consisted only of scouts (1.0). Pair-wise comparisons were performed if the overall p < 0.05 and p-values were corrected using sequential Bonferroni.

| Colony size | Kruskal-Wallis Test |  | p-value of pair-wise comparisons |  |  |
| --- | --- | --- | --- | --- | --- |
| | $\chi^2$ | p-value | 0.2 vs. 0.4 | 0.2 vs. 1.0 | 0.4 vs. 1.0 |
| 20 | 12.7 | 0.002 | 0.018 | 0.002 | 0.42 |
| 30 | 2.33 | 0.31 | NA | NA | NA |
| 40 | 3.92 | 0.14 | NA | NA | NA |
| 50 | 1.81 | 0.41 | NA | NA | NA |
| 60 | 8.98 | 0.01 | 0.077 | 0.4 | 0.01 |
| 70 | 22.85 | <0.0001 | 0.0007 | 0.35 | <0.0001 |
| 80 | 20.6 | <0.0001 | 0.13 | 0.007 | <0.0001 |
| 90 | 15.86 | 0.0004 | 0.09 | 0.048 | 0.0002 |
| 100 | 19.17 | <0.0001 | 0.053 | 0.027 | <0.0001 |
| 200 | 46.59 | <0.0001 | 0.006 | 0.0001 | <0.0001 |

To explore this further, we simulated different scout-recruit ratios and different colony sizes to test how the balance between scouts and recruits affects colony foraging success. Simulations suggest that the optimal proportion of scouts is ~40% for the simulated environment, irrespective of colony size (Fig. 7). Interestingly, deviations from the optimal ratio have a larger negative impact in larger colonies (see “pointiness” of curves in Fig. 7). For example, there is no difference in success when colonies with 50 agents contain 40% or 80% of scouts (W = 119; p = 0.54). When colony size is 200, however, colonies with 80% scouts collect 31.5% less energy than colonies with 40% scouts (W = 199, p = 0.0001).

**Fig. 7.**
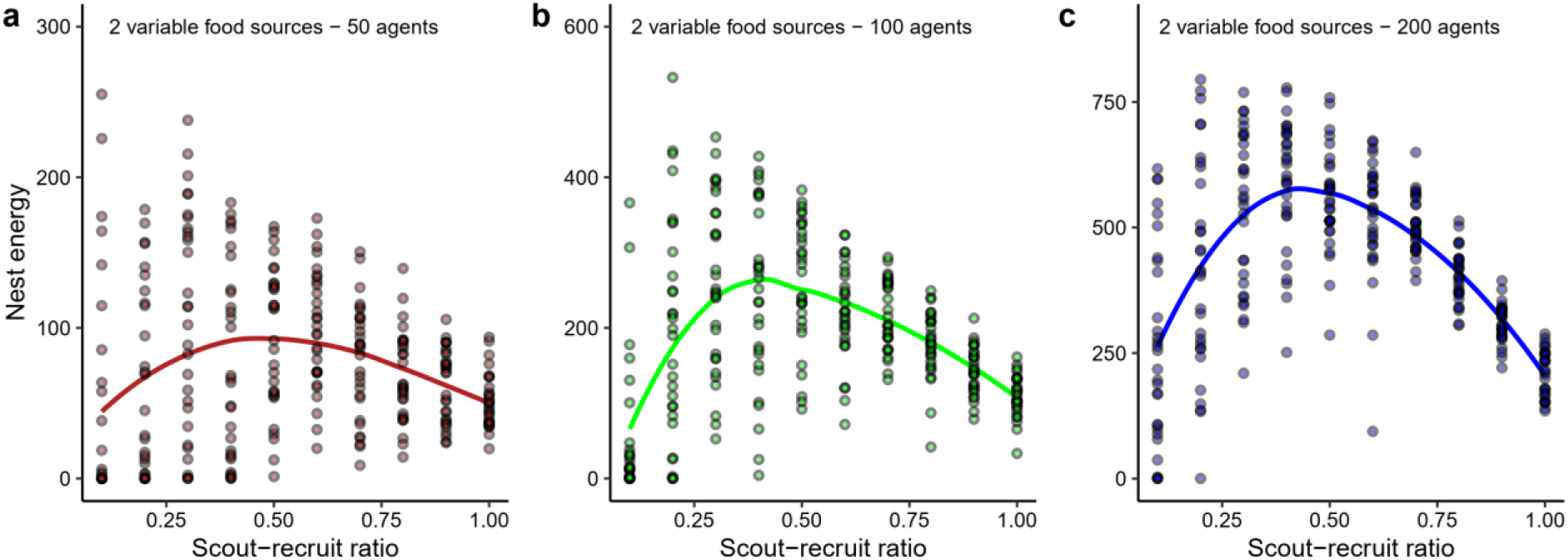
The effect of the scout-recruit ratio with three different colony sizes. The line shows the best fit line based on local polynomial regression using the LOESS method (locally estimated scatterplot smoothing). The smallest ratio was 0.1. A ratio of 1.0 refers to colonies containing only scouts.

### Discovery times

Unsurprisingly, foragers needed more time to find their first food source in an environment with few food sources compared to when there were many food sources (Fig. 8). Recruits needed less time in an environment with few, high-quality food sources compared to scouts (Wilcoxon-signed rank test: W = 143, p-value < 0.0001), whereas there was no difference when food sources were variable in quality (Fig. 8a) (W = 348, p-value = 0.13). However, in an environment with many food sources, scouts did comparatively better and needed a similar amount of time to locate their first food source when food sources were all high-quality (W = 327, p-value = 0.07). With many, variable food sources, scouts were significantly faster than recruits (Fig. 8b) (W = 720, p-value < 0.0001).

**Fig. 8.**
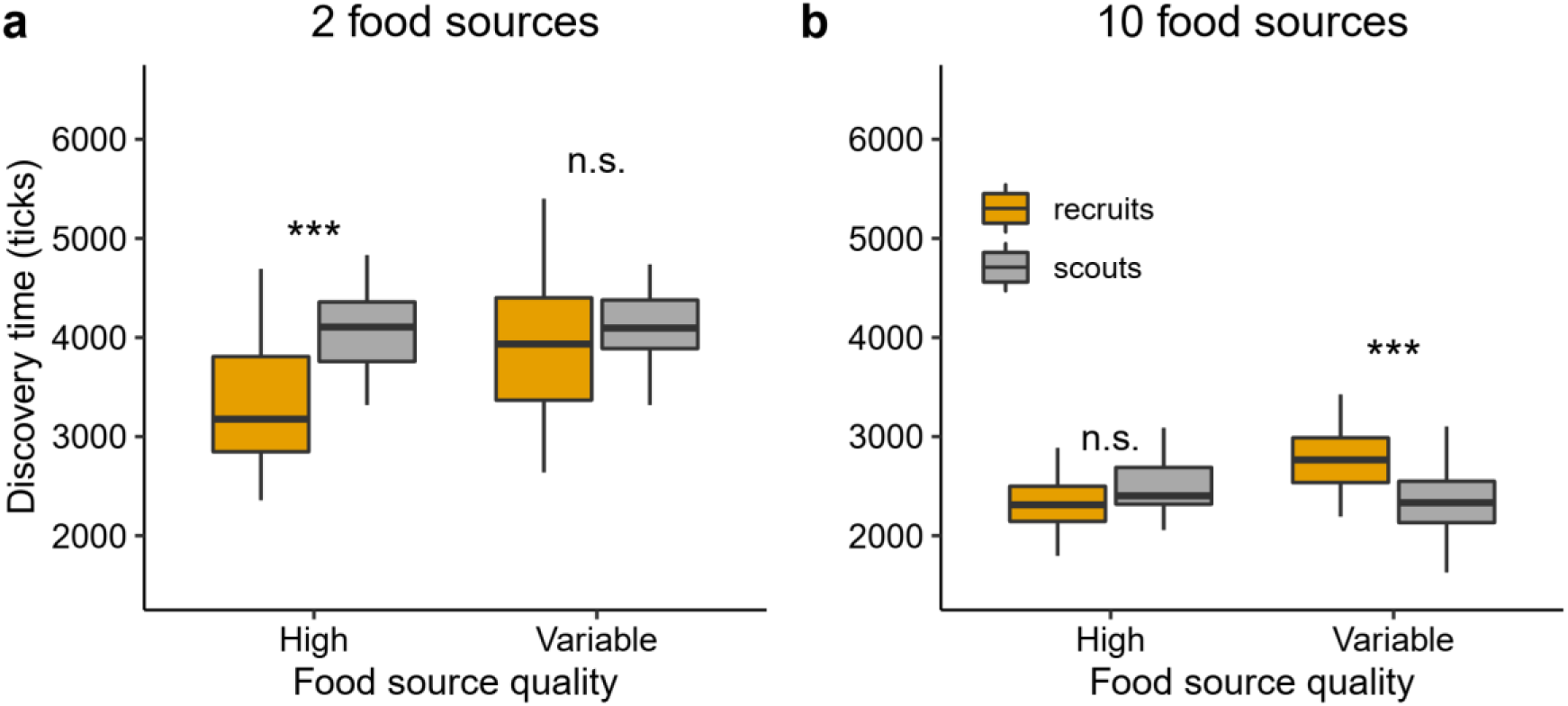
Time until agents located their first food source in environments with few (a) or many (b) food sources of constant or variable quality. For recruits, the food discovery time consisted of the time waiting inside the nest and the tandem run duration. Default settings were used for the other parameters.

## Discussion

Our simulations show that the spatio-temporal distribution of food sources greatly affects whether colonies with communication are more successful than colonies that employ a scouting strategy. Tandem running was beneficial when colonies were in an environment with few food sources (+ 57-83% nest energy) and when food sources were of variable quality (Fig. 2a,b). Colonies without communication were more successful (~15%) in a rich environment that offered only high-quality food sources. This is in line with studies that simulated honeybee foraging and found that communicating food source locations by waggle dancing is most beneficial if food sources are hard to find and of variable quality (Dornhaus et al. 2006; Beekman and Lew 2008; Schürch and Grüter 2014; I’Anson Price et al. 2019). Under such circumstances, the probability that scouts find high-quality food sources on their own is low and communicating the location of a relatively small number of high-quality patches becomes advantageous. As food source variability decreases and the number of high-quality food sources increases, scouts become more successful. Even though colonies with communication also collect more energy in such an environment, the benefits of communication no longer offset the costs of recruits waiting for information inside the nest. This highlights that communication often has considerable time and opportunity costs (Seeley 1983; Seeley and Visscher 1988; Dechaume-Moncharmont et al. 2005; Schürch and Grüter 2014; I’Anson Price et al. 2019).

It has been hypothesised that recruitment communication is particularly beneficial in an ephemeral environment (Sherman and Visscher 2002; Dornhaus and Chittka 2004; Grüter and Ratnieks 2011), *i.e*. when food sources last only for short time-periods and, thus, need to be exploited quickly. Counterintuitively, a simulation model of honeybee foraging has found that communication was less beneficial if food sources were shorter-lived (Schürch and Grüter 2014). Our simulations support their findings by showing that tandem running was a very successful strategy in a stable environment with relatively long foraging durations (*i.e*. with longer simulations) and few, variable food sources (Fig. 3a). A long-lasting food source could be a large insect (Lanan 2014), floral nectars or a group of honeydew secreting insects (Carroll and Janzen 1973; Quinet and Pasteels 1996; Völkl et al. 1999; Mailleux et al. 2003; Lanan 2014). A very different pattern was observed when resources were shorter-lived: colonies without communication were always more successful, irrespective of the foraging (simulation) duration (Fig. 3c,d). The most likely explanation is that colonies with communication pay time costs without being able to take advantage of the benefits of communication over longer time periods (see also Schürch and Grüter 2014). Our model differs from theirs in that our food sources only disappeared if they were exploited, rather than with a constant probability. A food source that disappears after it has been exploited could be a droplet of honeydew that fell on vegetation. Honeydew droplets on leaf surfaces represent an important food source for the tandem recruiting *Temnothorax curvispinosus* (Lynch et al. 1988).

Tandem runs occasionally break-up and success rates of ~50% to 90% are not uncommon (Wilson 1959; Pratt 2008; Kaur et al. 2017; Glaser and Grüter 2018; Grüter et al. 2018). We simulated different success rates and found that colonies with more successful tandem runs collected more energy (Fig 4). If the success rate was about 50%, colonies consisting only of scouts collected more energy in an environment with few food sources, *i.e*. a virtual environment that normally favours tandem running. When foraging durations were longer, on the other hand, colonies with tandem runs gained the upper hand over scouting colonies even though half of all tandem runs failed (Fig. 5). Under these circumstances, even a relatively low number of successful recruitment events can be very important because the discovered high-quality food sources can be exploited for longer time periods by successful recruits. Additionally, tandem recruitment can lead to an exponential increase of ants at a feeder even if a leader recruits <1 follower per trip. With exponential growth, the impact of communication will increase over time (Fig. 2e).

We found that colony size had a considerable effect on the value of tandem communication (Fig. 6). This contrasts with models of honeybee communication, where colony size did not greatly affect the benefits of communication (Dornhaus et al. 2006; Schürch and Grüter 2014), but is consistent with an empirical study on honeybee colony foraging success (Donaldson-Matasci et al. 2013) and a mathematical model of ant communication (Planqué et al. 2010). If colonies contained 60 or more foragers, tandem communication was usually beneficial. However, a pure scouting strategy was equally or more successful when colonies had 20 to 50 foragers, even in environments with few and variable food sources, *i.e*. a virtual environment that normally favours tandem running. This number of foragers could be expected in ant colonies with ~80-250 workers (assuming that foragers make up 20-30% of the worker population, e.g. Shaffer et al. 2013), which is also the typical colony size of many ant species that use tandem running and species with solitary foraging (Beckers et al. 1989). Our simulation results could explain why some species, e.g. in the genera *Diacamma* or *Neoponera*, do not perform tandem runs during foraging even though they use this recruitment method during migrations (Hölldobler 1984; Traniello and Hölldobler 1984; Maschwitz et a. 1986). Whether colonies employ tandem running might depend on the food sources they collect (e.g. small or large items) and whether they are risk-averse or risk-prone because tandem recruitment was often associated with a more unpredictable outcome in our simulations (greater variation in nest energy gain among simulations of a particular situation, see Fig. 2). A better understanding of the natural history of these species and similar species that do perform tandem runs (e.g. *Neoponera* vs. *Pachycondyla*) is needed to understand why some species use communication, while others forage solitarily.

Colony foraging performance depended on the proportions of scouts and recruits (Fig. 7). In our simulations with few food sources, colonies were most successful if scouts represented about 40% of the forager population, but this is likely to depend on the number and variability of food sources (see Fig. 2). Interestingly, having the right scout-recruit ratio is more important in larger colonies than in smaller ones, possibly because the foraging success of smaller colonies depends more on chance events, such as the discovery of a high-quality food source by a single scout. This suggests that larger colonies would benefit from having the ability to assess their current environment and adjust their use of communication accordingly. Whether this is indeed common is not well known, but it has recently been reported that honeybees are able to assess the value of communication and reduce their reliance on waggle dances if dance information is not beneficial in the current environment (I’Anson Price et al. 2019).

In the simulations, we measured the time recruits and scouts need to locate their first food source in environments with many or few food sources. We found that the food discovery time depends strongly on the environment. Recruits were faster in environments with few high-quality food sources, whereas scouts found a food source sooner in an environment with many, variable food sources. Our measurements also included the time that recruits wait inside the nest to find a tandem leader. Franks and Richardson (2006) found that tandem followers found a food source faster in their experiment with one food source, which, in combination with their other findings, indicated that tandem running fulfils the criteria for animal teaching set out by Caro and Hauser (1992; namely, a teacher [*i*] modifies its behaviour in the presence of a naive observer, [*ii*] at some cost to the leader [*iii*] so that the observer can learn more quickly or efficiently). Our simulations suggest that this is the case only in certain environments, namely those with few, high-quality resources. In other situations, scouts are likely to learn food source locations quicker and tandem running might no longer fulfil the criteria for animal teaching (namely that a follower acquires knowledge or learns a skill more rapidly or efficiently than it might otherwise do, or that it would not learn at all, see Caro and Hauser 1992).

Taken together, our simulations show that the value of tandem communication is highly dependent on the environment and the size and composition of the colony. Future studies should explore whether and how foragers can assess their foraging environment and modify their communication behaviour (see also Grüter and Czaczkes 2019). It would also be desirable to test the conclusions from our simulations empirically, but so far it has been challenging to stop ants from performing tandem runs without affecting their behaviour.

## Supporting information

Netlogo 1_10FS_no tandems

Netlogo 2_10FS_tandems

Netlogo3_2FS_no tandems

Netlogo 4_2FS_tandems

## Acknowledgments

We would like to thank the AG Foitzik for discussion and feedback on the model. S.M.G. was funded by the German Research Foundation (DFG: GR 4986/1-1).

---

**Fig. S1.**
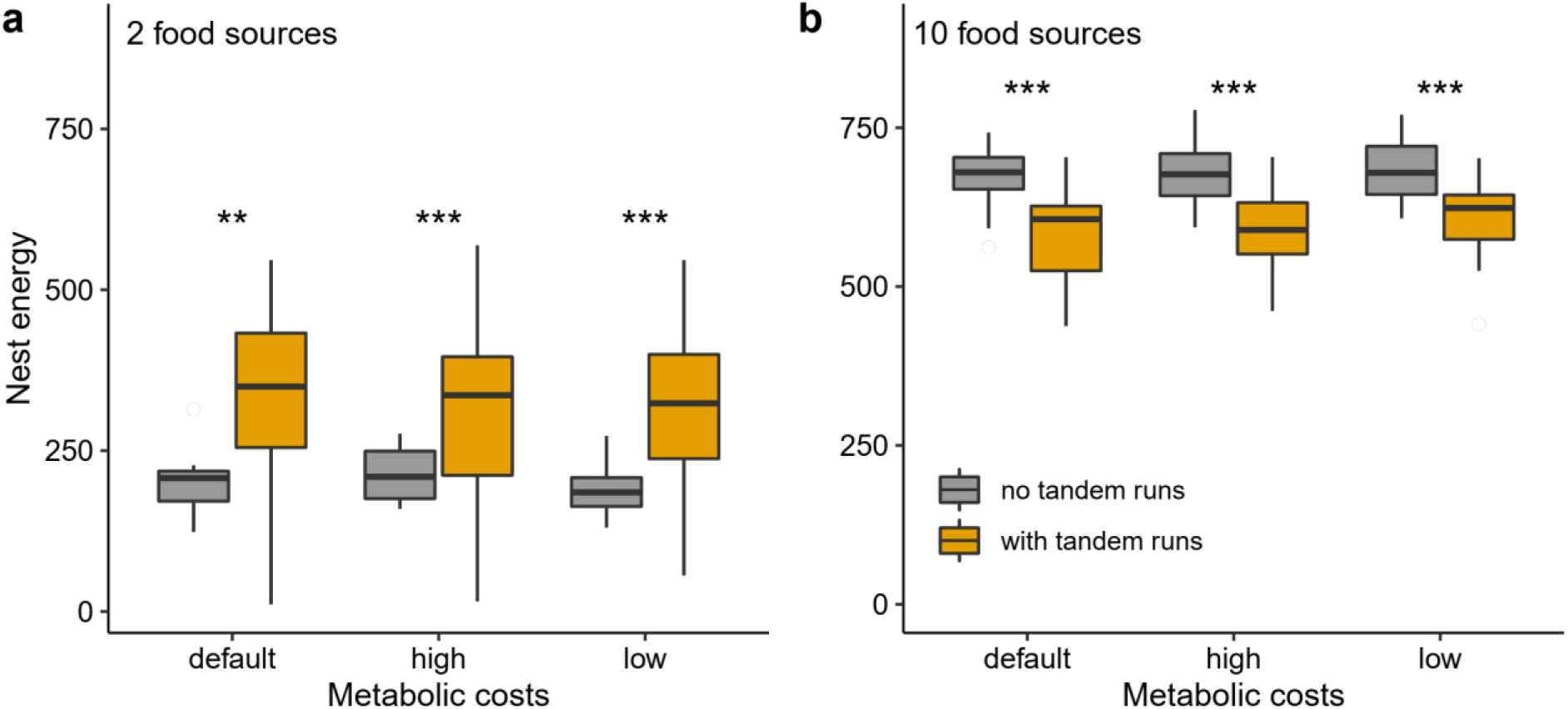
The effects of 10-times higher and 10-times lower metabolic costs on nest energy (see Table 1). Two (a) and ten (b) high-quality food sources were offered, default values were used for all other parameters. The default conditions match those shown in Fig. 2a and 2b. Mann-Whitney U tests, **p<0.001, ***p<0.0001.

**Fig. S2.**
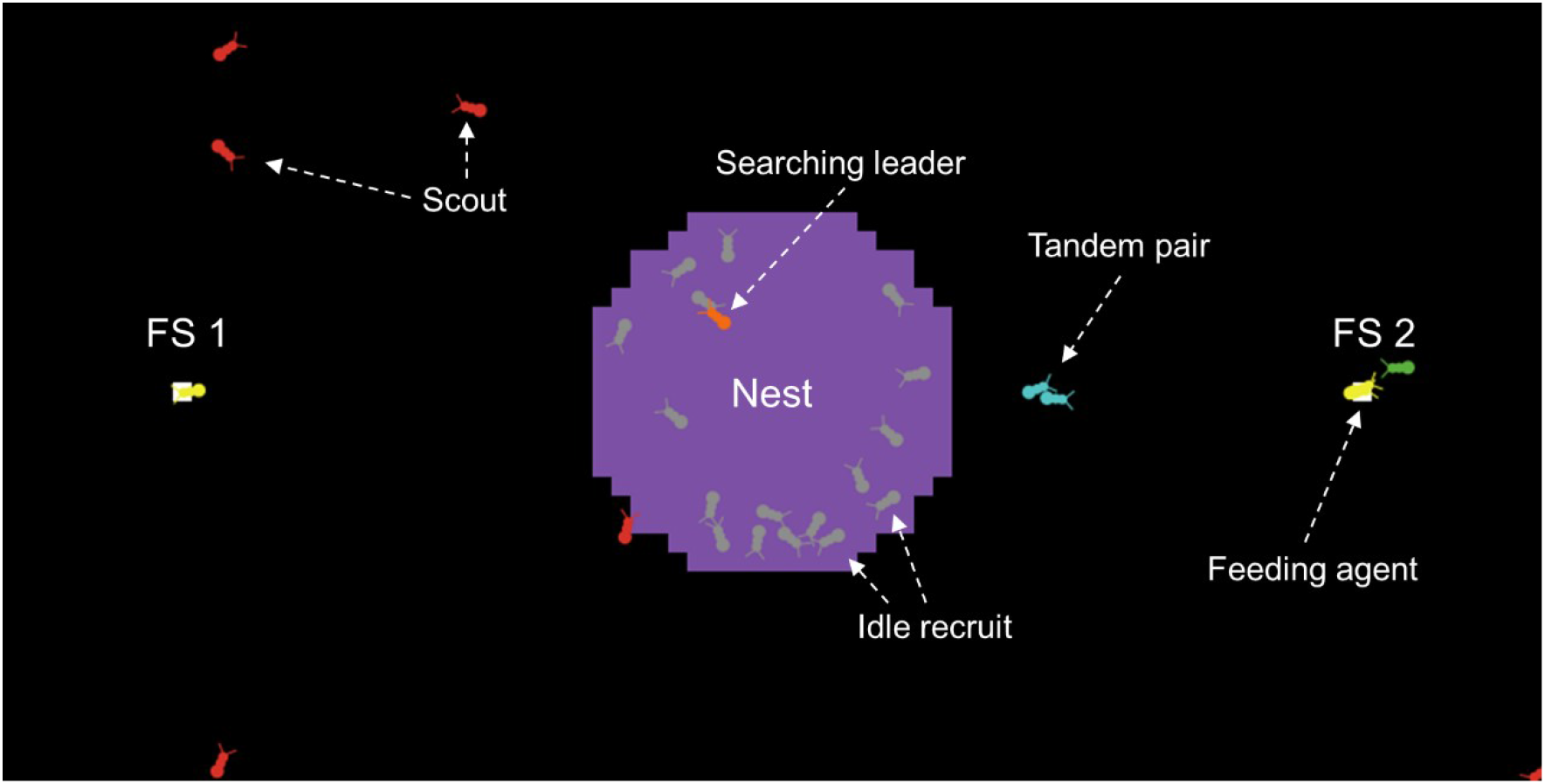
NetLogo interface showing some of the different agent types in different colours. In this situation, two food sources (FS 1 and FS 2) were offered.

**Fig. S3.**
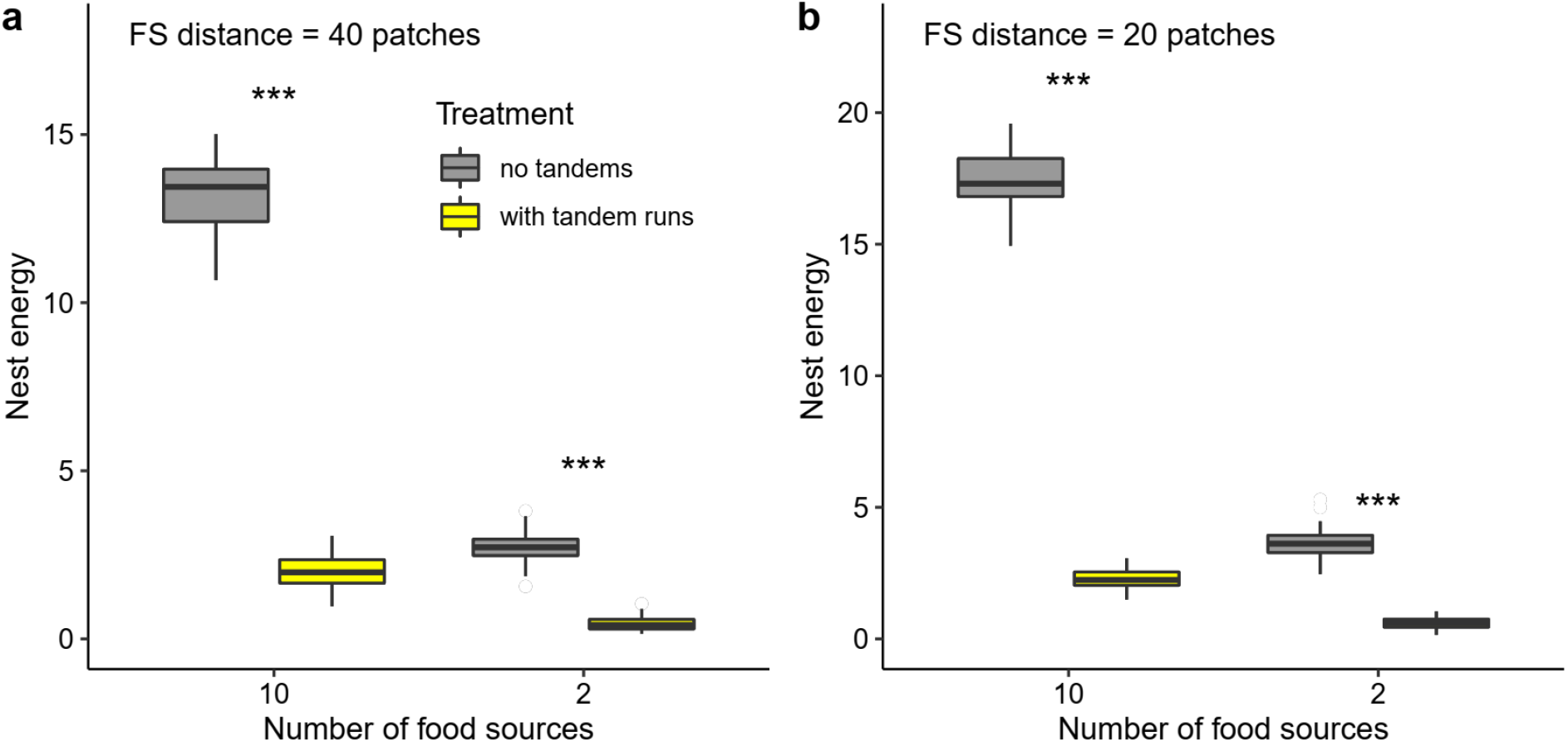
Nest energy of colonies with or without tandem runs when all food sources are of low quality. Two food source distances were simulated, 40 patches (a) or 20 patches (b). Default values were used for all other parameters. Mann-Whitney U tests, ***p<0.0001.

