## Supplementary material for "The adaptive value of tandem communication in ants: insights from an agent-based model": Netlogo 2_10FS_tandems

globals

[ nest-energy ]                                                ;; variable for measuring the colonys overall foraging success

breed [recruits recruit]                                       ;; waiting for Tandemstarters to begin Tandem
breed [scouts scout]                                           ;; outgoing ants
breed [waiting-scouts waiting-scout ]                          ;; unsuccessful scout waiting at the nest for going out again
breed [returning-scouts returning-scout]                       ;; scouts returning to the nest after being to far to long out
breed [drinking-ants drinking-ant ]                            ;; all ants at food-sources, drinking
breed [unsatisfied-foragers unsatisfied-forager]               ;; foragers that did not complete feeding and breeds (-50% of lost-TRS) after search time for food source is over

breed [happy-successful-foragers-1 happy-successful-forager-1 ];; happy foragers returning to nest from food source 1
breed [happy-successful-foragers-2 happy-successful-forager-2 ];; happy foragers returning to nest from food source 2
breed [happy-successful-foragers-3 happy-successful-forager-3 ];; happy foragers returning to nest from food source 3
breed [happy-successful-foragers-4 happy-successful-forager-4 ];; happy foragers returning to nest from food source 4
breed [happy-successful-foragers-5 happy-successful-forager-5 ];; happy foragers returning to nest from food source 5
breed [happy-successful-foragers-6 happy-successful-forager-6 ];; happy foragers returning to nest from food source 6
breed [happy-successful-foragers-7 happy-successful-forager-7 ];; happy foragers returning to nest from food source 7
breed [happy-successful-foragers-8 happy-successful-forager-8 ];; happy foragers returning to nest from food source 8
breed [happy-successful-foragers-9 happy-successful-forager-9 ];; happy foragers returning to nest from food source 9
breed [happy-successful-foragers-10 happy-successful-forager-10 ];; happy foragers returning to nest from food source 10


breed [successful-foragers-1 successful-forager-1]             ;; successful foragers satisfied with food source 1, memory to return to it
breed [successful-foragers-2 successful-forager-2]             ;; successful foragers satisfied with food source 2, memory to return to it
breed [successful-foragers-3 successful-forager-3]             ;; successful foragers satisfied with food source 3, memory to return to it
breed [successful-foragers-4 successful-forager-4]             ;; successful foragers satisfied with food source 4, memory to return to it
breed [successful-foragers-5 successful-forager-5]             ;; successful foragers satisfied with food source 5, memory to return to it
breed [successful-foragers-6 successful-forager-6]             ;; successful foragers satisfied with food source 6, memory to return to it
breed [successful-foragers-7 successful-forager-7]             ;; successful foragers satisfied with food source 7, memory to return to it
breed [successful-foragers-8 successful-forager-8]             ;; successful foragers satisfied with food source 8, memory to return to it
breed [successful-foragers-9 successful-forager-9]             ;; successful foragers satisfied with food source 9, memory to return to it
breed [successful-foragers-10 successful-forager-10]           ;; successful foragers satisfied with food source 10, memory to return to it

breed [Tandemstarters-1 Tandemstarter-1]                       ;; potential Tandemstarters satisfied with food source 1, memory to return to it
breed [Tandemstarters-2 Tandemstarter-2]                       ;; potential Tandemstarters satisfied with food source 2, memory to return to it
breed [Tandemstarters-3 Tandemstarter-3]                       ;; potential Tandemstarters satisfied with food source 3, memory to return to it
breed [Tandemstarters-4 Tandemstarter-4]                       ;; potential Tandemstarters satisfied with food source 4, memory to return to it
breed [Tandemstarters-5 Tandemstarter-5]                       ;; potential Tandemstarters satisfied with food source 5, memory to return to it
breed [Tandemstarters-6 Tandemstarter-6]                       ;; potential Tandemstarters satisfied with food source 6, memory to return to it
breed [Tandemstarters-7 Tandemstarter-7]                       ;; potential Tandemstarters satisfied with food source 7, memory to return to it
breed [Tandemstarters-8 Tandemstarter-8]                       ;; potential Tandemstarters satisfied with food source 8, memory to return to it
breed [Tandemstarters-9 Tandemstarter-9]                       ;; potential Tandemstarters satisfied with food source 9, memory to return to it
breed [Tandemstarters-10 Tandemstarter-10]                     ;; potential Tandemstarters satisfied with food source 10, memory to return to it

breed [Tandemleaders-1 Tandemleader-1]                         ;; Tandemleader to food source 1
breed [Tandemleaders-2 Tandemleader-2]                         ;; Tandemleader to food source 2
breed [Tandemleaders-3 Tandemleader-3]                         ;; Tandemleader to food source 3
breed [Tandemleaders-4 Tandemleader-4]                         ;; Tandemleader to food source 4
breed [Tandemleaders-5 Tandemleader-5]                         ;; Tandemleader to food source 5
breed [Tandemleaders-6 Tandemleader-6]                         ;; Tandemleader to food source 6
breed [Tandemleaders-7 Tandemleader-7]                         ;; Tandemleader to food source 7
breed [Tandemleaders-8 Tandemleader-8]                         ;; Tandemleader to food source 8
breed [Tandemleaders-9 Tandemleader-9]                         ;; Tandemleader to food source 9
breed [Tandemleaders-10 Tandemleader-10]                       ;; Tandemleader to food source 10

breed [Tandemfollowers-1 Tandemfollower-1]                     ;; Tandemfollower to food source 1
breed [Tandemfollowers-2 Tandemfollower-2]                     ;; Tandemfollower to food source 2
breed [Tandemfollowers-3 Tandemfollower-3]                     ;; Tandemfollower to food source 3
breed [Tandemfollowers-4 Tandemfollower-4]                     ;; Tandemfollower to food source 4
breed [Tandemfollowers-5 Tandemfollower-5]                     ;; Tandemfollower to food source 5
breed [Tandemfollowers-6 Tandemfollower-6]                     ;; Tandemfollower to food source 6
breed [Tandemfollowers-7 Tandemfollower-7]                     ;; Tandemfollower to food source 7
breed [Tandemfollowers-8 Tandemfollower-8]                     ;; Tandemfollower to food source 8
breed [Tandemfollowers-9 Tandemfollower-9]                     ;; Tandemfollower to food source 9
breed [Tandemfollowers-10 Tandemfollower-10]                   ;; Tandemfollower to food source 10

breed [lost-TRs lost-TR ]                                      ;; Tandemfollower that lost food source direction

turtles-own

[ feeding-time                                                 ;; time an ant feeds, default 120 ticks
  scouts-time                                                  ;; time of a scout for scouting, default 600 ticks
  nest-stay-time                                               ;; time an ant stays in the nest, 60 ticks for all breeds except Tandemstarters (120 ticks)
  search-time                                                  ;; time a successful forager searches for "her" foodsource. When over, then memory loss/give up and scouting again, default 120 ticks
  forager-timer                                                ;; timer for foragers (scouts, recruits and followers)
  forager-energy                                               ;; energy of a forager
  happiness ]                                                  ;; happiness of a forager

patches-own
[ nest?                                                        ;; true on nest patches, false elsewhere
 if-food?                                                      ;; true on FS patches, false elsewhere
 nest-scent                                                    ;; scent of nest, stronger when nearer to the nest
 FS-number                                                     ;; number of the food sources (1 to 10)
 FS-scent-1                                                    ;; scent of food source 1, number that is higher closer to it
 FS-scent-2                                                    ;; scent of food source 2, number that is higher closer to it
 FS-scent-3                                                    ;; scent of food source 3, number that is higher closer to it
 FS-scent-4                                                    ;; scent of food source 4, number that is higher closer to it
 FS-scent-5                                                    ;; scent of food source 5, number that is higher closer to it
 FS-scent-6                                                    ;; scent of food source 6, number that is higher closer to it
 FS-scent-7                                                    ;; scent of food source 7, number that is higher closer to it
 FS-scent-8                                                    ;; scent of food source 8, number that is higher closer to it
 FS-scent-9                                                    ;; scent of food source 9, number that is higher closer to it
 FS-scent-10                                                   ;; scent of food source 10, number that is higher closer to it

  FS-timer-1                                                   ;; time till the vanished food source 1 reappears. Adjustable via slider FS-timer, default 1/10 of experimenatl durance
  FS-timer-2                                                   ;; time till the vanished food source 2 reappears. Adjustable via slider FS-timer, default 1/10 of experimenatl durance  ;; time till FS-scent-2 runs out, adjustable via slider FS-timer, default 300 ticks
  FS-timer-3                                                   ;; time till the vanished food source 3 reappears. Adjustable via slider FS-timer, default 1/10 of experimenatl durance
  FS-timer-4                                                   ;; time till the vanished food source 4 reappears. Adjustable via slider FS-timer, default 1/10 of experimenatl durance
  FS-timer-5                                                   ;; time till the vanished food source 5 reappears. Adjustable via slider FS-timer, default 1/10 of experimenatl durance
  FS-timer-6                                                   ;; time till the vanished food source 6 reappears. Adjustable via slider FS-timer, default 1/10 of experimenatl durance
  FS-timer-7                                                   ;; time till the vanished food source 7 reappears. Adjustable via slider FS-timer, default 1/10 of experimenatl durance
  FS-timer-8                                                   ;; time till the vanished food source 8 reappears. Adjustable via slider FS-timer, default 1/10 of experimenatl durance
  FS-timer-9                                                   ;; time till the vanished food source 9 reappears. Adjustable via slider FS-timer, default 1/10 of experimenatl durance
  FS-timer-10                                                  ;; time till the vanished food source 10 reappears. Adjustable via slider FS-timer, default 1/10 of experimenatl durance

  FS-quality                                                   ;; quality of food sources: if reward = 0.75 it has a high quality (1), else it has low quality (0)
  FS-energy ]                                                  ;; energy of food source: if empty then the FS vanishes

;;;;;;;;;;;;;;;;;;;;;;;;
;;; Setup procedures ;;;
;;;;;;;;;;;;;;;;;;;;;;;;

to setup
  clear-all                                                     ;; (re)creation of an empty arena
  set-default-shape turtles "bug"                               ;; form of all agents shall be ants
  create-scouts number-of-scouts                                ;; an adjustable number of scouts will be created
  create-recruits number-of-recruits                            ;; an adjustable number of recruits will be created
ask turtles                                                     ;; all agents at the start of the simulation have the same
   [ set size 2                                                 ;; size
    set forager-energy 0                                        ;; 0 energy
    set feeding-time 120                                        ;; same feeding time
        set search-time FS-searching-time                       ;; same search time for a food source
    set nest-stay-time 60                                       ;; and their nest stay time is initially set for the (adjustable) unloading time
 ifelse breed = scouts                                          ;; if they are scouts,
        [ set color red ]                                       ;; they shall be red
        [ set color grey ] ]                                    ;; else they are recruits and are grey
  setup-patches                                                 ;; orders to create the world
  reset-ticks                                                   ;; reset tick counter
end

to setup-patches
ask patches
[ setup-nest                                                     ;; nest is created
  setup-food                                                     ;; food sources are created
  color-patch ]                                                  ;; nest and food sources are given color
end

to setup-nest                                                    ;; creation of nest
set nest? (distancexy 0 0) < nest-size                           ;; set nest?-variable: true inside the nest, false elsewhere
set nest-scent 200 - distancexy 0 0                              ;; spread a nest-scent over the whole arena which is stronger near the nest
set nest-energy 0                                                ;; initial nest energy is set to 0
end

to setup-food                                                    ;; setup of food sources with scent and quality

if (distancexy 0 51) < Food-size                                 ;; creation of a food source at 40 patches distance from the north side of the nest
[ set FS-number 1                                                ;; give it the number 1
   ifelse reward-FS-1 = 0.75                                     ;; if it is the food source 1 and the (adjustable) reward is high enough then
  [ set  FS-quality  1                                           ;; it has a high quality
    set FS-energy food-energy-high ]                             ;; and a high energy
  [ set  FS-quality  0                                           ;; it has a low quality
      set FS-energy food-energy-low ] ]                          ;; and low energy
  set FS-scent-1 200 - distancexy 0 51                           ;; scent of food source 1

if (distancexy 30 39 ) < Food-size                               ;; creation of food source 2 at 40 patches distance from the northeast of the nest
[ set FS-number 2                                                ;; give it the number 2
   ifelse reward-FS-2 = 0.75                                     ;; if it is the food source 2 and the (adjustable) reward is high enough then
  [ set  FS-quality  1                                           ;; it has a high quality
    set FS-energy food-energy-high ]                             ;; and a high energy
  [ set  FS-quality  0                                           ;; it has a low quality
      set FS-energy food-energy-low ] ]                          ;; and low energy
  set FS-scent-2 200 - distancexy 30 39                          ;; scent of food source 2


  if (distancexy 48 15) < Food-size                              ;; creation of a food source at 40 patches distance from the east of the nest
[ set FS-number 3                                                ;; give it the number 3
   ifelse reward-FS-3 = 0.75                                     ;; if it is the food source 3 and the (adjustable) reward is high enough then
  [ set  FS-quality  1                                           ;; it has a high quality
    set FS-energy food-energy-high ]                             ;; and a high energy
  [ set  FS-quality  0                                           ;; it has a low quality
      set FS-energy food-energy-low ] ]                          ;; and low energy
  set FS-scent-3 200 - distancexy 48 15                          ;; scent of food source 3

  if (distancexy 48 -17) < Food-size                             ;; creation of a food source at 40 patches distance from the east-south of the nest
[ set FS-number 4                                                ;; give it the number 4
   ifelse reward-FS-4 = 0.75                                     ;; if it is the food source 4 and the (adjustable) reward is high enough then
  [ set  FS-quality  1                                           ;; it has a high quality
    set FS-energy food-energy-high ]                             ;; and a high energy
  [ set  FS-quality  0                                           ;; it has a low quality
      set FS-energy food-energy-low ] ]                          ;; and low energy
  set FS-scent-4 200 - distancexy 48 -17                         ;; scent of food source 4

  if (distancexy 30 -39) < Food-size                             ;; creation of a food source at 40 patches distance from southeast of the nest
[ set FS-number 5                                                ;; give it the number 5
   ifelse reward-FS-5 = 0.75                                     ;; if it is the food source 5 and the (adjustable) reward is high enough then
  [ set  FS-quality  1                                           ;; it has a high quality
    set FS-energy food-energy-high ]                             ;; and a high energy
  [ set  FS-quality  0                                           ;; it has a low quality
      set FS-energy food-energy-low ] ]                          ;; and low energy
  set FS-scent-5 200 - distancexy 30 -39                         ;; scent of food source 5

  if (distancexy 0 -51) < Food-size                              ;; creation of a food source at 40 patches distance south of the nest
[ set FS-number 6                                                ;; give it the number 6
   ifelse reward-FS-6 = 0.75                                     ;; if it is the food source 6 and the (adjustable) reward is high enough then
  [ set  FS-quality  1                                           ;; it has a high quality
    set FS-energy food-energy-high ]                             ;; and a high energy
  [ set  FS-quality  0                                           ;; it has a low quality
      set FS-energy food-energy-low ] ]                          ;; and low energy
  set FS-scent-6 200 - distancexy 0 -51                          ;; scent of food source 6

  if (distancexy -30 -39) < Food-size                            ;; creation of a food source at 40 patches distance southwest of the nest
[ set FS-number 7                                                ;; give it the number 7
   ifelse reward-FS-7 = 0.75                                     ;; if it is the food source 7 and the (adjustable) reward is high enough then
  [ set  FS-quality  1                                           ;; it has a high quality
    set FS-energy food-energy-high ]                             ;; and a high energy
  [ set  FS-quality  0                                           ;; it has a low quality
      set FS-energy food-energy-low ] ]                          ;; and low energy
  set FS-scent-7 200 - distancexy -30 -39                        ;; scent of food source 7

  if (distancexy -48 -17) < Food-size                            ;; creation of a food source at 40 patches distance west-south of the nest
[ set FS-number 8                                                ;; give it the number 8
   ifelse reward-FS-8 = 0.75                                     ;; if it is the food source 8 and the (adjustable) reward is high enough then
  [ set  FS-quality  1                                           ;; it has a high quality
    set FS-energy food-energy-high ]                             ;; and a high energy
  [ set  FS-quality  0                                           ;; it has a low quality
      set FS-energy food-energy-low ] ]                          ;; and low energy
  set FS-scent-8 200 - distancexy -48 -17                        ;; scent of food source 8

  if (distancexy -48 15) < Food-size                             ;; creation of a food source at 40 patches distance west of the nest
[ set FS-number 9                                                ;; give it the number 9
   ifelse reward-FS-9 = 0.75                                     ;; if it is the food source 9 and the (adjustable) reward is high enough then
  [ set  FS-quality  1                                           ;; it has a high quality
    set FS-energy food-energy-high ]                             ;; and a high energy
  [ set  FS-quality  0                                           ;; it has a low quality
      set FS-energy food-energy-low ] ]                          ;; and low energy
  set FS-scent-9 200 - distancexy -48 15                         ;; scent of food source 9

  if (distancexy -30 39) < Food-size                             ;; creation of a food source at 40 patches distance northwest of the nest
[ set FS-number 10                                               ;; give it the number 10
   ifelse reward-FS-10 = 0.75                                    ;; if it is the food source 10 and the (adjustable) reward is high enough then
  [ set  FS-quality  1                                           ;; it has a high quality
    set FS-energy food-energy-high ]                             ;; and a high energy
  [ set  FS-quality  0                                           ;; it has a low quality
      set FS-energy food-energy-low ] ]                          ;; and low energy
  set FS-scent-10 200 - distancexy -30 39                        ;; scent of food source 10

ifelse FS-number > 0                                             ;; if it is a food source
  [ set if-food? true ]                                          ;; it has food to offer
  [ set if-food? false ]                                         ;; else: it has no food to offer and is just a regular patch
  set FS-timer-1  FS-timer                                       ;; set time till the vanished food source 1 reappears to the adjustable FS-timer
  set FS-timer-2  FS-timer                                       ;; set time till the vanished food source 2 reappears to the adjustable FS-timer
  set FS-timer-3  FS-timer                                       ;; set time till the vanished food source 3 reappears to the adjustable FS-timer
  set FS-timer-4  FS-timer                                       ;; set time till the vanished food source 4 reappears to the adjustable FS-timer
  set FS-timer-5  FS-timer                                       ;; set time till the vanished food source 5 reappears to the adjustable FS-timer
  set FS-timer-6  FS-timer                                       ;; set time till the vanished food source 6 reappears to the adjustable FS-timer
  set FS-timer-7  FS-timer                                       ;; set time till the vanished food source 7 reappears to the adjustable FS-timer
  set FS-timer-8  FS-timer                                       ;; set time till the vanished food source 8 reappears to the adjustable FS-timer
  set FS-timer-9  FS-timer                                       ;; set time till the vanished food source 9 reappears to the adjustable FS-timer
  set FS-timer-10  FS-timer                                      ;; set time till the vanished food source 10 reappears to the adjustable FS-timer
end

to color-patch                                                   ;; colour the world
ifelse nest?                                                     ;; nest is violet
[ set pcolor violet ]
[ if FS-number > 0                                               ;; if there are food sources, their color is
[ set pcolor white ] ]                                           ;; white
end

;;;;;;;;;;;;;;;;;;;;;;;;;;;;;;;;;
;;; START OF REPEATED ACTIONS ;;;
;;;;;;;;;;;;;;;;;;;;;;;;;;;;;;;;;

to go

  ask turtles
 [ if nest?
    [ set happiness 0 ] ]                                         ;; reset happiness for all agents to 0 (becoming ready for another foraging trip)

;;;;;;;;;;;;;;;;;;;;;;;;
;;; RECRUITS AT NEST ;;;
;;;;;;;;;;;;;;;;;;;;;;;;

ask recruits                                                      ;; recruits shall be
[  set color grey                                                 ;; grey
if nest?                                                          ;; in the nest
[  wiggle                                                         ;; randomly change direction inside the nest
   nest-stay                                                      ;; and walk slowly inside the nest
   set forager-timer forager-timer + 1 ] ]                        ;; Timer for time spend waiting in the nest advances

;;;;;;;;;;;;;;
;;; SCOUTS ;;;
;;;;;;;;;;;;;;

ask scouts
   [ set color red                                                ;; all scouts are of red colour
     scouting                                                     ;; general procedure of scouts
if if-food? = true                                                ;; if food is found
[ ask scouts-here                                                 ;; scouts at a FS
  [ output-show forager-timer                                     ;; trigger the timer to display their time spend scouting
    set breed drinking-ants                                       ;; become drinking-ants
    stop ] ]                                                      ;; do not "overrun" small food sources
        if scouts-time >= 600                                     ;; if they are to long out
        [ decision-of-foraging ] ]                                ;; face decision wether to go on scouting or return to nest
  ask returning-scouts                                            ;; returning scouts
    [ return-to-nest                                              ;; return to the nest
 if nest?                                                         ;; at the nest
[ set breed waiting-scouts   ] ]                                  ;; scouts from unsuccessful scouting wait at nest
  ask waiting-scouts                                              ;; scouts waiting at the nest
  [ nest-stay                                                     ;; and walk slowly inside the nest
    set nest-stay-time nest-stay-time - 1                         ;; time they stay in the nest counts down
ifelse nest-stay-time <= 0                                         ;; if their nest stay time is over
  [ set breed scouts                                              ;; become scouts again
      reset-times ]                                               ;; reset internal timers
[ nest-stay ] ]                                                   ;; if time is not over, stay at nest as waiting scouts

;;;;;;;;;;;;;;;;;;;;;;
;;; OTHER FORAGERS ;;;
;;;;;;;;;;;;;;;;;;;;;;

  ask drinking-ants                                               ;; ants at a FS
  [ set color yellow                                              ;; are yellow in color
    ifelse if-food? = true                                        ;; if there is food
    [ feed ]                                                      ;; feed on it
    [ set breed unsatisfied-foragers ] ]                          ;; become an unsatisfied forager

;;;;;;;;;;;;;;;;;;;;;;;;;;;;;;;;;
;;; HAPPY ANTS RETURN TO NEST ;;;
;;;;;;;;;;;;;;;;;;;;;;;;;;;;;;;;;

ask happy-successful-foragers-1                                   ;; ants happy with food source 1
[  set color green                                                ;; are green
    ifelse nest?                                                  ;; if there are at nest
  [  nest-stay                                                    ;; walk slowly
     set nest-stay-time nest-stay-time - 1 ]                      ;; time they stay in the nest counts down
   [ return-to-nest ]                                             ;; else they are outside and return to the nest
if nest? and nest-stay-time = 0                                   ;; if they are at the nest and their nest stay time is over
[ unload                                                          ;; they give their forager energy (positive or negative value) to the nest (changing nest energy)
  decision-of-recruitment-1 ] ]                                   ;; face decision whether to recruit to food source 1 or not

ask happy-successful-foragers-2                                   ;; ants happy with food source 2
[  set color green                                                ;; are green
    ifelse nest?                                                  ;; if there are at nest
  [  nest-stay                                                    ;; walk slowly
     set nest-stay-time nest-stay-time - 1 ]                      ;; time they stay in the nest counts down
   [ return-to-nest ]                                             ;; else they are outside and return to the nest
if nest? and nest-stay-time = 0                                   ;; if they are at the nest and their nest stay time is over
[ unload                                                          ;; they give their forager energy (positive or negative value) to the nest (changing nest energy)
  decision-of-recruitment-2 ] ]                                   ;; face decision whether to recruit to food source 2 or not

  ask happy-successful-foragers-3                                 ;; ants happy with food source 3
[  set color green                                                ;; are green
    ifelse nest?                                                  ;; if there are at nest
  [  nest-stay                                                    ;; walk slowly
     set nest-stay-time nest-stay-time - 1 ]                      ;; time they stay in the nest counts down
   [ return-to-nest ]                                             ;; else they are outside and return to the nest
if nest? and nest-stay-time = 0                                   ;; if they are at the nest and their nest stay time is over
[ unload                                                          ;; they give their forager energy (positive or negative value) to the nest (changing nest energy)
  decision-of-recruitment-3 ] ]                                   ;; face decision whether to recruit to food source 3 or not

  ask happy-successful-foragers-4                                 ;; ants happy with food source 4
[  set color green                                                ;; are green
    ifelse nest?                                                  ;; if there are at nest
  [  nest-stay                                                    ;; walk slowly
     set nest-stay-time nest-stay-time - 1 ]                      ;; time they stay in the nest counts down
   [ return-to-nest ]                                             ;; else they are outside and return to the nest
if nest? and nest-stay-time = 0                                   ;; if they are at the nest and their nest stay time is over
[ unload                                                          ;; they give their forager energy (positive or negative value) to the nest (changing nest energy)
  decision-of-recruitment-4 ] ]                                   ;; face decision whether to recruit to food source 4 or not

  ask happy-successful-foragers-5                                 ;; ants happy with food source 5
[  set color green                                                ;; are green
    ifelse nest?                                                  ;; if there are at nest
  [  nest-stay                                                    ;; walk slowly
     set nest-stay-time nest-stay-time - 1 ]                      ;; time they stay in the nest counts down
   [ return-to-nest ]                                             ;; else they are outside and return to the nest
if nest? and nest-stay-time = 0                                   ;; if they are at the nest and their nest stay time is over
[ unload                                                          ;; they give their forager energy (positive or negative value) to the nest (changing nest energy)
  decision-of-recruitment-5 ] ]                                   ;; face decision whether to recruit to food source 5 or not

  ask happy-successful-foragers-6                                 ;; ants happy with food source 6
[  set color green                                                ;; are green
    ifelse nest?                                                  ;; if there are at nest
  [  nest-stay                                                    ;; walk slowly
     set nest-stay-time nest-stay-time - 1 ]                      ;; time they stay in the nest counts down
   [ return-to-nest ]                                             ;; else they are outside and return to the nest
if nest? and nest-stay-time = 0                                   ;; if they are at the nest and their nest stay time is over
[ unload                                                          ;; they give their forager energy (positive or negative value) to the nest (changing nest energy)
  decision-of-recruitment-6 ] ]                                   ;; face decision whether to recruit to food source 6 or not

  ask happy-successful-foragers-7                                 ;; ants happy with food source 7
[  set color green                                                ;; are green
    ifelse nest?                                                  ;; if there are at nest
  [  nest-stay                                                    ;; walk slowly
     set nest-stay-time nest-stay-time - 1 ]                      ;; time they stay in the nest counts down
   [ return-to-nest ]                                             ;; else they are outside and return to the nest
if nest? and nest-stay-time = 0                                   ;; if they are at the nest and their nest stay time is over
[ unload                                                          ;; they give their forager energy (positive or negative value) to the nest (changing nest energy)
  decision-of-recruitment-7 ] ]                                   ;; face decision whether to recruit to food source 7 or not

  ask happy-successful-foragers-8                                 ;; ants happy with food source 8
[  set color green                                                ;; are green
    ifelse nest?                                                  ;; if there are at nest
  [  nest-stay                                                    ;; walk slowly
     set nest-stay-time nest-stay-time - 1 ]                      ;; time they stay in the nest counts down
   [ return-to-nest ]                                             ;; else they are outside and return to the nest
if nest? and nest-stay-time = 0                                   ;; if they are at the nest and their nest stay time is over
[ unload                                                          ;; they give their forager energy (positive or negative value) to the nest (changing nest energy)
  decision-of-recruitment-8 ] ]                                   ;; face decision whether to recruit to food source 8 or not

  ask happy-successful-foragers-9                                 ;; ants happy with food source 9
[  set color green                                                ;; are green
    ifelse nest?                                                  ;; if there are at nest
  [  nest-stay                                                    ;; walk slowly
     set nest-stay-time nest-stay-time - 1 ]                      ;; time they stay in the nest counts down
   [ return-to-nest ]                                             ;; else they are outside and return to the nest
if nest? and nest-stay-time = 0                                   ;; if they are at the nest and their nest stay time is over
[ unload                                                          ;; they give their forager energy (positive or negative value) to the nest (changing nest energy)
  decision-of-recruitment-9 ] ]                                   ;; face decision whether to recruit to food source 9 or not

  ask happy-successful-foragers-10                                ;; ants happy with food source 10
[  set color green                                                ;; are green
    ifelse nest?                                                  ;; if there are at nest
  [  nest-stay                                                    ;; walk slowly
     set nest-stay-time nest-stay-time - 1 ]                      ;; time they stay in the nest counts down
   [ return-to-nest ]                                             ;; else they are outside and return to the nest
if nest? and nest-stay-time = 0                                   ;; if they are at the nest and their nest stay time is over
[ unload                                                          ;; they give their forager energy (positive or negative value) to the nest (changing nest energy)
  decision-of-recruitment-10 ] ]                                  ;; face decision whether to recruit to food source 10 or not

;;;;;;;;;;;;;;;;;;;;;;;;;;;;;;;;;;;;;;;
;;; UNSATISFIED ANTS RETURN TO NEST ;;;
;;;;;;;;;;;;;;;;;;;;;;;;;;;;;;;;;;;;;;;

    ask unsatisfied-foragers
[  set color brown                                                ;; unsuccessful ants are brown
    if patch-ahead 1 = nobody                                     ;; if they encounter a wall (non-patch)
  [ rt 180 ]                                                      ;; change direction by 180°
    ifelse nest?                                                  ;; if there are at nest
  [  nest-stay                                                    ;; walk slowly
     set nest-stay-time nest-stay-time - 1 ]                      ;; time they stay in the nest counts down
   [ return-to-nest ]                                             ;; else they are outside and return to the nest
if nest? and nest-stay-time = 0                                   ;; if they are at the nest and nest stay time is over
[ unload                                                          ;; they give their forager energy (positive or negative value) to the nest (changing nest energy)
  decision-of-foraging-strategy ] ]                               ;; face decision which foraging strategy to chose: scouting or recruiting

;;;;;;;;;;;;;;;;;;;;;;;;;;;;;;;;;;;;;;;;;;
;;; HAPPY ANTS RETURN TO RESPECTIVE FS ;;;
;;;;;;;;;;;;;;;;;;;;;;;;;;;;;;;;;;;;;;;;;;

ask successful-foragers-1
 [ set color blue                                                 ;; their colour is blue
    if if-food? = true                                            ;; if there is food
    [ set breed drinking-ants                                     ;; become drinking ants
      stop ]                                                      ;; do not "overrun" small food sources
ifelse search-time < 0                                            ;; if food source 1 is not found within search time
    [ set nest-stay-time unloading-time                           ;; set nest stay time to the adjustable unloading time, default 60 ticks
      set breed unsatisfied-foragers ]                            ;; become unsatisfied foragers
    [ Head-to-FS-1  ] ]                                           ;; else: if there is time left, head for food source 1

ask successful-foragers-2
 [ set color blue                                                 ;; their colour is blue
    if if-food? = true                                            ;; if there is food
    [ set breed drinking-ants                                     ;; become drinking ants
      stop ]                                                      ;; do not "overrun" small food sources
ifelse search-time < 0                                            ;; if food source 2 is not found within search time
    [ set nest-stay-time unloading-time                           ;; set nest stay time to the adjustable unloading time, default 60 ticks
      set breed unsatisfied-foragers ]                            ;; become unsatisfied foragers
    [ Head-to-FS-2  ] ]                                           ;; else: if there is time left, head for food source 2

  ask successful-foragers-3
 [ set color blue                                                 ;; their colour is blue
    if if-food? = true                                            ;; if there is food
    [ set breed drinking-ants                                     ;; become drinking ants
      stop ]                                                      ;; do not "overrun" small food sources
ifelse search-time < 0                                            ;; if food source 3 is not found within search time
    [ set nest-stay-time unloading-time                           ;; set nest stay time to the adjustable unloading time, default 60 ticks
      set breed unsatisfied-foragers ]                            ;; become unsatisfied foragers
    [ Head-to-FS-3  ] ]                                           ;; else: if there is time left, head for food source 3

  ask successful-foragers-4
 [ set color blue                                                 ;; their colour is blue
    if if-food? = true                                            ;; if there is food
    [ set breed drinking-ants                                     ;; become drinking ants
      stop ]                                                      ;; do not "overrun" small food sources
ifelse search-time < 0                                            ;; if food source 4 is not found within search time
    [ set nest-stay-time unloading-time                           ;; set nest stay time to the adjustable unloading time, default 60 ticks
      set breed unsatisfied-foragers ]                            ;; become unsatisfied foragers
    [ Head-to-FS-4  ] ]                                           ;; else: if there is time left, head for food source 4

  ask successful-foragers-5
 [ set color blue                                                 ;; their colour is blue
    if if-food? = true                                            ;; if there is food
    [ set breed drinking-ants                                     ;; become drinking ants
      stop ]                                                      ;; do not "overrun" small food sources
ifelse search-time < 0                                            ;; if food source 5 is not found within search time
    [ set nest-stay-time unloading-time                           ;; set nest stay time to the adjustable unloading time, default 60 ticks
      set breed unsatisfied-foragers ]                            ;; become unsatisfied foragers
    [ Head-to-FS-5  ] ]                                           ;; else: if there is time left, head for food source 5

  ask successful-foragers-6
 [ set color blue                                                 ;; their colour is blue
    if if-food? = true                                            ;; if there is food
    [ set breed drinking-ants                                     ;; become drinking ants
      stop ]                                                      ;; do not "overrun" small food sources
ifelse search-time < 0                                            ;; if food source 6 is not found within search time
    [ set nest-stay-time unloading-time                           ;; set nest stay time to the adjustable unloading time, default 60 ticks
      set breed unsatisfied-foragers ]                            ;; become unsatisfied foragers
    [ Head-to-FS-6  ] ]                                           ;; else: if there is time left, head for food source 6

  ask successful-foragers-7
 [ set color blue                                                 ;; their colour is blue
    if if-food? = true                                            ;; if there is food
    [ set breed drinking-ants                                     ;; become drinking ants
      stop ]                                                      ;; do not "overrun" small food sources
ifelse search-time < 0                                            ;; if food source 7 is not found within search time
    [ set nest-stay-time unloading-time                           ;; set nest stay time to the adjustable unloading time, default 60 ticks
      set breed unsatisfied-foragers ]                            ;; become unsatisfied foragers
    [ Head-to-FS-7  ] ]                                           ;; else: if there is time left, head for food source 7

  ask successful-foragers-8
 [ set color blue                                                 ;; their colour is blue
    if if-food? = true                                            ;; if there is food
    [ set breed drinking-ants                                     ;; become drinking ants
      stop ]                                                      ;; do not "overrun" small food sources
ifelse search-time < 0                                            ;; if food source 8 is not found within search time
    [ set nest-stay-time unloading-time                           ;; set nest stay time to the adjustable unloading time, default 60 ticks
      set breed unsatisfied-foragers ]                            ;; become unsatisfied foragers
    [ Head-to-FS-8  ] ]                                           ;; else: if there is time left, head for food source 8

  ask successful-foragers-9
 [ set color blue                                                 ;; their colour is blue
    if if-food? = true                                            ;; if there is food
    [ set breed drinking-ants                                     ;; become drinking ants
      stop ]                                                      ;; do not "overrun" small food sources
ifelse search-time < 0                                            ;; if food source 9 is not found within search time
    [ set nest-stay-time unloading-time                           ;; set nest stay time to the adjustable unloading time, default 60 ticks
      set breed unsatisfied-foragers ]                            ;; become unsatisfied foragers
    [ Head-to-FS-9  ] ]                                           ;; else: if there is time left, head for food source 9

  ask successful-foragers-10
 [ set color blue                                                 ;; their colour is blue
    if if-food? = true                                            ;; if there is food
    [ set breed drinking-ants                                     ;; become drinking ants
      stop ]                                                      ;; do not "overrun" small food sources
ifelse search-time < 0                                            ;; if food source 10 is not found within search time
    [ set nest-stay-time unloading-time                           ;; set nest stay time to the adjustable unloading time, default 60 ticks
      set breed unsatisfied-foragers ]                            ;; become unsatisfied foragers
    [ Head-to-FS-10  ] ]                                           ;; else: if there is time left, head for food source 10

;;;;;;;;;;;;;;;;;;;;;;;;;
;;; TANDEM-INITIATION ;;;
;;;;;;;;;;;;;;;;;;;;;;;;;

ask Tandemstarters-1                                              ;; happy successful foragers 1 at the nest not knowing if recruits are present
  [ set color orange                                              ;; their colour is orange
    nest-stay                                                     ;; walk slowly
    set nest-stay-time nest-stay-time - 1                         ;; time they stay in the nest counts down
ifelse any? Tandemstarters-1 and any? recruits                    ;; if there are Tandemstarters 1 and recruits at the nest
   [ Tandemformation-1 ]                                          ;; start formation of a tandem to food source 1
  [ nest-stay ]                                                   ;; else: stay in the nest
if nest-stay-time < 0                                             ;; if nest stay time is over,
  [ set breed successful-foragers-1                               ;; become successful-foragers for food source 1 regardless of recruits presence
    reset-times  ] ]                                              ;; and reset internal times

ask Tandemstarters-2                                              ;; happy successful foragers 2 at the nest not knowing if recruits are present
  [ set color orange                                              ;; their colour is orange
    nest-stay                                                     ;; walk slowly
    set nest-stay-time nest-stay-time - 1                         ;; time they stay in the nest counts down
ifelse any? Tandemstarters-2 and any? recruits                    ;; if there are Tandemstarters 2 and recruits at the nest
   [ Tandemformation-2 ]                                          ;; start formation of a tandem to food source 2
  [ nest-stay ]                                                   ;; else: stay in the nest
if nest-stay-time < 0                                             ;; if nest stay time is over,
  [ set breed successful-foragers-2                               ;; become successful-foragers for food source 2 regardless of recruits presence
    reset-times  ] ]                                              ;; and reset internal times

  ask Tandemstarters-3                                            ;; happy successful foragers 3 at the nest not knowing if recruits are present
  [ set color orange                                              ;; their colour is orange
    nest-stay                                                     ;; walk slowly
    set nest-stay-time nest-stay-time - 1                         ;; time they stay in the nest counts down
ifelse any? Tandemstarters-3 and any? recruits                    ;; if there are Tandemstarters 3 and recruits at the nest
   [ Tandemformation-3 ]                                          ;; start formation of a tandem to food source 3
  [ nest-stay ]                                                   ;; else: stay in the nest
if nest-stay-time < 0                                             ;; if nest stay time is over,
  [ set breed successful-foragers-3                               ;; become successful-foragers for food source 3 regardless of recruits presence
    reset-times  ] ]                                              ;; and reset internal times

  ask Tandemstarters-4                                            ;; happy successful foragers 4 at the nest not knowing if recruits are present
  [ set color orange                                              ;; their colour is orange
    nest-stay                                                     ;; walk slowly
    set nest-stay-time nest-stay-time - 1                         ;; time they stay in the nest counts down
ifelse any? Tandemstarters-4 and any? recruits                    ;; if there are Tandemstarters 4 and recruits at the nest
   [ Tandemformation-4 ]                                          ;; start formation of a tandem to food source 4
  [ nest-stay ]                                                   ;; else: stay in the nest
if nest-stay-time < 0                                             ;; if nest stay time is over,
  [ set breed successful-foragers-4                               ;; become successful-foragers for food source 4 regardless of recruits presence
    reset-times  ] ]                                              ;; and reset internal times

  ask Tandemstarters-5                                            ;; happy successful foragers 5 at the nest not knowing if recruits are present
  [ set color orange                                              ;; their colour is orange
    nest-stay                                                     ;; walk slowly
    set nest-stay-time nest-stay-time - 1                         ;; time they stay in the nest counts down
ifelse any? Tandemstarters-5 and any? recruits                    ;; if there are Tandemstarters 5 and recruits at the nest
   [ Tandemformation-5 ]                                          ;; start formation of a tandem to food source 5
  [ nest-stay ]                                                   ;; else: stay in the nest
if nest-stay-time < 0                                             ;; if nest stay time is over,
  [ set breed successful-foragers-5                               ;; become successful-foragers for food source 5 regardless of recruits presence
    reset-times  ] ]                                              ;; and reset internal times

  ask Tandemstarters-6                                            ;; happy successful foragers 6 at the nest not knowing if recruits are present
  [ set color orange                                              ;; their colour is orange
    nest-stay                                                     ;; walk slowly
    set nest-stay-time nest-stay-time - 1                         ;; time they stay in the nest counts down
ifelse any? Tandemstarters-6 and any? recruits                    ;; if there are Tandemstarters 6 and recruits at the nest
   [ Tandemformation-6 ]                                          ;; start formation of a tandem to food source 6
  [ nest-stay ]                                                   ;; else: stay in the nest
if nest-stay-time < 0                                             ;; if nest stay time is over,
  [ set breed successful-foragers-6                               ;; become successful-foragers for food source 6 regardless of recruits presence
    reset-times  ] ]                                              ;; and reset internal times

  ask Tandemstarters-7                                            ;; happy successful foragers 7 at the nest not knowing if recruits are present
  [ set color orange                                              ;; their colour is orange
    nest-stay                                                     ;; walk slowly
    set nest-stay-time nest-stay-time - 1                         ;; time they stay in the nest counts down
ifelse any? Tandemstarters-7 and any? recruits                    ;; if there are Tandemstarters 7 and recruits at the nest
   [ Tandemformation-7 ]                                          ;; start formation of a tandem to food source 7
  [ nest-stay ]                                                   ;; else: stay in the nest
if nest-stay-time < 0                                             ;; if nest stay time is over,
  [ set breed successful-foragers-7                               ;; become successful-foragers for food source 7 regardless of recruits presence
    reset-times  ] ]                                              ;; and reset internal times

  ask Tandemstarters-8                                            ;; happy successful foragers 8 at the nest not knowing if recruits are present
  [ set color orange                                              ;; their colour is orange
    nest-stay                                                     ;; walk slowly
    set nest-stay-time nest-stay-time - 1                         ;; time they stay in the nest counts down
ifelse any? Tandemstarters-8 and any? recruits                    ;; if there are Tandemstarters 8 and recruits at the nest
   [ Tandemformation-8 ]                                          ;; start formation of a tandem to food source 8
  [ nest-stay ]                                                   ;; else: stay in the nest
if nest-stay-time < 0                                             ;; if nest stay time is over,
  [ set breed successful-foragers-8                               ;; become successful-foragers for food source 8 regardless of recruits presence
    reset-times  ] ]                                              ;; and reset internal times

  ask Tandemstarters-9                                            ;; happy successful foragers 9 at the nest not knowing if recruits are present
  [ set color orange                                              ;; their colour is orange
    nest-stay                                                     ;; walk slowly
    set nest-stay-time nest-stay-time - 1                         ;; time they stay in the nest counts down
ifelse any? Tandemstarters-9 and any? recruits                    ;; if there are Tandemstarters 9 and recruits at the nest
   [ Tandemformation-9 ]                                          ;; start formation of a tandem to food source 9
  [ nest-stay ]                                                   ;; else: stay in the nest
if nest-stay-time < 0                                             ;; if nest stay time is over,
  [ set breed successful-foragers-9                               ;; become successful-foragers for food source 9 regardless of recruits presence
    reset-times  ] ]                                              ;; and reset internal times

  ask Tandemstarters-10                                           ;; happy successful foragers 10 at the nest not knowing if recruits are present
  [ set color orange                                              ;; their colour is orange
    nest-stay                                                     ;; walk slowly
    set nest-stay-time nest-stay-time - 1                         ;; time they stay in the nest counts down
ifelse any? Tandemstarters-10 and any? recruits                   ;; if there are Tandemstarters 10 and recruits at the nest
   [ Tandemformation-10 ]                                         ;; start formation of a tandem to food source 10
  [ nest-stay ]                                                   ;; else: stay in the nest
if nest-stay-time < 0                                             ;; if nest stay time is over,
  [ set breed successful-foragers-10                              ;; become successful-foragers for food source 10 regardless of recruits presence
    reset-times  ] ]                                              ;; and reset internal times

;;;;;;;;;;;;;;;;;;;;;;
;;; TANDEM TO FS-1 ;;;
;;;;;;;;;;;;;;;;;;;;;;

ask Tandemleaders-1                                               ;; recruiters at  the nest, leading the tandem to food source 1
  [ set color cyan                                                ;; their colour is cyan
    if if-food? = true                                            ;; if there is food
    [ set breed drinking-ants                                     ;; become drinking ants
      stop ]                                                      ;; do not "overrun" small food sources
ifelse search-time < 0                                            ;; if food source 1 is not found within search time
    [  set breed unsatisfied-foragers ]                           ;; become unsatisfied foragers
    [ tandem-FS-1  ] ]                                            ;; else: if there is time left, head for food source 1

ask Tandemfollowers-1                                             ;; former recruits at the nest
    [ set color cyan                                              ;; their colour is cyan
    set forager-timer forager-timer + 1                           ;; timer for time spend in the tandem advances
     tandem-breakup                                               ;; 5 ‰ chance per tick to loose food source direction
    if if-food? = true                                            ;; if there is food
    [ output-show forager-timer                                   ;; trigger the timer to display their time spend in the tandem
    set breed drinking-ants                                       ;; become drinking ants
      stop ]                                                      ;; do not "overrun" small food sources
ifelse search-time < 0                                            ;; if food source 1 is not found within search time
    [  set breed unsatisfied-foragers ]                           ;; become unsatisfied foragers
          [ tandem-FS-1  ] ]                                      ;; else: if there is time left, head for food source 1

;;;;;;;;;;;;;;;;;;;;;;
;;; TANDEM TO FS-2 ;;;
;;;;;;;;;;;;;;;;;;;;;;

ask Tandemleaders-2                                               ;; recruiters at  the nest, leading the tandem to food source 2
  [ set color cyan                                                ;; their colour is cyan
    if if-food? = true                                            ;; if there is food
    [ set breed drinking-ants                                     ;; become drinking ants
      stop ]                                                      ;; do not "overrun" small food sources
ifelse search-time < 0                                            ;; if food source 2 is not found within search time
    [  set breed unsatisfied-foragers ]                           ;; become unsatisfied foragers
    [ tandem-FS-2  ] ]                                            ;; else: if there is time left, head for food source 2

ask Tandemfollowers-2                                             ;; former recruits at the nest
    [ set color cyan                                              ;; their colour is cyan
    set forager-timer forager-timer + 1                           ;; timer for time spend in the tandem advances
     tandem-breakup                                               ;; 5 ‰ chance per tick to loose food source direction
    if if-food? = true                                            ;; if there is food
    [ output-show forager-timer                                   ;; trigger the timer to display their time spend in the tandem
     set breed drinking-ants                                      ;; become drinking ants
      stop ]                                                      ;; do not "overrun" small food sources
ifelse search-time < 0                                            ;; if food source 2 is not found within search time
    [  set breed unsatisfied-foragers ]                           ;; become unsatisfied foragers
    [ tandem-FS-2  ] ]                                            ;; else: if there is time left, head for food source 2

;;;;;;;;;;;;;;;;;;;;;;
;;; TANDEM TO FS-3 ;;;
;;;;;;;;;;;;;;;;;;;;;;

ask Tandemleaders-3                                               ;; recruiters at  the nest, leading the tandem to food source 3
  [ set color cyan                                                ;; their colour is cyan
    if if-food? = true                                            ;; if there is food
    [ set breed drinking-ants                                     ;; become drinking ants
      stop ]                                                      ;; do not "overrun" small food sources
ifelse search-time < 0                                            ;; if food source 3 is not found within search time
    [  set breed unsatisfied-foragers ]                           ;; become unsatisfied foragers
    [ tandem-FS-3  ] ]                                            ;; else: if there is time left, head for food source 3

ask Tandemfollowers-3                                             ;; former recruits at the nest
    [ set color cyan                                              ;; their colour is cyan
    set forager-timer forager-timer + 1                           ;; timer for time spend in the tandem advances
     tandem-breakup                                               ;; 5 ‰ chance per tick to loose food source direction
    if if-food? = true                                            ;; if there is food
    [ output-show forager-timer                                   ;; trigger the timer to display their time spend in the tandem
    set breed drinking-ants                                       ;; become drinking ants
      stop ]                                                      ;; do not "overrun" small food sources
ifelse search-time < 0                                            ;; if food source 3 is not found within search time
    [  set breed unsatisfied-foragers ]                           ;; become unsatisfied foragers
          [ tandem-FS-3  ] ]                                      ;; else: if there is time left, head for food source 3

;;;;;;;;;;;;;;;;;;;;;;
;;; TANDEM TO FS-4 ;;;
;;;;;;;;;;;;;;;;;;;;;;

ask Tandemleaders-4                                               ;; recruiters at  the nest, leading the tandem to food source 4
  [ set color cyan                                                ;; their colour is cyan
    if if-food? = true                                            ;; if there is food
    [ set breed drinking-ants                                     ;; become drinking ants
      stop ]                                                      ;; do not "overrun" small food sources
ifelse search-time < 0                                            ;; if food source 4 is not found within search time
    [  set breed unsatisfied-foragers ]                           ;; become unsatisfied foragers
    [ tandem-FS-4  ] ]                                            ;; else: if there is time left, head for food source 4

ask Tandemfollowers-4                                             ;; former recruits at the nest
    [ set color cyan                                              ;; their colour is cyan
    set forager-timer forager-timer + 1                           ;; timer for time spend in the tandem advances
     tandem-breakup                                               ;; 5 ‰ chance per tick to loose food source direction
    if if-food? = true                                            ;; if there is food
    [ output-show forager-timer                                   ;; trigger the timer to display their time spend in the tandem
    set breed drinking-ants                                       ;; become drinking ants
      stop ]                                                      ;; do not "overrun" small food sources
ifelse search-time < 0                                            ;; if food source 4 is not found within search time
    [  set breed unsatisfied-foragers ]                           ;; become unsatisfied foragers
          [ tandem-FS-4  ] ]                                      ;; else: if there is time left, head for food source 4

;;;;;;;;;;;;;;;;;;;;;;
;;; TANDEM TO FS-5 ;;;
;;;;;;;;;;;;;;;;;;;;;;

ask Tandemleaders-5                                               ;; recruiters at  the nest, leading the tandem to food source 5
  [ set color cyan                                                ;; their colour is cyan
    if if-food? = true                                            ;; if there is food
    [ set breed drinking-ants                                     ;; become drinking ants
      stop ]                                                      ;; do not "overrun" small food sources
ifelse search-time < 0                                            ;; if food source 1 is not found within search time
    [  set breed unsatisfied-foragers ]                           ;; become unsatisfied foragers
    [ tandem-FS-5  ] ]                                            ;; else: if there is time left, head for food source 5

ask Tandemfollowers-5                                             ;; former recruits at the nest
    [ set color cyan                                              ;; their colour is cyan
    set forager-timer forager-timer + 1                           ;; timer for time spend in the tandem advances
     tandem-breakup                                               ;; 5 ‰ chance per tick to loose food source direction
    if if-food? = true                                            ;; if there is food
    [ output-show forager-timer                                   ;; trigger the timer to display their time spend in the tandem
    set breed drinking-ants                                       ;; become drinking ants
      stop ]                                                      ;; do not "overrun" small food sources
ifelse search-time < 0                                            ;; if food source 5 is not found within search time
    [  set breed unsatisfied-foragers ]                           ;; become unsatisfied foragers
          [ tandem-FS-5  ] ]                                      ;; else: if there is time left, head for food source 5

;;;;;;;;;;;;;;;;;;;;;;
;;; TANDEM TO FS-6 ;;;
;;;;;;;;;;;;;;;;;;;;;;

ask Tandemleaders-6                                               ;; recruiters at  the nest, leading the tandem to food source 6
  [ set color cyan                                                ;; their colour is cyan
    if if-food? = true                                            ;; if there is food
    [ set breed drinking-ants                                     ;; become drinking ants
      stop ]                                                      ;; do not "overrun" small food sources
ifelse search-time < 0                                            ;; if food source 6 is not found within search time
    [  set breed unsatisfied-foragers ]                           ;; become unsatisfied foragers
    [ tandem-FS-6  ] ]                                            ;; else: if there is time left, head for food source 6

ask Tandemfollowers-6                                             ;; former recruits at the nest
    [ set color cyan                                              ;; their colour is cyan
    set forager-timer forager-timer + 1                           ;; timer for time spend in the tandem advances
     tandem-breakup                                               ;; 5 ‰ chance per tick to loose food source direction
    if if-food? = true                                            ;; if there is food
    [ output-show forager-timer                                   ;; trigger the timer to display their time spend in the tandem
    set breed drinking-ants                                       ;; become drinking ants
      stop ]                                                      ;; do not "overrun" small food sources
ifelse search-time < 0                                            ;; if food source 6 is not found within search time
    [  set breed unsatisfied-foragers ]                           ;; become unsatisfied foragers
          [ tandem-FS-6  ] ]                                      ;; else: if there is time left, head for food source 6

;;;;;;;;;;;;;;;;;;;;;;
;;; TANDEM TO FS-7 ;;;
;;;;;;;;;;;;;;;;;;;;;;

ask Tandemleaders-7                                               ;; recruiters at  the nest, leading the tandem to food source 7
  [ set color cyan                                                ;; their colour is cyan
    if if-food? = true                                            ;; if there is food
    [ set breed drinking-ants                                     ;; become drinking ants
      stop ]                                                      ;; do not "overrun" small food sources
ifelse search-time < 0                                            ;; if food source 1 is not found within search time
    [  set breed unsatisfied-foragers ]                           ;; become unsatisfied foragers
    [ tandem-FS-7  ] ]                                            ;; else: if there is time left, head for food source 7

ask Tandemfollowers-7                                             ;; former recruits at the nest
    [ set color cyan                                              ;; their colour is cyan
    set forager-timer forager-timer + 1                           ;; timer for time spend in the tandem advances
     tandem-breakup                                               ;; 5 ‰ chance per tick to loose food source direction
    if if-food? = true                                            ;; if there is food
    [ output-show forager-timer                                   ;; trigger the timer to display their time spend in the tandem
    set breed drinking-ants                                       ;; become drinking ants
      stop ]                                                      ;; do not "overrun" small food sources
ifelse search-time < 0                                            ;; if food source 7 is not found within search time
    [  set breed unsatisfied-foragers ]                           ;; become unsatisfied foragers
          [ tandem-FS-7  ] ]                                      ;; else: if there is time left, head for food source 7

;;;;;;;;;;;;;;;;;;;;;;
;;; TANDEM TO FS-8 ;;;
;;;;;;;;;;;;;;;;;;;;;;

ask Tandemleaders-8                                               ;; recruiters at  the nest, leading the tandem to food source 8
  [ set color cyan                                                ;; their colour is cyan
    if if-food? = true                                            ;; if there is food
    [ set breed drinking-ants                                     ;; become drinking ants
      stop ]                                                      ;; do not "overrun" small food sources
ifelse search-time < 0                                            ;; if food source 8 is not found within search time
    [  set breed unsatisfied-foragers ]                           ;; become unsatisfied foragers
    [ tandem-FS-8  ] ]                                            ;; else: if there is time left, head for food source 8

ask Tandemfollowers-8                                             ;; former recruits at the nest
    [ set color cyan                                              ;; their colour is cyan
    set forager-timer forager-timer + 1                           ;; timer for time spend in the tandem advances
     tandem-breakup                                               ;; 5 ‰ chance per tick to loose food source direction
    if if-food? = true                                            ;; if there is food
    [ output-show forager-timer                                   ;; trigger the timer to display their time spend in the tandem
    set breed drinking-ants                                       ;; become drinking ants
      stop ]                                                      ;; do not "overrun" small food sources
ifelse search-time < 0                                            ;; if food source 8 is not found within search time
    [  set breed unsatisfied-foragers ]                           ;; become unsatisfied foragers
          [ tandem-FS-8  ] ]                                      ;; else: if there is time left, head for food source 8

;;;;;;;;;;;;;;;;;;;;;;
;;; TANDEM TO FS-9 ;;;
;;;;;;;;;;;;;;;;;;;;;;

ask Tandemleaders-9                                               ;; recruiters at  the nest, leading the tandem to food source 9
  [ set color cyan                                                ;; their colour is cyan
    if if-food? = true                                            ;; if there is food
    [ set breed drinking-ants                                     ;; become drinking ants
      stop ]                                                      ;; do not "overrun" small food sources
ifelse search-time < 0                                            ;; if food source 1 is not found within search time
    [  set breed unsatisfied-foragers ]                           ;; become unsatisfied foragers
    [ tandem-FS-9  ] ]                                            ;; else: if there is time left, head for food source 9

ask Tandemfollowers-9                                             ;; former recruits at the nest
    [ set color cyan                                              ;; their colour is cyan
    set forager-timer forager-timer + 1                           ;; timer for time spend in the tandem advances
     tandem-breakup                                               ;; 5 ‰ chance per tick to loose food source direction
    if if-food? = true                                            ;; if there is food
    [ output-show forager-timer                                   ;; trigger the timer to display their time spend in the tandem
    set breed drinking-ants                                       ;; become drinking ants
      stop ]                                                      ;; do not "overrun" small food sources
ifelse search-time < 0                                            ;; if food source 9 is not found within search time
    [  set breed unsatisfied-foragers ]                           ;; become unsatisfied foragers
          [ tandem-FS-9  ] ]                                      ;; else: if there is time left, head for food source 9

;;;;;;;;;;;;;;;;;;;;;;
;;; TANDEM TO FS-10 ;;;
;;;;;;;;;;;;;;;;;;;;;;

ask Tandemleaders-10                                              ;; recruiters at  the nest, leading the tandem to food source 10
  [ set color cyan                                                ;; their colour is cyan
    if if-food? = true                                            ;; if there is food
    [ set breed drinking-ants                                     ;; become drinking ants
      stop ]                                                      ;; do not "overrun" small food sources
ifelse search-time < 0                                            ;; if food source 10 is not found within search time
    [  set breed unsatisfied-foragers ]                           ;; become unsatisfied foragers
    [ tandem-FS-10  ] ]                                           ;; else: if there is time left, head for food source 10

ask Tandemfollowers-10                                            ;; former recruits at the nest
    [ set color cyan                                              ;; their colour is cyan
    set forager-timer forager-timer + 1                           ;; timer for time spend in the tandem advances
     tandem-breakup                                               ;; 5 ‰ chance per tick to loose food source direction
    if if-food? = true                                            ;; if there is food
    [ output-show forager-timer                                   ;; trigger the timer to display their time spend in the tandem
    set breed drinking-ants                                       ;; become drinking ants
      stop ]                                                      ;; do not "overrun" small food sources
ifelse search-time < 0                                            ;; if food source 10 is not found within search time
    [  set breed unsatisfied-foragers ]                           ;; become unsatisfied foragers
          [ tandem-FS-10  ] ]                                     ;; else: if there is time left, head for food source 10

;;;;;;;;;;;;;;;;;;;;;;;;;;;
;;; LOST TANDEM-RUNNERS ;;;
;;;;;;;;;;;;;;;;;;;;;;;;;;;

ask lost-TRs                                                      ;; tandem followers which have lost food source direction
  [  if patch-ahead 1 = nobody                                    ;; if they encounter a wall (non-patch)
    [ rt 180 ]                                                    ;; change direction by 180°
    if if-food? = true                                            ;; if there is food
    [ output-show forager-timer                                   ;; trigger the timer to display their time spend in the tandem
    set breed drinking-ants                                       ;; become drinking ants
      stop ]                                                      ;; do not "overrun" small food sources
     ifelse nest?                                                 ;; if there are at nest
  [   nest-stay                                                   ;; and walk slowly
     set nest-stay-time nest-stay-time - 1                        ;; time they stay in the nest counts down
ifelse nest-stay-time = 0                                         ;; if they are at the nest and nest stay time is over
[ unload                                                          ;; they give their forager energy (positive or negative value) to the nest (changing nest energy)
  decision-of-foraging-strategy ]                                 ;; face decision which foraging strategy to chose: scouting or recruiting
        [ nest-stay  ] ]                                          ;; else: if there is in the nest left stay at the nest
   [ wiggle                                                       ;; randomly change direction ("searching the food source")
    fd (velocity-outside / 2 )                                    ;; move with half speed/tandem-speed
    set forager-energy forager-energy - metabolic-cost            ;; and with energy-costs
    set forager-timer forager-timer + 1                           ;; timer for time spend searching a food source advances
        set search-time search-time - 1 ]                         ;; time they search for a food source counts down
 if (search-time) < 0                                             ;; if a food source is not found within search time
            [  decision-of-lost-TRS ] ]                           ;; decide on becoming a scout or return to nest

;;;;;;;;;;;;;;;;;;
;;; FS-ACTIONS ;;;
;;;;;;;;;;;;;;;;;;

   FS-Disappearence                                               ;; food sources disappear when having no energy

  if not any? patches with [ FS-number = 1 ]                      ;; if there is no food source with the number 1
  [ ask patches                                                   ;; ask all patches
   [ set FS-timer-1  FS-timer-1  - 1                              ;; to count-down the timer for food source 1
     if FS-timer-1  = 0                                           ;; if the count-down is over
      [  FS-Renewal-1                                             ;; let food source 1 reappear
         set FS-timer-1  FS-timer ] ] ]                           ;; reset timer for food source 1

  if not any? patches with [ FS-number = 2 ]                      ;; if there is no food source with the number 2
  [ ask patches                                                   ;; ask all patches
   [ set FS-timer-2  FS-timer-2  - 1                              ;; to count-down the timer for food source 2
     if FS-timer-2  = 0                                           ;; if the count-down is over
      [  FS-Renewal-2                                             ;; let food source 2 reappear
         set FS-timer-2  FS-timer ] ] ]                           ;; reset timer for food source 2

    if not any? patches with [ FS-number = 3 ]                    ;; if there is no food source with the number 3
  [ ask patches                                                   ;; ask all patches
   [ set FS-timer-3  FS-timer-3  - 1                              ;; to count-down the timer for food source 3
     if FS-timer-3  = 0                                           ;; if the count-down is over
      [  FS-Renewal-3                                             ;; let food source 3 reappear
         set FS-timer-3  FS-timer ] ] ]                           ;; reset timer for food source 3

    if not any? patches with [ FS-number = 4 ]                    ;; if there is no food source with the number 4
  [ ask patches                                                   ;; ask all patches
   [ set FS-timer-4  FS-timer-4  - 1                              ;; to count-down the timer for food source 4
     if FS-timer-4  = 0                                           ;; if the count-down is over
      [  FS-Renewal-4                                             ;; let food source 4 reappear
         set FS-timer-4  FS-timer ] ] ]                           ;; reset timer for food source 4

    if not any? patches with [ FS-number = 5 ]                    ;; if there is no food source with the number 5
  [ ask patches                                                   ;; ask all patches
   [ set FS-timer-5  FS-timer-5  - 1                              ;; to count-down the timer for food source 5
     if FS-timer-5  = 0                                           ;; if the count-down is over
      [  FS-Renewal-5                                             ;; let food source 5 reappear
         set FS-timer-5  FS-timer ] ] ]                           ;; reset timer for food source 5

    if not any? patches with [ FS-number = 6 ]                    ;; if there is no food source with the number 6
  [ ask patches                                                   ;; ask all patches
   [ set FS-timer-6  FS-timer-6  - 1                              ;; to count-down the timer for food source 6
     if FS-timer-6  = 0                                           ;; if the count-down is over
      [  FS-Renewal-6                                             ;; let food source 6 reappear
         set FS-timer-6  FS-timer ] ] ]                           ;; reset timer for food source 6

    if not any? patches with [ FS-number = 7 ]                    ;; if there is no food source with the number 7
  [ ask patches                                                   ;; ask all patches
   [ set FS-timer-7  FS-timer-7  - 1                              ;; to count-down the timer for food source 7
     if FS-timer-7  = 0                                           ;; if the count-down is over
      [  FS-Renewal-7                                             ;; let food source 7 reappear
         set FS-timer-7  FS-timer ] ] ]                           ;; reset timer for food source 7

    if not any? patches with [ FS-number = 8 ]                    ;; if there is no food source with the number 8
  [ ask patches                                                   ;; ask all patches
   [ set FS-timer-8  FS-timer-8  - 1                              ;; to count-down the timer for food source 8
     if FS-timer-8  = 0                                           ;; if the count-down is over
      [  FS-Renewal-8                                             ;; let food source 8 reappear
         set FS-timer-8  FS-timer ] ] ]                           ;; reset timer for food source 8

    if not any? patches with [ FS-number = 9 ]                    ;; if there is no food source with the number 9
  [ ask patches                                                   ;; ask all patches
   [ set FS-timer-9  FS-timer-9  - 1                              ;; to count-down the timer for food source 9
     if FS-timer-9  = 0                                           ;; if the count-down is over
      [  FS-Renewal-9                                             ;; let food source 9 reappear
         set FS-timer-9  FS-timer ] ] ]                           ;; reset timer for food source 9

    if not any? patches with [ FS-number = 10 ]                   ;; if there is no food source with the number 10
  [ ask patches                                                   ;; ask all patches
   [ set FS-timer-10  FS-timer-10  - 1                            ;; to count-down the timer for food source 10
     if FS-timer-10  = 0                                          ;; if the count-down is over
      [  FS-Renewal-10                                            ;; let food source 10 reappear
         set FS-timer-10  FS-timer ] ] ]                          ;; reset timer for food source 10

plot nest-energy                                                  ;; plot global variable nest energy

tick                                                              ;; advance tick counter

if ticks = 5400 = true [ export-output "Forager-Times.csv" ]      ;; after 5400 ticks write Timer-data of output-window to a file name "Forager-Times"

end

;;;;;;;;;;;;;;;;;;;;;;;;;
;;; MOVING PROCEDURES ;;;
;;;;;;;;;;;;;;;;;;;;;;;;;

to wiggle                                                         ;; general random orientation procedure: ant "chooses" random angle of orientation
rt random 45                                                      ;; random right angle
lt random 45                                                      ;; random left angle
end

to movement                                                       ;; general movement procedure
 ifelse nest?                                                     ;; if there are at nest
  [ ifelse breed = scouts or breed = successful-foragers-1        ;; and are scouts or successful-foragers
    or breed = successful-foragers-2 or                           ;; to any of the food sources
    breed = successful-foragers-3 or breed = successful-foragers-4;; to any of the food sources
    or breed = successful-foragers-5 or                           ;; to any of the food sources
    breed = successful-foragers-6 or breed = successful-foragers-7;; to any of the food sources
    or breed = successful-foragers-8 or                           ;; to any of the food sources
    breed = successful-foragers-9 or                              ;; to any of the food sources
    breed = successful-foragers-10                                ;; to any of the food sources
  [ fd velocity-outside ]                                         ;; move without energy-costs but fast
    [ fd velocity-nest ] ]                                        ;; else there are other breeds: walk slowly inside the nest
[ fd velocity-outside                                             ;; else the agents are outside: move with fast speed
  set forager-energy forager-energy - metabolic-cost ]            ;; and with energy-costs
end

to scouting                                                       ;; general procedure of scouts
ifelse patch-ahead 1 = nobody                                     ;; if they encounter a wall (non-patch)
    [ rt 180 ]                                                    ;; change direction by 180°
  [ wiggle                                                        ;; random change of orientation
   movement                                                       ;; general movement
   set scouts-time scouts-time + 1                                ;; advance scouting time
   set forager-timer forager-timer + 1 ]                          ;; timer for time spend scouting advances
 end

to nest-stay                                                      ;; general procedure of ants staying in the nest
ifelse [nest?] of patch-ahead 1 = true                            ;; if they are at the nest
 [ wiggle                                                         ;; randomly change direction inside the nest
   movement ]                                                     ;; walk with slow velocity and without energy-costs
 [ rt 180                                                         ;; else: if not in the nest, turn around 180 degrees for not leaving the nest
   return-to-nest ]
end

to return-to-nest                                                 ;; return to the nest
  movement                                                        ;; general movement
  nest-scent-direction                                            ;; "smell" nest-scent
end

to nest-scent-direction                                           ;; foragers go where the strongest nest-scent is
   let scent-ahead nest-scent-at-angle   0                        ;; local variable smell ahead of forager shall be nest-scent (at angle 0)
   let scent-right nest-scent-at-angle  45                        ;; local variable smell right of forager shall be nest-scent (at angle 45)
   let scent-left  nest-scent-at-angle -45                        ;; local variable smell left of forager shall be nest-scent (at angle -45)
if (scent-right > scent-ahead) or (scent-left > scent-ahead)      ;; if the scent to the right is stronger or the scent to the left is stronger than the one straight forward
[ ifelse scent-right > scent-left                                 ;; if the scent to the right is stronger than the one to the left
 [ rt 45 ]                                                        ;; the forager turns right by number degrees: If the number is negative, it turns left (follow only positive smell)
 [ lt 45 ] ]                                                      ;; else the forager turns left by number degrees: If the number is negative, it turns right (follow only positive smell)
end

to-report nest-scent-at-angle [angle]                             ;; reporter of the nest-scent-variable, feedback on how strong the nest scent is
 let p patch-right-and-ahead angle 1                              ;; p is a local variable and set to: The forager "looks" 1 degrees right of its current heading
 if p = nobody [ report 0 ]                                       ;; if the patch at this angle is non-existing (outside the arena), then it reports 0
 report [nest-scent] of p                                         ;; otherwise it reports the nest-scent of the "foragers look"
end

;;;;;;;;;;;;;;;;;;;;;;
;;; MOVEMENT TO FS ;;;
;;;;;;;;;;;;;;;;;;;;;;

to Head-to-FS-1                                                    ;; successful foragers move to food source 1
FS-1-direction                                                     ;; "smell" FS-1
movement                                                           ;; general movement
set search-time search-time - 1                                    ;; time they search for food source 1 counts down
end

to Head-to-FS-2                                                    ;; successful foragers move to food source 2
FS-2-direction                                                     ;; "smell" FS-2
movement                                                           ;; general movement
set search-time search-time - 1                                    ;; time they search for food source 2 counts down
end

to Head-to-FS-3                                                    ;; successful foragers move to food source 3
FS-3-direction                                                     ;; "smell" FS-3
movement                                                           ;; general movement
set search-time search-time - 1                                    ;; time they search for food source 3 counts down
end

to Head-to-FS-4                                                    ;; successful foragers move to food source 4
FS-4-direction                                                     ;; "smell" FS-4
movement                                                           ;; general movement
set search-time search-time - 1                                    ;; time they search for food source 4 counts down
end

to Head-to-FS-5                                                    ;; successful foragers move to food source 5
FS-5-direction                                                     ;; "smell" FS-5
movement                                                           ;; general movement
set search-time search-time - 1                                    ;; time they search for food source 5 counts down
end

to Head-to-FS-6                                                    ;; successful foragers move to food source 6
FS-6-direction                                                     ;; "smell" FS-6
movement                                                           ;; general movement
set search-time search-time - 1                                    ;; time they search for food source 6 counts down
end

to Head-to-FS-7                                                    ;; successful foragers move to food source 7
FS-7-direction                                                     ;; "smell" FS-7
movement                                                           ;; general movement
set search-time search-time - 1                                    ;; time they search for food source 7 counts down
end

to Head-to-FS-8                                                    ;; successful foragers move to food source 8
FS-8-direction                                                     ;; "smell" FS-8
movement                                                           ;; general movement
set search-time search-time - 1                                    ;; time they search for food source 8 counts down
end

to Head-to-FS-9                                                    ;; successful foragers move to food source 9
FS-9-direction                                                     ;; "smell" FS-9
movement                                                           ;; general movement
set search-time search-time - 1                                    ;; time they search for food source 9 counts down
end

to Head-to-FS-10                                                   ;; successful foragers move to food source 10
FS-10-direction                                                    ;; "smell" FS-10
movement                                                           ;; general movement
set search-time search-time - 1                                    ;; time they search for food source 10 counts down
end

;;;;;;;;;;;;;;;;;;;;;;;;;;;;;;;;
;;; MOVEMENT OF TANDEM to FS ;;;
;;;;;;;;;;;;;;;;;;;;;;;;;;;;;;;;

to tandem-FS-1                                                     ;; Tandem-1 heads to FS-1 together
ifelse nest?                                                       ;; if there are at nest
  [ FS-1-direction                                                 ;; "smell" FS-1
    fd (velocity-outside / 2 )                                     ;; move with half speed ("tandem-speed") to FS-1
    set search-time search-time - 1 ]                              ;; time they search for FS-1 counts down
 [  FS-1-direction                                                 ;; "smell" FS-1
    fd (velocity-outside / 2 )                                     ;; move with half speed ("tandem-speed") to FS-1
    set forager-energy forager-energy - metabolic-cost             ;; and with energy-costs
    set search-time search-time - 1 ]                              ;; time they search for FS-1 counts down
end

to tandem-FS-2                                                     ;; Tandem-2 heads to FS-2 together
ifelse nest?                                                       ;; if there are at nest
  [ FS-2-direction                                                 ;; "smell" FS-2
    fd (velocity-outside / 2 )                                     ;; move with half speed ("tandem-speed") to FS-2
    set search-time search-time - 1 ]                              ;; time they search for FS-1 counts down
 [  FS-2-direction                                                 ;; "smell" FS-2
    fd (velocity-outside / 2 )                                     ;; move with half speed ("tandem-speed") to FS-2
    set forager-energy forager-energy - metabolic-cost             ;; and with energy-costs
    set search-time search-time - 1 ]                              ;; time they search for FS-2 counts down
end

to tandem-FS-3                                                     ;; Tandem-3 heads to FS-3 together
ifelse nest?                                                       ;; if there are at nest
  [ FS-3-direction                                                 ;; "smell" FS-3
    fd (velocity-outside / 2 )                                     ;; move with half speed ("tandem-speed") to FS-3
    set search-time search-time - 1 ]                              ;; time they search for FS-3 counts down
 [  FS-3-direction                                                 ;; "smell" FS-3
    fd (velocity-outside / 2 )                                     ;; move with half speed ("tandem-speed") to FS-3
    set forager-energy forager-energy - metabolic-cost             ;; and with energy-costs
    set search-time search-time - 1 ]                              ;; time they search for FS-3 counts down
end

to tandem-FS-4                                                     ;; Tandem-4 heads to FS-4 together
ifelse nest?                                                       ;; if there are at nest
  [ FS-4-direction                                                 ;; "smell" FS-4
    fd (velocity-outside / 2 )                                     ;; move with half speed ("tandem-speed") to FS-4
    set search-time search-time - 1 ]                              ;; time they search for FS-1 counts down
 [  FS-4-direction                                                 ;; "smell" FS-4
    fd (velocity-outside / 2 )                                     ;; move with half speed ("tandem-speed") to FS-4
    set forager-energy forager-energy - metabolic-cost             ;; and with energy-costs
    set search-time search-time - 1 ]                              ;; time they search for FS-4 counts down
end

to tandem-FS-5                                                     ;; Tandem-5 heads to FS-5 together
ifelse nest?                                                       ;; if there are at nest
  [ FS-5-direction                                                 ;; "smell" FS-5
    fd (velocity-outside / 2 )                                     ;; move with half speed ("tandem-speed") to FS-5
    set search-time search-time - 1 ]                              ;; time they search for FS-1 counts down
 [  FS-5-direction                                                 ;; "smell" FS-5
    fd (velocity-outside / 2 )                                     ;; move with half speed ("tandem-speed") to FS-5
    set forager-energy forager-energy - metabolic-cost             ;; and with energy-costs
    set search-time search-time - 1 ]                              ;; time they search for FS-5 counts down
end

to tandem-FS-6                                                     ;; Tandem-6 heads to FS-6 together
ifelse nest?                                                       ;; if there are at nest
  [ FS-6-direction                                                 ;; "smell" FS-6
    fd (velocity-outside / 2 )                                     ;; move with half speed ("tandem-speed") to FS-6
    set search-time search-time - 1 ]                              ;; time they search for FS-6 counts down
 [  FS-6-direction                                                 ;; "smell" FS-6
    fd (velocity-outside / 2 )                                     ;; move with half speed ("tandem-speed") to FS-6
    set forager-energy forager-energy - metabolic-cost             ;; and with energy-costs
    set search-time search-time - 1 ]                              ;; time they search for FS-6 counts down
end

to tandem-FS-7                                                     ;; Tandem-7 heads to FS-7 together
ifelse nest?                                                       ;; if there are at nest
  [ FS-7-direction                                                 ;; "smell" FS-7
    fd (velocity-outside / 2 )                                     ;; move with half speed ("tandem-speed") to FS-7
    set search-time search-time - 1 ]                              ;; time they search for FS-1 counts down
 [  FS-7-direction                                                 ;; "smell" FS-7
    fd (velocity-outside / 2 )                                     ;; move with half speed ("tandem-speed") to FS-7
    set forager-energy forager-energy - metabolic-cost             ;; and with energy-costs
    set search-time search-time - 1 ]                              ;; time they search for FS-7 counts down
end

to tandem-FS-8                                                     ;; Tandem-8 heads to FS-8 together
ifelse nest?                                                       ;; if there are at nest
  [ FS-8-direction                                                 ;; "smell" FS-1
    fd (velocity-outside / 2 )                                     ;; move with half speed ("tandem-speed") to FS-8
    set search-time search-time - 1 ]                              ;; time they search for FS-8 counts down
 [  FS-8-direction                                                 ;; "smell" FS-8
    fd (velocity-outside / 2 )                                     ;; move with half speed ("tandem-speed") to FS-8
    set forager-energy forager-energy - metabolic-cost             ;; and with energy-costs
    set search-time search-time - 1 ]                              ;; time they search for FS-8 counts down
end

to tandem-FS-9                                                     ;; Tandem-9 heads to FS-9 together
ifelse nest?                                                       ;; if there are at nest
  [ FS-9-direction                                                 ;; "smell" FS-9
    fd (velocity-outside / 2 )                                     ;; move with half speed ("tandem-speed") to FS-9
    set search-time search-time - 1 ]                              ;; time they search for FS-9 counts down
 [  FS-9-direction                                                 ;; "smell" FS-9
    fd (velocity-outside / 2 )                                     ;; move with half speed ("tandem-speed") to FS-9
    set forager-energy forager-energy - metabolic-cost             ;; and with energy-costs
    set search-time search-time - 1 ]                              ;; time they search for FS-9 counts down
end

to tandem-FS-10                                                    ;; Tandem-10 heads to FS-10 together
ifelse nest?                                                       ;; if there are at nest
  [ FS-10-direction                                                ;; "smell" FS-10
    fd (velocity-outside / 2 )                                     ;; move with half speed ("tandem-speed") to FS-10
    set search-time search-time - 1 ]                              ;; time they search for FS-1 counts down
 [  FS-10-direction                                                ;; "smell" FS-10
    fd (velocity-outside / 2 )                                     ;; move with half speed ("tandem-speed") to FS-10
    set forager-energy forager-energy - metabolic-cost             ;; and with energy-costs
    set search-time search-time - 1 ]                              ;; time they search for FS-10 counts down
end

to FS-1-direction                                                  ;; foragers go where the strongest FS-scent-1 is
 let scent-ahead food-scent-at-angle-1   0                         ;; local variable smell ahead of forager shall be food-scent-1 (at angle 0)
  let scent-right food-scent-at-angle-1  45                        ;; local variable smell ahead of forager shall be food-scent-1 (at angle 45)
  let scent-left  food-scent-at-angle-1 -45                        ;; local variable smell ahead of forager shall be food-scent-1 (at angle -45)
  if (scent-right > scent-ahead) or (scent-left > scent-ahead)     ;; if the scent to the right is stronger or the scent to the left is stronger than the one straight forward
 [ ifelse scent-right > scent-left                                 ;; if the scent to the right is stronger than the one to the left
 [ rt 45 ]                                                         ;; the forager turns right by number degrees: If the number is negative, it turns left (follow only positive smell)
 [ lt 45 ] ]                                                       ;; else the forager turns left by number degrees: If the number is negative, it turns right (follow only positive smell)
end

to-report food-scent-at-angle-1 [angle]                            ;; reporter of the food-scent-1-variable, feedback on how strong the food scent of FS-1 is
let p patch-right-and-ahead angle 1                                ;; p is a local variable and set to: The forager "looks" 1 degrees right of its current heading
if p = nobody [ report 0 ]                                         ;; if the patch at this angle is non-existing (outside the arena), then it reports 0
report [ FS-scent-1 ] of p                                         ;; otherwise it reports the FS-scent-1 of the "foragers look"
end

to FS-2-direction                                                  ;; foragers go where the strongest FS-scent-2 is
 let scent-ahead food-scent-at-angle-2   0                         ;; local variable smell ahead of forager shall be food-scent-2 (at angle 0)
  let scent-right food-scent-at-angle-2  45                        ;; local variable smell ahead of forager shall be food-scent-2 (at angle 45)
  let scent-left  food-scent-at-angle-2 -45                        ;; local variable smell ahead of forager shall be food-scent-2 (at angle -45)
  if (scent-right > scent-ahead) or (scent-left > scent-ahead)     ;; if the scent to the right is stronger or the scent to the left is stronger than the one straight forward
 [ ifelse scent-right > scent-left                                 ;; if the scent to the right is stronger than the one to the left
 [ rt 45 ]                                                         ;; the forager turns right by number degrees: If the number is negative, it turns left (follow only positive smell)
 [ lt 45 ] ]                                                       ;; else the forager turns left by number degrees: If the number is negative, it turns right (follow only positive smell)
end

to-report food-scent-at-angle-2 [angle]                            ;; reporter of the food-scent-2-variable, feedback on how strong the food scent of FS-2 is
let p patch-right-and-ahead angle 1                                ;; p is a local variable and set to: The forager "looks" 1 degrees right of its current heading
if p = nobody [ report 0 ]                                         ;; if the patch at this angle is non-existing (outside the arena), then it reports 0
report [ FS-scent-2 ] of p                                         ;; otherwise it reports the FS-scent-2 of the "foragers look"
end

to FS-3-direction                                                     ;; foragers go where the strongest FS-scent-3 is
 let scent-ahead food-scent-at-angle-3   0                             ;; local variable smell ahead of forager shall be food-scent-3 (at angle 0)
  let scent-right food-scent-at-angle-3  45                            ;; local variable smell ahead of forager shall be food-scent-3 (at angle 45)
  let scent-left  food-scent-at-angle-3 -45                            ;; local variable smell ahead of forager shall be food-scent-3 (at angle -45)
  if (scent-right > scent-ahead) or (scent-left > scent-ahead)         ;; if the scent to the right is stronger or the scent to the left is stronger than the one straight forward
 [ ifelse scent-right > scent-left                                     ;; if the scent to the right is stronger than the one to the left
 [ rt 45 ]                                                             ;; the forager turns right by number degrees: If the number is negative, it turns left (follow only positive smell)
 [ lt 45 ] ]                                                           ;; else the forager turns left by number degrees: If the number is negative, it turns right (follow only positive smell)
end

to-report food-scent-at-angle-3 [angle]                                ;; reporter of the food-scent-3-variable, feedback on how strong the food scent of FS-3 is
let p patch-right-and-ahead angle 1                                    ;; p is a local variable and set to: The forager "looks" 1 degrees right of its current heading
if p = nobody [ report 0 ]                                             ;; if the patch at this angle is non-existing (outside the arena), then it reports 0
report [ FS-scent-3 ] of p                                             ;; otherwise it reports the FS-scent-3 of the "foragers look"
end

to FS-4-direction                                                      ;; foragers go where the strongest FS-scent-4 is
 let scent-ahead food-scent-at-angle-4   0                             ;; local variable smell ahead of forager shall be food-scent-4 (at angle 0)
  let scent-right food-scent-at-angle-4  45                            ;; local variable smell ahead of forager shall be food-scent-4 (at angle 45)
  let scent-left  food-scent-at-angle-4 -45                            ;; local variable smell ahead of forager shall be food-scent-4 (at angle -45)
  if (scent-right > scent-ahead) or (scent-left > scent-ahead)         ;; if the scent to the right is stronger or the scent to the left is stronger than the one straight forward
 [ ifelse scent-right > scent-left                                     ;; if the scent to the right is stronger than the one to the left
 [ rt 45 ]                                                             ;; the forager turns right by number degrees: If the number is negative, it turns left (follow only positive smell)
 [ lt 45 ] ]                                                           ;; else the forager turns left by number degrees: If the number is negative, it turns right (follow only positive smell)
end

to-report food-scent-at-angle-4 [angle]                                ;; reporter of the food-scent-4-variable, feedback on how strong the food scent of FS-4 is
let p patch-right-and-ahead angle 1                                    ;; p is a local variable and set to: The forager "looks" 1 degrees right of its current heading
if p = nobody [ report 0 ]                                             ;; if the patch at this angle is non-existing (outside the arena), then it reports 0
report [ FS-scent-4 ] of p                                             ;; otherwise it reports the FS-scent-4 of the "foragers look"
end

to FS-5-direction                                                      ;; foragers go where the strongest FS-scent-5 is
 let scent-ahead food-scent-at-angle-5   0                             ;; local variable smell ahead of forager shall be food-scent-5 (at angle 0)
  let scent-right food-scent-at-angle-5  45                            ;; local variable smell ahead of forager shall be food-scent-5 (at angle 45)
  let scent-left  food-scent-at-angle-5 -45                            ;; local variable smell ahead of forager shall be food-scent-5 (at angle -45)
  if (scent-right > scent-ahead) or (scent-left > scent-ahead)         ;; if the scent to the right is stronger or the scent to the left is stronger than the one straight forward
 [ ifelse scent-right > scent-left                                     ;; if the scent to the right is stronger than the one to the left
 [ rt 45 ]                                                             ;; the forager turns right by number degrees: If the number is negative, it turns left (follow only positive smell)
 [ lt 45 ] ]                                                           ;; else the forager turns left by number degrees: If the number is negative, it turns right (follow only positive smell)
end

to-report food-scent-at-angle-5 [angle]                                ;; reporter of the food-scent-5-variable, feedback on how strong the food scent of FS-5 is
let p patch-right-and-ahead angle 1                                    ;; p is a local variable and set to: The forager "looks" 1 degrees right of its current heading
if p = nobody [ report 0 ]                                             ;; if the patch at this angle is non-existing (outside the arena), then it reports 0
report [ FS-scent-5 ] of p                                             ;; otherwise it reports the FS-scent-5 of the "foragers look"
end

to FS-6-direction                                                     ;; foragers go where the strongest FS-scent-6 is
 let scent-ahead food-scent-at-angle-6   0                             ;; local variable smell ahead of forager shall be food-scent-6 (at angle 0)
  let scent-right food-scent-at-angle-6  45                            ;; local variable smell ahead of forager shall be food-scent-6 (at angle 45)
  let scent-left  food-scent-at-angle-6 -45                            ;; local variable smell ahead of forager shall be food-scent-6 (at angle -45)
  if (scent-right > scent-ahead) or (scent-left > scent-ahead)         ;; if the scent to the right is stronger or the scent to the left is stronger than the one straight forward
 [ ifelse scent-right > scent-left                                     ;; if the scent to the right is stronger than the one to the left
 [ rt 45 ]                                                             ;; the forager turns right by number degrees: If the number is negative, it turns left (follow only positive smell)
 [ lt 45 ] ]                                                           ;; else the forager turns left by number degrees: If the number is negative, it turns right (follow only positive smell)
end

to-report food-scent-at-angle-6 [angle]                                ;; reporter of the food-scent-6-variable, feedback on how strong the food scent of FS-1 is
let p patch-right-and-ahead angle 1                                    ;; p is a local variable and set to: The forager "looks" 1 degrees right of its current heading
if p = nobody [ report 0 ]                                             ;; if the patch at this angle is non-existing (outside the arena), then it reports 0
report [ FS-scent-6 ] of p                                             ;; otherwise it reports the FS-scent-6 of the "foragers look"
end

to FS-7-direction                                                      ;; foragers go where the strongest FS-scent-7 is
 let scent-ahead food-scent-at-angle-7   0                             ;; local variable smell ahead of forager shall be food-scent-7 (at angle 0)
  let scent-right food-scent-at-angle-7  45                            ;; local variable smell ahead of forager shall be food-scent-7 (at angle 45)
  let scent-left  food-scent-at-angle-7 -45                            ;; local variable smell ahead of forager shall be food-scent-7 (at angle -45)
  if (scent-right > scent-ahead) or (scent-left > scent-ahead)         ;; if the scent to the right is stronger or the scent to the left is stronger than the one straight forward
 [ ifelse scent-right > scent-left                                     ;; if the scent to the right is stronger than the one to the left
 [ rt 45 ]                                                             ;; the forager turns right by number degrees: If the number is negative, it turns left (follow only positive smell)
 [ lt 45 ] ]                                                           ;; else the forager turns left by number degrees: If the number is negative, it turns right (follow only positive smell)
end

to-report food-scent-at-angle-7 [angle]                                ;; reporter of the food-scent-7-variable, feedback on how strong the food scent of FS-1 is
let p patch-right-and-ahead angle 1                                    ;; p is a local variable and set to: The forager "looks" 1 degrees right of its current heading
if p = nobody [ report 0 ]                                             ;; if the patch at this angle is non-existing (outside the arena), then it reports 0
report [ FS-scent-7 ] of p                                             ;; otherwise it reports the FS-scent-7 of the "foragers look"
end

to FS-8-direction                                                      ;; foragers go where the strongest FS-scent-8 is
 let scent-ahead food-scent-at-angle-8   0                             ;; local variable smell ahead of forager shall be food-scent-8 (at angle 0)
  let scent-right food-scent-at-angle-8  45                            ;; local variable smell ahead of forager shall be food-scent-8 (at angle 45)
  let scent-left  food-scent-at-angle-8 -45                            ;; local variable smell ahead of forager shall be food-scent-8 (at angle -45)
  if (scent-right > scent-ahead) or (scent-left > scent-ahead)         ;; if the scent to the right is stronger or the scent to the left is stronger than the one straight forward
 [ ifelse scent-right > scent-left                                     ;; if the scent to the right is stronger than the one to the left
 [ rt 45 ]                                                             ;; the forager turns right by number degrees: If the number is negative, it turns left (follow only positive smell)
 [ lt 45 ] ]                                                           ;; else the forager turns left by number degrees: If the number is negative, it turns right (follow only positive smell)
end

to-report food-scent-at-angle-8 [angle]                                ;; reporter of the food-scent-8-variable, feedback on how strong the food scent of FS-8 is
let p patch-right-and-ahead angle 1                                    ;; p is a local variable and set to: The forager "looks" 1 degrees right of its current heading
if p = nobody [ report 0 ]                                             ;; if the patch at this angle is non-existing (outside the arena), then it reports 0
report [ FS-scent-8 ] of p                                             ;; otherwise it reports the FS-scent-8 of the "foragers look"
end

to FS-9-direction                                                      ;; foragers go where the strongest FS-scent-9 is
 let scent-ahead food-scent-at-angle-9   0                             ;; local variable smell ahead of forager shall be food-scent-9 (at angle 0)
  let scent-right food-scent-at-angle-9  45                            ;; local variable smell ahead of forager shall be food-scent-9 (at angle 45)
  let scent-left  food-scent-at-angle-9 -45                            ;; local variable smell ahead of forager shall be food-scent-9 (at angle -45)
  if (scent-right > scent-ahead) or (scent-left > scent-ahead)         ;; if the scent to the right is stronger or the scent to the left is stronger than the one straight forward
 [ ifelse scent-right > scent-left                                     ;; if the scent to the right is stronger than the one to the left
 [ rt 45 ]                                                             ;; the forager turns right by number degrees: If the number is negative, it turns left (follow only positive smell)
 [ lt 45 ] ]                                                           ;; else the forager turns left by number degrees: If the number is negative, it turns right (follow only positive smell)
end

to-report food-scent-at-angle-9 [angle]                                ;; reporter of the food-scent-9-variable, feedback on how strong the food scent of FS-9 is
let p patch-right-and-ahead angle 1                                    ;; p is a local variable and set to: The forager "looks" 1 degrees right of its current heading
if p = nobody [ report 0 ]                                             ;; if the patch at this angle is non-existing (outside the arena), then it reports 0
report [ FS-scent-9 ] of p                                             ;; otherwise it reports the FS-scent-9 of the "foragers look"
end

to FS-10-direction                                                     ;; foragers go where the strongest FS-scent-10 is
 let scent-ahead food-scent-at-angle-10   0                            ;; local variable smell ahead of forager shall be food-scent-10 (at angle 0)
  let scent-right food-scent-at-angle-10  45                           ;; local variable smell ahead of forager shall be food-scent-10 (at angle 45)
  let scent-left  food-scent-at-angle-10 -45                           ;; local variable smell ahead of forager shall be food-scent-10 (at angle -45)
  if (scent-right > scent-ahead) or (scent-left > scent-ahead)         ;; if the scent to the right is stronger or the scent to the left is stronger than the one straight forward
 [ ifelse scent-right > scent-left                                     ;; if the scent to the right is stronger than the one to the left
 [ rt 45 ]                                                             ;; the forager turns right by number degrees: If the number is negative, it turns left (follow only positive smell)
 [ lt 45 ] ]                                                           ;; else the forager turns left by number degrees: If the number is negative, it turns right (follow only positive smell)
end

to-report food-scent-at-angle-10 [angle]                               ;; reporter of the food-scent-10-variable, feedback on how strong the food scent of FS-10 is
let p patch-right-and-ahead angle 1                                    ;; p is a local variable and set to: The forager "looks" 1 degrees right of its current heading
if p = nobody [ report 0 ]                                             ;; if the patch at this angle is non-existing (outside the arena), then it reports 0
report [ FS-scent-10 ] of p                                            ;; otherwise it reports the FS-scent-10 of the "foragers look"
end

to tandem-breakup
  let breakup-prob random-float 1.000                              ;; probability is random from 0-0.999
if breakup-prob < TR-breakup-probability                           ;; default at 0.005, meanig a 5/1000 chance of becoming lost (0-0.0049 lost-TR, 0.005-1.000 remain Tandem-Follower(-1 or -2).
  [ set color sky                                                  ;; tandem follower suffering a tandem breakup are sky in color
  set forager-timer forager-timer - 1                              ;; timer for time spend searching a food source is reset by 1 for transformation
  set search-time search-time + 1                                  ;; time they search for a food source is reset by 1 for transformation
  set forager-energy forager-energy + metabolic-cost               ;; energy-costs is reset by one metabolic-cost unit for transformation
    set breed lost-TRs ]                                           ;; and lost
 end

;;;;;;;;;;;;;;;;;;;;;;;;;
;;; OTHER ACTIONS ;;;;;;;
;;;;;;;;;;;;;;;;;;;;;;;;;

to feed                                                            ;; general feeding procedure performed by drinking ants
if FS-number = 1                                                   ;; if the food source number is 1
   [ set FS-energy FS-energy - (reward-FS-1 / 120)                 ;; energy of the food source 1 is diminished by the reward of food source 1 divided by the feeding time
     set forager-energy forager-energy + (reward-FS-1 / 120)       ;; get the reward of food source 1 divided by the feeding time (incomplete feeding possible)
    set feeding-time feeding-time - 1                              ;; the feeding-time counts down
if (feeding-time = 0)                                              ;; if the feeding time is over
    [ decision-of-happiness-1 ] ]                                  ;; decide wether to be happy or unsatisfied with food source 1

if FS-number = 2                                                   ;; if the food source number is 2
   [ set FS-energy FS-energy - (reward-FS-2 / 120)                 ;; energy of the food source 2 is diminished by the reward of food source 2 divided by the feeding time
     set forager-energy forager-energy + (reward-FS-2 / 120)       ;; get the reward of food source 1 divided by the feeding time (incomplete feeding possible)
    set feeding-time feeding-time - 1                              ;; the feeding-time counts down
if (feeding-time = 0)                                              ;; if the feeding time is over
    [ decision-of-happiness-2 ] ]                                  ;; decide wether to be happy or unsatisfied with food source 2

  if FS-number = 3                                                 ;; if the food source number is 3
   [ set FS-energy FS-energy - (reward-FS-3 / 120)                 ;; energy of the food source 3 is diminished by the reward of food source 3 divided by the feeding time
     set forager-energy forager-energy + (reward-FS-3 / 120)       ;; get the reward of food source 3 divided by the feeding time (incomplete feeding possible)
    set feeding-time feeding-time - 1                              ;; the feeding-time counts down
if (feeding-time = 0)                                              ;; if the feeding time is over
    [ decision-of-happiness-3 ] ]                                  ;; decide wether to be happy or unsatisfied with food source 3

  if FS-number = 4                                                 ;; if the food source number is 4
   [ set FS-energy FS-energy - (reward-FS-4 / 120)                 ;; energy of the food source 4 is diminished by the reward of food source 4 divided by the feeding time
     set forager-energy forager-energy + (reward-FS-4 / 120)       ;; get the reward of food source 4 divided by the feeding time (incomplete feeding possible)
    set feeding-time feeding-time - 1                              ;; the feeding-time counts down
if (feeding-time = 0)                                              ;; if the feeding time is over
    [ decision-of-happiness-4 ] ]                                  ;; decide wether to be happy or unsatisfied with food source 4

  if FS-number = 5                                                 ;; if the food source number is 5
   [ set FS-energy FS-energy - (reward-FS-5 / 120)                 ;; energy of the food source 5 is diminished by the reward of food source 5 divided by the feeding time
     set forager-energy forager-energy + (reward-FS-5 / 120)       ;; get the reward of food source 5 divided by the feeding time (incomplete feeding possible)
    set feeding-time feeding-time - 1                              ;; the feeding-time counts down
if (feeding-time = 0)                                              ;; if the feeding time is over
    [ decision-of-happiness-5 ] ]                                  ;; decide wether to be happy or unsatisfied with food source 5

  if FS-number = 6                                                 ;; if the food source number is 6
   [ set FS-energy FS-energy - (reward-FS-6 / 120)                 ;; energy of the food source 1 is diminished by the reward of food source 6 divided by the feeding time
     set forager-energy forager-energy + (reward-FS-6 / 120)       ;; get the reward of food source 6 divided by the feeding time (incomplete feeding possible)
    set feeding-time feeding-time - 1                              ;; the feeding-time counts down
if (feeding-time = 0)                                              ;; if the feeding time is over
    [ decision-of-happiness-6 ] ]                                  ;; decide wether to be happy or unsatisfied with food source 6

  if FS-number = 7                                                 ;; if the food source number is 7
   [ set FS-energy FS-energy - (reward-FS-7 / 120)                 ;; energy of the food source 7 is diminished by the reward of food source 7 divided by the feeding time
     set forager-energy forager-energy + (reward-FS-7 / 120)       ;; get the reward of food source 7 divided by the feeding time (incomplete feeding possible)
    set feeding-time feeding-time - 1                              ;; the feeding-time counts down
if (feeding-time = 0)                                              ;; if the feeding time is over
    [ decision-of-happiness-7 ] ]                                  ;; decide wether to be happy or unsatisfied with food source 7

  if FS-number = 8                                                 ;; if the food source number is 8
   [ set FS-energy FS-energy - (reward-FS-8 / 120)                 ;; energy of the food source 8 is diminished by the reward of food source 8 divided by the feeding time
     set forager-energy forager-energy + (reward-FS-8 / 120)       ;; get the reward of food source 8 divided by the feeding time (incomplete feeding possible)
    set feeding-time feeding-time - 1                              ;; the feeding-time counts down
if (feeding-time = 0)                                              ;; if the feeding time is over
    [ decision-of-happiness-8 ] ]                                  ;; decide wether to be happy or unsatisfied with food source 8

  if FS-number = 9                                                 ;; if the food source number is 9
   [ set FS-energy FS-energy - (reward-FS-9 / 120)                 ;; energy of the food source 9 is diminished by the reward of food source 9 divided by the feeding time
     set forager-energy forager-energy + (reward-FS-9 / 120)       ;; get the reward of food source 9 divided by the feeding time (incomplete feeding possible)
    set feeding-time feeding-time - 1                              ;; the feeding-time counts down
if (feeding-time = 0)                                              ;; if the feeding time is over
    [ decision-of-happiness-9 ] ]                                  ;; decide wether to be happy or unsatisfied with food source 9

    if FS-number = 10                                              ;; if the food source number is 10
   [ set FS-energy FS-energy - (reward-FS-10 / 120)                ;; energy of the food source 10 is diminished by the reward of food source 10 divided by the feeding time
     set forager-energy forager-energy + (reward-FS-10 / 120)      ;; get the reward of food source 10 divided by the feeding time (incomplete feeding possible)
    set feeding-time feeding-time - 1                              ;; the feeding-time counts down
if (feeding-time = 0)                                              ;; if the feeding time is over
    [ decision-of-happiness-10 ] ]                                 ;; decide wether to be happy or unsatisfied with food source 10
end

to unload                                                          ;; procedure at the nest: transfer forager energy to nest
  set nest-energy nest-energy + forager-energy                     ;; turnover energy (positive and negative values possible)
  set forager-energy 0                                             ;; reset forager-energy to 0 (all energy has been transferred to nest)
end

to reset-times                                                     ;; reset internal timers of the agents
  set feeding-time 120                                             ;; feeding time is default 120 ticks
  set scouts-time 0                                                ;; scouting time is default 0
  set nest-stay-time unloading-time                                ;; nest stay time is the adjustable unloading time, default 60 ticks
  set search-time FS-searching-time                                ;; search time for a food source is default 180 ticks
  set forager-timer 0                                              ;; forager time is reset to 0
end

;;;;;;;;;;;;;;;;;;;;;;;;
;;; TANDEM-FORMATION ;;;
;;;;;;;;;;;;;;;;;;;;;;;;

to Tandemformation-1                                               ;; tandem to food source 1 forms
  let Follower one-of recruits-here                                ;; Tandemfollower-1 shall be a random recruit
if Follower != nobody                                              ;; got a recruit?
[ ask Tandemstarters-1-here                                        ;; if a Tandemstarter-1 encounters a recruit on the same patch
   [ set breed Tandemleaders-1                                     ;; and he becomes a Tandemleader-1
      reset-times ]                                                ;; reset internal timers
ask Follower                                                       ;; the recruit becomes a Follower
[ output-show forager-timer                                        ;; and triggers the timer to display their time spend waiting to be recruited
  set breed Tandemfollowers-1                                      ;; become Tandem-Followers to FS-1
  reset-times ] ]                                                  ;; reset internal timers
end

to Tandemformation-2                                               ;; tandem to food source 2 forms
  let Follower one-of recruits-here                                ;; Tandemfollower-2 shall be a random recruit
if Follower != nobody                                              ;; got a recruit?
[ ask Tandemstarters-2-here                                        ;; if a Tandemstarter-2 encounters a recruit on the same patch
   [ set breed Tandemleaders-2                                     ;; and he becomes a Tandemleader-2
      reset-times ]                                                ;; reset internal timers
ask Follower                                                       ;; the recruit becomes a Follower
[ output-show forager-timer                                        ;; and triggers the timer to display their time spend waiting to be recruited
  set breed Tandemfollowers-2                                      ;; become Tandem-Followers to FS-2
  reset-times ] ]                                                  ;; reset internal timers
end

to Tandemformation-3                                               ;; tandem to food source 3 forms
  let Follower one-of recruits-here                                ;; Tandemfollower-3 shall be a random recruit
if Follower != nobody                                              ;; got a recruit?
[ ask Tandemstarters-3-here                                        ;; if a Tandemstarter-3 encounters a recruit on the same patch
   [ set breed Tandemleaders-3                                     ;; and he becomes a Tandemleader-3
      reset-times ]                                                ;; reset internal timers
ask Follower                                                       ;; the recruit becomes a Follower
[ output-show forager-timer                                        ;; and triggers the timer to display their time spend waiting to be recruited
  set breed Tandemfollowers-3                                      ;; become Tandem-Followers to FS-3
  reset-times ] ]                                                  ;; reset internal timers
end

to Tandemformation-4                                               ;; tandem to food source 4 forms
  let Follower one-of recruits-here                                ;; Tandemfollower-4 shall be a random recruit
if Follower != nobody                                              ;; got a recruit?
[ ask Tandemstarters-4-here                                        ;; if a Tandemstarter-4 encounters a recruit on the same patch
   [ set breed Tandemleaders-4                                     ;; and he becomes a Tandemleader-4
      reset-times ]                                                ;; reset internal timers
ask Follower                                                       ;; the recruit becomes a Follower
[ output-show forager-timer                                        ;; and triggers the timer to display their time spend waiting to be recruited
  set breed Tandemfollowers-4                                      ;; become Tandem-Followers to FS-4
  reset-times ] ]                                                  ;; reset internal timers
end

to Tandemformation-5                                               ;; tandem to food source 5 forms
  let Follower one-of recruits-here                                ;; Tandemfollower-5 shall be a random recruit
if Follower != nobody                                              ;; got a recruit?
[ ask Tandemstarters-5-here                                        ;; if a Tandemstarter-5 encounters a recruit on the same patch
   [ set breed Tandemleaders-5                                     ;; and he becomes a Tandemleader-5
      reset-times ]                                                ;; reset internal timers
ask Follower                                                       ;; the recruit becomes a Follower
[ output-show forager-timer                                        ;; and triggers the timer to display their time spend waiting to be recruited
  set breed Tandemfollowers-5                                      ;; become Tandem-Followers to FS-5
  reset-times ] ]                                                  ;; reset internal timers
end

to Tandemformation-6                                               ;; tandem to food source 6 forms
  let Follower one-of recruits-here                                ;; Tandemfollower-6 shall be a random recruit
if Follower != nobody                                              ;; got a recruit?
[ ask Tandemstarters-6-here                                        ;; if a Tandemstarter-6 encounters a recruit on the same patch
   [ set breed Tandemleaders-6                                     ;; and he becomes a Tandemleader-6
      reset-times ]                                                ;; reset internal timers
ask Follower                                                       ;; the recruit becomes a Follower
[ output-show forager-timer                                        ;; and triggers the timer to display their time spend waiting to be recruited
  set breed Tandemfollowers-6                                      ;; become Tandem-Followers to FS-6
  reset-times ] ]                                                  ;; reset internal timers
end

to Tandemformation-7                                               ;; tandem to food source 7 forms
  let Follower one-of recruits-here                                ;; Tandemfollower-7 shall be a random recruit
if Follower != nobody                                              ;; got a recruit?
[ ask Tandemstarters-7-here                                        ;; if a Tandemstarter-7 encounters a recruit on the same patch
   [ set breed Tandemleaders-7                                     ;; and he becomes a Tandemleader-7
      reset-times ]                                                ;; reset internal timers
ask Follower                                                       ;; the recruit becomes a Follower
[ output-show forager-timer                                        ;; and triggers the timer to display their time spend waiting to be recruited
  set breed Tandemfollowers-7                                      ;; become Tandem-Followers to FS-7
  reset-times ] ]                                                  ;; reset internal timers
end

to Tandemformation-8                                               ;; tandem to food source 8 forms
  let Follower one-of recruits-here                                ;; Tandemfollower-8 shall be a random recruit
if Follower != nobody                                              ;; got a recruit?
[ ask Tandemstarters-8-here                                        ;; if a Tandemstarter-8 encounters a recruit on the same patch
   [ set breed Tandemleaders-8                                     ;; and he becomes a Tandemleader-8
      reset-times ]                                                ;; reset internal timers
ask Follower                                                       ;; the recruit becomes a Follower
[ output-show forager-timer                                        ;; and triggers the timer to display their time spend waiting to be recruited
  set breed Tandemfollowers-8                                      ;; become Tandem-Followers to FS-8
  reset-times ] ]                                                  ;; reset internal timers
end

to Tandemformation-9                                               ;; tandem to food source 9 forms
  let Follower one-of recruits-here                                ;; Tandemfollower-9 shall be a random recruit
if Follower != nobody                                              ;; got a recruit?
[ ask Tandemstarters-9-here                                        ;; if a Tandemstarter-9 encounters a recruit on the same patch
   [ set breed Tandemleaders-9                                     ;; and he becomes a Tandemleader-9
      reset-times ]                                                ;; reset internal timers
ask Follower                                                       ;; the recruit becomes a Follower
[ output-show forager-timer                                        ;; and triggers the timer to display their time spend waiting to be recruited
  set breed Tandemfollowers-9                                      ;; become Tandem-Followers to FS-9
  reset-times ] ]                                                  ;; reset internal timers
end

to Tandemformation-10                                              ;; tandem to food source 10 forms
  let Follower one-of recruits-here                                ;; Tandemfollower-10 shall be a random recruit
if Follower != nobody                                              ;; got a recruit?
[ ask Tandemstarters-10-here                                       ;; if a Tandemstarter-1 encounters a recruit on the same patch
   [ set breed Tandemleaders-10                                    ;; and he becomes a Tandemleader-10
      reset-times ]                                                ;; reset internal timers
ask Follower                                                       ;; the recruit becomes a Follower
[ output-show forager-timer                                        ;; and triggers the timer to display their time spend waiting to be recruited
  set breed Tandemfollowers-10                                     ;; become Tandem-Followers to FS-10
  reset-times ] ]                                                  ;; reset internal timers
end

;;;;;;;;;;;;;;;;;
;;; DECISIONS ;;;
;;;;;;;;;;;;;;;;;

to decision-of-foraging                                            ;; decision wether to go on foraging
  ifelse abs pxcor >= 35 or abs pycor >= 35                        ;; if to far out (half arena size perimeter)
  [ set breed returning-scouts ]                                   ;; return to nest
    [ set scouts-time 300 ]                                        ;; otherwise get an additionally 300 ticks scouting time
end

to decision-of-happiness-1                                         ;; decision wether to be happy with FS-1 or not
  reset-times                                                      ;; reset internal timers of agents
  ifelse  FS-quality  > 0                                          ;; if the quality of food source 1 is high
  [ set happiness 100 ]                                            ;; be 100% satisfied with it and become happy-sucessful-foragers-1
  [ set happiness random 100 ]                                     ;; else the FS-quality is low; the foragers are randomly happy with it
ifelse happiness >= happiness-threshold                            ;; if their happiness is same or higher than the adjustable happiness-threshold (default at 90 %, meaning only 10 % of ants will be happy with a low quality FS-1).
  [ set breed happy-successful-foragers-1 ]                        ;; be happy with FS-1 and become happy-successful-foragers-1 (90-99: happy)
  [ set breed unsatisfied-foragers ]                               ;; be unhappy with FS-1 and become unsatisfied-foragers (0-89: unhappy)
end

to decision-of-happiness-2                                         ;; decision wether to be happy with FS-2 or not
  reset-times                                                      ;; reset internal timers
  ifelse  FS-quality  > 0                                          ;; if the quality of food source 2 is high
  [ set happiness 100 ]                                            ;; be 100% satisfied with it and become happy-sucessful-foragers-2
  [ set happiness random 100 ]                                     ;; else the FS-quality is low; the foragers are randomly happy with it
ifelse happiness >= happiness-threshold                            ;; if their happiness is same or higher than the adjustable happiness-threshold (default at 90 %, meaning only 10 % of ants will be happy with a low quality FS-2).
  [ set breed happy-successful-foragers-2 ]                        ;; be happy with FS-2 and become happy-successful-foragers-2 (90-99: happy)
  [ set breed unsatisfied-foragers ]                               ;; be unhappy with FS-2 and become unsatisfied-foragers (0-89: unhappy)
end

to decision-of-happiness-3                                         ;; decision wether to be happy with FS-3 or not
  reset-times                                                      ;; reset internal timers of agents
  ifelse  FS-quality  > 0                                          ;; if the quality of food source 3 is high
  [ set happiness 100 ]                                            ;; be 100% satisfied with it and become happy-sucessful-foragers-3
  [ set happiness random 100 ]                                     ;; else the FS-quality is low; the foragers are randomly happy with it
ifelse happiness >= happiness-threshold                            ;; if their happiness is same or higher than the adjustable happiness-threshold (default at 90 %, meaning only 10 % of ants will be happy with a low quality FS-1).
  [ set breed happy-successful-foragers-3 ]                        ;; be happy with FS-3 and become happy-successful-foragers-3 (90-99: happy)
  [ set breed unsatisfied-foragers ]                               ;; be unhappy with FS-3 and become unsatisfied-foragers (0-89: unhappy)
end

to decision-of-happiness-4                                         ;; decision wether to be happy with FS-4 or not
  reset-times                                                      ;; reset internal timers of agents
  ifelse  FS-quality  > 0                                          ;; if the quality of food source 4 is high
  [ set happiness 100 ]                                            ;; be 100% satisfied with it and become happy-sucessful-foragers-4
  [ set happiness random 100 ]                                     ;; else the FS-quality is low; the foragers are randomly happy with it
ifelse happiness >= happiness-threshold                            ;; if their happiness is same or higher than the adjustable happiness-threshold (default at 90 %, meaning only 10 % of ants will be happy with a low quality FS-1).
  [ set breed happy-successful-foragers-4 ]                        ;; be happy with FS-4 and become happy-successful-foragers-4 (90-99: happy)
  [ set breed unsatisfied-foragers ]                               ;; be unhappy with FS-4 and become unsatisfied-foragers (0-89: unhappy)
end

to decision-of-happiness-5                                         ;; decision wether to be happy with FS-5 or not
  reset-times                                                      ;; reset internal timers of agents
  ifelse  FS-quality  > 0                                          ;; if the quality of food source 5 is high
  [ set happiness 100 ]                                            ;; be 100% satisfied with it and become happy-sucessful-foragers-5
  [ set happiness random 100 ]                                     ;; else the FS-quality is low; the foragers are randomly happy with it
ifelse happiness >= happiness-threshold                            ;; if their happiness is same or higher than the adjustable happiness-threshold (default at 90 %, meaning only 10 % of ants will be happy with a low quality FS-1).
  [ set breed happy-successful-foragers-5 ]                        ;; be happy with FS-5 and become happy-successful-foragers-5 (90-99: happy)
  [ set breed unsatisfied-foragers ]                               ;; be unhappy with FS-5 and become unsatisfied-foragers (0-89: unhappy)
end

to decision-of-happiness-6                                         ;; decision wether to be happy with FS-6 or not
  reset-times                                                      ;; reset internal timers of agents
  ifelse  FS-quality  > 0                                          ;; if the quality of food source 6 is high
  [ set happiness 100 ]                                            ;; be 100% satisfied with it and become happy-sucessful-foragers-6
  [ set happiness random 100 ]                                     ;; else the FS-quality is low; the foragers are randomly happy with it
ifelse happiness >= happiness-threshold                            ;; if their happiness is same or higher than the adjustable happiness-threshold (default at 90 %, meaning only 10 % of ants will be happy with a low quality FS-1).
  [ set breed happy-successful-foragers-6 ]                        ;; be happy with FS-6 and become happy-successful-foragers-6 (90-99: happy)
  [ set breed unsatisfied-foragers ]                               ;; be unhappy with FS-6 and become unsatisfied-foragers (0-89: unhappy)
end

to decision-of-happiness-7                                         ;; decision wether to be happy with FS-7 or not
  reset-times                                                      ;; reset internal timers of agents
  ifelse  FS-quality  > 0                                          ;; if the quality of food source 7 is high
  [ set happiness 100 ]                                            ;; be 100% satisfied with it and become happy-sucessful-foragers-7
  [ set happiness random 100 ]                                     ;; else the FS-quality is low; the foragers are randomly happy with it
ifelse happiness >= happiness-threshold                            ;; if their happiness is same or higher than the adjustable happiness-threshold (default at 90 %, meaning only 10 % of ants will be happy with a low quality FS-1).
  [ set breed happy-successful-foragers-7 ]                        ;; be happy with FS-7 and become happy-successful-foragers-7 (90-99: happy)
  [ set breed unsatisfied-foragers ]                               ;; be unhappy with FS-7 and become unsatisfied-foragers (0-89: unhappy)
end

to decision-of-happiness-8                                         ;; decision wether to be happy with FS-8 or not
  reset-times                                                      ;; reset internal timers of agents
  ifelse  FS-quality  > 0                                          ;; if the quality of food source 8 is high
  [ set happiness 100 ]                                            ;; be 100% satisfied with it and become happy-sucessful-foragers-8
  [ set happiness random 100 ]                                     ;; else the FS-quality is low; the foragers are randomly happy with it
ifelse happiness >= happiness-threshold                            ;; if their happiness is same or higher than the adjustable happiness-threshold (default at 90 %, meaning only 10 % of ants will be happy with a low quality FS-1).
  [ set breed happy-successful-foragers-8 ]                        ;; be happy with FS-8 and become happy-successful-foragers-8 (90-99: happy)
  [ set breed unsatisfied-foragers ]                               ;; be unhappy with FS-8 and become unsatisfied-foragers (0-89: unhappy)
end

to decision-of-happiness-9                                         ;; decision wether to be happy with FS-9 or not
  reset-times                                                      ;; reset internal timers of agents
  ifelse  FS-quality  > 0                                          ;; if the quality of food source 9 is high
  [ set happiness 100 ]                                            ;; be 100% satisfied with it and become happy-sucessful-foragers-9
  [ set happiness random 100 ]                                     ;; else the FS-quality is low; the foragers are randomly happy with it
ifelse happiness >= happiness-threshold                            ;; if their happiness is same or higher than the adjustable happiness-threshold (default at 90 %, meaning only 10 % of ants will be happy with a low quality FS-1).
  [ set breed happy-successful-foragers-9 ]                        ;; be happy with FS-9 and become happy-successful-foragers-9 (90-99: happy)
  [ set breed unsatisfied-foragers ]                               ;; be unhappy with FS-9 and become unsatisfied-foragers (0-89: unhappy)
end

to decision-of-happiness-10                                        ;; decision wether to be happy with FS-10 or not
  reset-times                                                      ;; reset internal timers of agents
  ifelse  FS-quality  > 0                                          ;; if the quality of food source 10 is high
  [ set happiness 100 ]                                            ;; be 100% satisfied with it and become happy-sucessful-foragers-10
  [ set happiness random 100 ]                                     ;; else the FS-quality is low; the foragers are randomly happy with it
ifelse happiness >= happiness-threshold                            ;; if their happiness is same or higher than the adjustable happiness-threshold (default at 90 %, meaning only 10 % of ants will be happy with a low quality FS-1).
  [ set breed happy-successful-foragers-10 ]                       ;; be happy with FS-10 and become happy-successful-foragers-10 (90-99: happy)
  [ set breed unsatisfied-foragers ]                               ;; be unhappy with FS-10 and become unsatisfied-foragers (0-89: unhappy)
end

to decision-of-recruitment-1                                       ;; decision if recruitment to FS-1 happens
  reset-times                                                      ;; reset internal timers
  let recruitprob random 100                                       ;; probability to recruit is random from 0-99
ifelse recruitment-probability-1 > recruitprob                     ;; if the recruitment-probability-1 (default 50) is higher than random value
  [ set nest-stay-time 120                                         ;; their nest-stay-time is set to 120 and
    set breed Tandemstarters-1 ]                                   ;; recruitment will iniate.
  [ set breed successful-foragers-1 ]                              ;; else: head to FS-1 alone with private information
end

to decision-of-recruitment-2                                       ;; decision if recruitment to FS-2 happens
  reset-times                                                      ;; reset internal timers
  let recruitprob random 100                                       ;; probability to recruit is random from 0-99
ifelse recruitment-probability-2 > recruitprob                     ;; if the recruitment-probability-2 (default 50) is higher than random value
  [ set nest-stay-time 120                                         ;; their nest-stay-time is set to 120 and
    set breed Tandemstarters-2 ]                                   ;; recruitment will iniate.
  [ set breed successful-foragers-2 ]                              ;; head to FS-2 alone with private information
end

to decision-of-recruitment-3                                       ;; decision if recruitment to FS-3 happens
  reset-times                                                      ;; reset internal timers
  let recruitprob random 100                                       ;; probability to recruit is random from 0-99
ifelse recruitment-probability-3 > recruitprob                     ;; if the recruitment-probability-3 (default 50) is higher than random value
  [ set nest-stay-time 120                                         ;; their nest-stay-time is set to 120 and
    set breed Tandemstarters-3 ]                                   ;; recruitment will iniate.
  [ set breed successful-foragers-3 ]                              ;; else: head to FS-3 alone with private information
end

to decision-of-recruitment-4                                       ;; decision if recruitment to FS-4 happens
  reset-times                                                      ;; reset internal timers
  let recruitprob random 100                                       ;; probability to recruit is random from 0-99
ifelse recruitment-probability-4 > recruitprob                     ;; if the recruitment-probability-4 (default 50) is higher than random value
  [ set nest-stay-time 120                                         ;; their nest-stay-time is set to 120 and
    set breed Tandemstarters-4 ]                                   ;; recruitment will iniate.
  [ set breed successful-foragers-4 ]                              ;; else: head to FS-4 alone with private information
end

to decision-of-recruitment-5                                       ;; decision if recruitment to FS-5 happens
  reset-times                                                      ;; reset internal timers
  let recruitprob random 100                                       ;; probability to recruit is random from 0-99
ifelse recruitment-probability-5 > recruitprob                     ;; if the recruitment-probability-5 (default 50) is higher than random value
  [ set nest-stay-time 120                                         ;; their nest-stay-time is set to 120 and
    set breed Tandemstarters-5 ]                                   ;; recruitment will iniate.
  [ set breed successful-foragers-5 ]                              ;; else: head to FS-5 alone with private information
end

to decision-of-recruitment-6                                       ;; decision if recruitment to FS-6 happens
  reset-times                                                      ;; reset internal timers
  let recruitprob random 100                                       ;; probability to recruit is random from 0-99
ifelse recruitment-probability-6 > recruitprob                     ;; if the recruitment-probability-6 (default 50) is higher than random value
  [ set nest-stay-time 120                                         ;; their nest-stay-time is set to 120 and
    set breed Tandemstarters-6 ]                                   ;; recruitment will iniate.
  [ set breed successful-foragers-6 ]                              ;; else: head to FS-6 alone with private information
end

to decision-of-recruitment-7                                       ;; decision if recruitment to FS-7 happens
  reset-times                                                      ;; reset internal timers
  let recruitprob random 100                                       ;; probability to recruit is random from 0-99
ifelse recruitment-probability-7 > recruitprob                     ;; if the recruitment-probability-7 (default 50) is higher than random value
  [ set nest-stay-time 120                                         ;; their nest-stay-time is set to 120 and
    set breed Tandemstarters-7 ]                                   ;; recruitment will iniate.
  [ set breed successful-foragers-7 ]                              ;; else: head to FS-7 alone with private information
end

to decision-of-recruitment-8                                       ;; decision if recruitment to FS-8 happens
  reset-times                                                      ;; reset internal timers
  let recruitprob random 100                                       ;; probability to recruit is random from 0-99
ifelse recruitment-probability-8 > recruitprob                     ;; if the recruitment-probability-8 (default 50) is higher than random value
  [ set nest-stay-time 120                                         ;; their nest-stay-time is set to 120 and
    set breed Tandemstarters-8 ]                                   ;; recruitment will iniate.
  [ set breed successful-foragers-8 ]                              ;; else: head to FS-8 alone with private information
end

to decision-of-recruitment-9                                       ;; decision if recruitment to FS-9 happens
  reset-times                                                      ;; reset internal timers
  let recruitprob random 100                                       ;; probability to recruit is random from 0-99
ifelse recruitment-probability-9 > recruitprob                     ;; if the recruitment-probability-9 (default 50) is higher than random value
  [ set nest-stay-time 120                                         ;; their nest-stay-time is set to 120 and
    set breed Tandemstarters-9 ]                                   ;; recruitment will iniate.
  [ set breed successful-foragers-9 ]                              ;; else: head to FS-9 alone with private information
end

to decision-of-recruitment-10                                      ;; decision if recruitment to FS-1 happens
  reset-times                                                      ;; reset internal timers
  let recruitprob random 100                                       ;; probability to recruit is random from 0-99
ifelse recruitment-probability-10 > recruitprob                    ;; if the recruitment-probability-1 (default 50) is higher than random value
  [ set nest-stay-time 120                                         ;; their nest-stay-time is set to 120 and
    set breed Tandemstarters-10 ]                                  ;; recruitment will iniate.
  [ set breed successful-foragers-10 ]                             ;; else: head to FS-1 alone with private information
end

to decision-of-foraging-strategy                                   ;; decision of unsatisfied foragers  whether to become scouts or recruits when their time in nest is over
  reset-times                                                      ;; reset internal timers
  let strategyprob random 100                                      ;; probability to become a scout or recruit is random from 0-99
ifelse strategyprob < 50                                           ;; chance of becoming a scout or recruit is equal! (0-49: scout, 50-99 recruit)
  [ set breed scouts                                               ;; become scouts
    set color red ]                                                ;; and are red
  [ set breed recruits                                             ;; become recruits
    set color grey ]                                               ;; and are grey
end

to decision-of-lost-TRS                                            ;; decision of lost-TRs wether to become a scout or return to the nest when they failed to find a FS in the search time
  let strategyprob random 100                                      ;; probability to become a scout or return to nest is random from 0-99.
ifelse strategyprob < 50                                           ;; chance of scouting or return to nest is equal (0-49: scouts, 50-99 return to nest)
  [ set breed scouts                                               ;; become scouts
    set color red                                                  ;; and are red
      reset-times ]                                                ;; reset internal timers
  [ set breed unsatisfied-foragers ]                               ;; else: become an unsatisfied forager and return to nest
end

;;;;;;;;;;;;;;;;;;
;;; FS-ACTIONS ;;;
;;;;;;;;;;;;;;;;;;

to FS-Disappearence                                                ;; food sources vanishes when their energy is depleted
 ask patches with [ FS-number = 1 ]                                ;; all patches with food source number 1
 [ if FS-energy < 0.00001                                          ;; if the food source energy is too low to feed on it
 [ set if-food? false                                              ;; become a "normal" patch again
   set  FS-quality  0                                              ;; have 0 quality
   set pcolor black                                                ;; black color
   set FS-number 0 ] ]                                             ;; and no food source number
 ask patches with [ FS-number = 2 ]                                ;; all patches with food source number 2
 [ if FS-energy < 0.00001                                          ;; if the food source energy is too low to feed on it
 [ set if-food? false                                              ;; become a "normal" patch again
   set  FS-quality  0                                              ;; have 0 quality
   set pcolor black                                                ;; black color
   set FS-number 0 ] ]                                             ;; and no food source number
   ask patches with [ FS-number = 3 ]                              ;; all patches with food source number 3
 [ if FS-energy < 0.00001                                          ;; if the food source energy is too low to feed on it
 [ set if-food? false                                              ;; become a "normal" patch again
   set  FS-quality  0                                              ;; have 0 quality
   set pcolor black                                                ;; black color
   set FS-number 0 ] ]                                             ;; and no food source number
   ask patches with [ FS-number = 4 ]                              ;; all patches with food source number 4
 [ if FS-energy < 0.00001                                          ;; if the food source energy is too low to feed on it
 [ set if-food? false                                              ;; become a "normal" patch again
   set  FS-quality  0                                              ;; have 0 quality
   set pcolor black                                                ;; black color
   set FS-number 0 ] ]                                             ;; and no food source number
   ask patches with [ FS-number = 5 ]                              ;; all patches with food source number 5
 [ if FS-energy < 0.00001                                          ;; if the food source energy is too low to feed on it
 [ set if-food? false                                              ;; become a "normal" patch again
   set  FS-quality  0                                              ;; have 0 quality
   set pcolor black                                                ;; black color
   set FS-number 0 ] ]                                             ;; and no food source number
   ask patches with [ FS-number = 6 ]                              ;; all patches with food source number 6
 [ if FS-energy < 0.00001                                          ;; if the food source energy is too low to feed on it
 [ set if-food? false                                              ;; become a "normal" patch again
   set  FS-quality  0                                              ;; have 0 quality
   set pcolor black                                                ;; black color
   set FS-number 0 ] ]                                             ;; and no food source number
   ask patches with [ FS-number = 7 ]                              ;; all patches with food source number 7
 [ if FS-energy < 0.00001                                          ;; if the food source energy is too low to feed on it
 [ set if-food? false                                              ;; become a "normal" patch again
   set  FS-quality  0                                              ;; have 0 quality
   set pcolor black                                                ;; black color
   set FS-number 0 ] ]                                             ;; and no food source number
   ask patches with [ FS-number = 8 ]                              ;; all patches with food source number 8
 [ if FS-energy < 0.00001                                          ;; if the food source energy is too low to feed on it
 [ set if-food? false                                              ;; become a "normal" patch again
   set  FS-quality  0                                              ;; have 0 quality
   set pcolor black                                                ;; black color
   set FS-number 0 ] ]                                             ;; and no food source number
   ask patches with [ FS-number = 9 ]                              ;; all patches with food source number 9
 [ if FS-energy < 0.00001                                          ;; if the food source energy is too low to feed on it
 [ set if-food? false                                              ;; become a "normal" patch again
   set  FS-quality  0                                              ;; have 0 quality
   set pcolor black                                                ;; black color
   set FS-number 0 ] ]                                             ;; and no food source number
   ask patches with [ FS-number = 10 ]                             ;; all patches with food source number 10
 [ if FS-energy < 0.00001                                          ;; if the food source energy is too low to feed on it
 [ set if-food? false                                              ;; become a "normal" patch again
   set  FS-quality  0                                              ;; have 0 quality
   set pcolor black                                                ;; black color
   set FS-number 0 ] ]                                             ;; and no food source number
end

to FS-Renewal-1                                                    ;; food source 1 can reappear
    ask patch 0 51                                                 ;; at the same location as the former food source 1
 [ set pcolor white                                                ;; the food source center shall be white of color
   if (distancexy 0 51 ) < Food-size                               ;; the new food source the same size as the old food sources
 [ set FS-number 1                                                 ;; give it the number 1
   set if-food? true                                               ;; let it have food
   set FS-scent-1 200 - distancexy 0 51                            ;; and a food scent
ifelse reward-FS-1 >= 0.75                                         ;; if the depleted FS had high quality food,
 [ set  FS-quality  1                                              ;; the new FS also has a high quality
   set FS-energy food-energy-high ]                                ;; and high energy
 [ set  FS-quality  0                                              ;; else it has a low quality
   set FS-energy food-energy-low] ] ]                              ;; and a low energy
end

to FS-Renewal-2                                                    ;; food source 2 can reappear
    ask patch 30 39                                                ;; at the same location as the former food source 2
 [ set pcolor white                                                ;; the food source center shall be white of color
   if (distancexy 30 39 ) < Food-size                              ;; the new food source the same size as the old food sources
 [ set FS-number 2                                                 ;; give it the number 2
   set if-food? true                                               ;; let it have food
   set FS-scent-2 200 - distancexy 30 39                           ;; and a food scent
ifelse reward-FS-2 >= 0.75                                         ;; if the depleted FS had high quality food,
 [ set  FS-quality  1                                              ;; the new FS also has a high quality
   set FS-energy food-energy-high ]                                ;; and high energy
 [ set  FS-quality  0                                              ;; else it has a low quality
   set FS-energy food-energy-low] ] ]                              ;; and a low energy
end

to FS-Renewal-3                                                    ;; food source 3 can reappear
    ask patch 48 15                                                ;; at the same location as the former food source 3
 [ set pcolor white                                                ;; the food source center shall be white of color
   if (distancexy 48 15 ) < Food-size                              ;; the new food source the same size as the old food sources
 [ set FS-number 3                                                 ;; give it the number 3
   set if-food? true                                               ;; let it have food
   set FS-scent-3 200 - distancexy 48 15                           ;; and a food scent
ifelse reward-FS-3 >= 0.75                                         ;; if the depleted FS had high quality food,
 [ set  FS-quality  1                                              ;; the new FS also has a high quality
   set FS-energy food-energy-high ]                                ;; and high energy
 [ set  FS-quality  0                                              ;; else it has a low quality
   set FS-energy food-energy-low] ] ]                              ;; and a low energy
end

to FS-Renewal-4                                                    ;; food source 4 can reappear
    ask patch 48 -17                                               ;; at the same location as the former food source 4
 [ set pcolor white                                                ;; the food source center shall be white of color
   if (distancexy 48 -17 ) < Food-size                             ;; the new food source the same size as the old food sources
 [ set FS-number 4                                                 ;; give it the number 4
   set if-food? true                                               ;; let it have food
   set FS-scent-4 200 - distancexy 48 -17                          ;; and a food scent
ifelse reward-FS-4 >= 0.75                                         ;; if the depleted FS had high quality food,
 [ set  FS-quality  1                                              ;; the new FS also has a high quality
   set FS-energy food-energy-high ]                                ;; and high energy
 [ set  FS-quality  0                                              ;; else it has a low quality
   set FS-energy food-energy-low] ] ]                              ;; and a low energy
end

to FS-Renewal-5                                                    ;; food source 5 can reappear
    ask patch 30 -39                                               ;; at the same location as the former food source 5
 [ set pcolor white                                                ;; the food source center shall be white of color
   if (distancexy 30 -39 ) < Food-size                             ;; the new food source the same size as the old food sources
 [ set FS-number 5                                                 ;; give it the number 5
   set if-food? true                                               ;; let it have food
   set FS-scent-5 200 - distancexy 30 -39                          ;; and a food scent
ifelse reward-FS-5 >= 0.75                                         ;; if the depleted FS had high quality food,
 [ set  FS-quality  1                                              ;; the new FS also has a high quality
   set FS-energy food-energy-high ]                                ;; and high energy
 [ set  FS-quality  0                                              ;; else it has a low quality
   set FS-energy food-energy-low] ] ]                              ;; and a low energy
end

to FS-Renewal-6                                                    ;; food source 6 can reappear
    ask patch 0 -51                                                ;; at the same location as the former food source 6
 [ set pcolor white                                                ;; the food source center shall be white of color
   if (distancexy 0 -51 ) < Food-size                              ;; the new food source the same size as the old food sources
 [ set FS-number 6                                                 ;; give it the number 6
   set if-food? true                                               ;; let it have food
   set FS-scent-6 200 - distancexy 0 -51                           ;; and a food scent
ifelse reward-FS-6 >= 0.75                                         ;; if the depleted FS had high quality food,
 [ set  FS-quality  1                                              ;; the new FS also has a high quality
   set FS-energy food-energy-high ]                                ;; and high energy
 [ set  FS-quality  0                                              ;; else it has a low quality
   set FS-energy food-energy-low] ] ]                              ;; and a low energy
end

to FS-Renewal-7                                                    ;; food source 7 can reappear
    ask patch -30 -39                                              ;; at the same location as the former food source 7
 [ set pcolor white                                                ;; the food source center shall be white of color
   if (distancexy -30 -39 ) < Food-size                            ;; the new food source the same size as the old food sources
 [ set FS-number 7                                                 ;; give it the number 7
   set if-food? true                                               ;; let it have food
   set FS-scent-7 200 - distancexy -30 -39                         ;; and a food scent
ifelse reward-FS-7 >= 0.75                                         ;; if the depleted FS had high quality food,
 [ set  FS-quality  1                                              ;; the new FS also has a high quality
   set FS-energy food-energy-high ]                                ;; and high energy
 [ set  FS-quality  0                                              ;; else it has a low quality
   set FS-energy food-energy-low] ] ]                              ;; and a low energy
end

to FS-Renewal-8                                                    ;; food source 8 can reappear
    ask patch -48 -17                                              ;; at the same location as the former food source 8
 [ set pcolor white                                                ;; the food source center shall be white of color
   if (distancexy -48 -17 ) < Food-size                            ;; the new food source the same size as the old food sources
 [ set FS-number 8                                                 ;; give it the number 8
   set if-food? true                                               ;; let it have food
   set FS-scent-8 200 - distancexy -48 -17                         ;; and a food scent
ifelse reward-FS-8 >= 0.75                                         ;; if the depleted FS had high quality food,
 [ set  FS-quality  1                                              ;; the new FS also has a high quality
   set FS-energy food-energy-high ]                                ;; and high energy
 [ set  FS-quality  0                                              ;; else it has a low quality
   set FS-energy food-energy-low] ] ]                              ;; and a low energy
end

to FS-Renewal-9                                                    ;; food source 9 can reappear
    ask patch -48 15                                               ;; at the same location as the former food source 9
 [ set pcolor white                                                ;; the food source center shall be white of color
   if (distancexy -48 15 ) < Food-size                             ;; the new food source the same size as the old food sources
 [ set FS-number 9                                                 ;; give it the number 9
   set if-food? true                                               ;; let it have food
   set FS-scent-9 200 - distancexy -48 15                          ;; and a food scent
ifelse reward-FS-9 >= 0.75                                         ;; if the depleted FS had high quality food,
 [ set  FS-quality  1                                              ;; the new FS also has a high quality
   set FS-energy food-energy-high ]                                ;; and high energy
 [ set  FS-quality  0                                              ;; else it has a low quality
   set FS-energy food-energy-low] ] ]                              ;; and a low energy
end

to FS-Renewal-10                                                   ;; food source 10 can reappear
    ask patch -30 39                                               ;; at the same location as the former food source 10
 [ set pcolor white                                                ;; the food source center shall be white of color
   if (distancexy -30 39 ) < Food-size                             ;; the new food source the same size as the old food sources
 [ set FS-number 10                                                ;; give it the number 10
   set if-food? true                                               ;; let it have food
   set FS-scent-10 200 - distancexy -30 39                         ;; and a food scent
ifelse reward-FS-10 >= 0.75                                        ;; if the depleted FS had high quality food,
 [ set  FS-quality  1                                              ;; the new FS also has a high quality
   set FS-energy food-energy-high ]                                ;; and high energy
 [ set  FS-quality  0                                              ;; else it has a low quality
   set FS-energy food-energy-low] ] ]                              ;; and a low energy
end
@#$#@#$#@
GRAPHICS-WINDOW
427
12
1177
763
-1
-1
5.26241135
1
10
1
1
1
0
0
0
1
-70
70
-70
70
1
1
1
ticks
30.0

BUTTON
17
10
97
43
NIL
setup
NIL
1
T
OBSERVER
NIL
NIL
NIL
NIL
1

BUTTON
107
10
182
43
NIL
go
T
1
T
OBSERVER
NIL
NIL
NIL
NIL
0

BUTTON
195
10
289
43
go-one-tick
go
NIL
1
T
OBSERVER
NIL
NIL
NIL
NIL
1

SLIDER
18
81
191
114
Food-size
Food-size
1
10
1.0
1
1
NIL
HORIZONTAL

SLIDER
18
118
191
151
nest-size
nest-size
1
10
10.0
1
1
NIL
HORIZONTAL

SLIDER
210
46
386
79
number-of-recruits
number-of-recruits
0.0
100.0
80.0
1.0
1
NIL
HORIZONTAL

SLIDER
15
47
188
80
number-of-scouts
number-of-scouts
0.0
100.0
20.0
1.0
1
NIL
HORIZONTAL

SLIDER
24
542
197
575
velocity-nest
velocity-nest
0.01
1
0.1
0.01
1
NIL
HORIZONTAL

PLOT
23
651
226
780
Nestenergy
NIL
NIL
0.0
10.0
0.0
10.0
true
false
"" ""
PENS
"default" 1.0 0 -16777216 true "" "plot nest-energy"

SLIDER
214
576
391
609
unloading-time
unloading-time
1
120
60.0
1
1
NIL
HORIZONTAL

SLIDER
20
154
193
187
reward-FS-1
reward-FS-1
0.075
0.75
0.75
0.675
1
NIL
HORIZONTAL

SLIDER
210
154
386
187
reward-FS-2
reward-FS-2
0.075
0.75
0.75
0.675
1
NIL
HORIZONTAL

SLIDER
214
541
391
574
velocity-outside
velocity-outside
0.01
1
0.8
0.01
1
NIL
HORIZONTAL

SLIDER
21
330
195
363
recruitment-probability-1
recruitment-probability-1
1
100
50.0
1
1
NIL
HORIZONTAL

SLIDER
212
332
389
365
recruitment-probability-2
recruitment-probability-2
1
100
50.0
1
1
NIL
HORIZONTAL

SLIDER
212
118
387
151
happiness-threshold
happiness-threshold
1
100
90.0
1
1
NIL
HORIZONTAL

OUTPUT
1181
12
1433
758
12

SLIDER
24
576
198
609
metabolic-cost
metabolic-cost
0
10
2.44699E-8
0.000000000001
1
NIL
HORIZONTAL

SLIDER
23
611
199
644
food-energy-high
food-energy-high
7.5
7500
7500.0
0.75
1
NIL
HORIZONTAL

SLIDER
211
79
388
112
FS-timer
FS-timer
1
900
600.0
1
1
NIL
HORIZONTAL

SLIDER
214
507
391
540
FS-searching-time
FS-searching-time
60
600
180.0
1
1
NIL
HORIZONTAL

SLIDER
23
507
198
540
TR-breakup-probability
TR-breakup-probability
0.001
1.000
0.0
0.001
1
NIL
HORIZONTAL

SLIDER
215
611
392
644
food-energy-low
food-energy-low
0.75
750
7500.0
0.075
1
NIL
HORIZONTAL

SLIDER
20
190
191
223
reward-FS-3
reward-FS-3
0.075
0.75
0.75
0.675
1
NIL
HORIZONTAL

SLIDER
210
189
385
222
reward-FS-4
reward-FS-4
0.075
0.75
0.75
0.675
1
NIL
HORIZONTAL

SLIDER
21
225
190
258
reward-FS-5
reward-FS-5
0.075
0.75
0.75
0.675
1
NIL
HORIZONTAL

SLIDER
212
223
386
256
reward-FS-6
reward-FS-6
0.075
0.75
0.75
0.675
1
NIL
HORIZONTAL

SLIDER
19
259
191
292
reward-FS-7
reward-FS-7
0.075
0.75
0.75
0.675
1
NIL
HORIZONTAL

SLIDER
211
259
386
292
reward-FS-8
reward-FS-8
0.075
0.75
0.75
0.675
1
NIL
HORIZONTAL

SLIDER
21
294
191
327
reward-FS-9
reward-FS-9
0.075
0.75
0.75
0.675
1
NIL
HORIZONTAL

SLIDER
211
294
388
327
reward-FS-10
reward-FS-10
0.075
0.75
0.75
0.675
1
NIL
HORIZONTAL

SLIDER
21
366
195
399
recruitment-probability-3
recruitment-probability-3
1
100
50.0
1
1
NIL
HORIZONTAL

SLIDER
213
366
389
399
recruitment-probability-4
recruitment-probability-4
1
100
50.0
1
1
NIL
HORIZONTAL

SLIDER
21
401
195
434
recruitment-probability-5
recruitment-probability-5
1
100
50.0
1
1
NIL
HORIZONTAL

SLIDER
214
401
390
434
recruitment-probability-6
recruitment-probability-6
1
100
50.0
1
1
NIL
HORIZONTAL

SLIDER
22
437
195
470
recruitment-probability-7
recruitment-probability-7
1
100
50.0
1
1
NIL
HORIZONTAL

SLIDER
214
437
390
470
recruitment-probability-8
recruitment-probability-8
1
100
50.0
1
1
NIL
HORIZONTAL

SLIDER
23
472
194
505
recruitment-probability-9
recruitment-probability-9
1
100
50.0
1
1
NIL
HORIZONTAL

SLIDER
214
472
390
505
recruitment-probability-10
recruitment-probability-10
1
100
50.0
1
1
NIL
HORIZONTAL

@#$#@#$#@
## Foraging Ants


This model was created with the Netlogo software 6.1.0 and is analysed in the manuscript:

N. Goy, S. M. Glaser, C. Grüter, The adaptive value of tandem communication in ants: insights from an agent-based model. BioRxiv, 1–31 (2020).

* Wilensky, U. (1999). NetLogo. http://ccl.northwestern.edu/netlogo/. Center for Connected Learning and Computer-Based Modeling, Northwestern University, Evanston, IL.

* Wilensky, U. (1997).  NetLogo Ants model.  http://ccl.northwestern.edu/netlogo/models/Ants.  Center for Connected Learning and Computer-Based Modeling, Northwestern University, Evanston, IL.
@#$#@#$#@
default
true
0
Polygon -7500403 true true 150 5 40 250 150 205 260 250

airplane
true
0
Polygon -7500403 true true 150 0 135 15 120 60 120 105 15 165 15 195 120 180 135 240 105 270 120 285 150 270 180 285 210 270 165 240 180 180 285 195 285 165 180 105 180 60 165 15

arrow
true
0
Polygon -7500403 true true 150 0 0 150 105 150 105 293 195 293 195 150 300 150

box
false
0
Polygon -7500403 true true 150 285 285 225 285 75 150 135
Polygon -7500403 true true 150 135 15 75 150 15 285 75
Polygon -7500403 true true 15 75 15 225 150 285 150 135
Line -16777216 false 150 285 150 135
Line -16777216 false 150 135 15 75
Line -16777216 false 150 135 285 75

bug
true
0
Circle -7500403 true true 96 182 108
Circle -7500403 true true 110 127 80
Circle -7500403 true true 110 75 80
Line -7500403 true 150 100 80 30
Line -7500403 true 150 100 220 30

butterfly
true
0
Polygon -7500403 true true 150 165 209 199 225 225 225 255 195 270 165 255 150 240
Polygon -7500403 true true 150 165 89 198 75 225 75 255 105 270 135 255 150 240
Polygon -7500403 true true 139 148 100 105 55 90 25 90 10 105 10 135 25 180 40 195 85 194 139 163
Polygon -7500403 true true 162 150 200 105 245 90 275 90 290 105 290 135 275 180 260 195 215 195 162 165
Polygon -16777216 true false 150 255 135 225 120 150 135 120 150 105 165 120 180 150 165 225
Circle -16777216 true false 135 90 30
Line -16777216 false 150 105 195 60
Line -16777216 false 150 105 105 60

car
false
0
Polygon -7500403 true true 300 180 279 164 261 144 240 135 226 132 213 106 203 84 185 63 159 50 135 50 75 60 0 150 0 165 0 225 300 225 300 180
Circle -16777216 true false 180 180 90
Circle -16777216 true false 30 180 90
Polygon -16777216 true false 162 80 132 78 134 135 209 135 194 105 189 96 180 89
Circle -7500403 true true 47 195 58
Circle -7500403 true true 195 195 58

circle
false
0
Circle -7500403 true true 0 0 300

circle 2
false
0
Circle -7500403 true true 0 0 300
Circle -16777216 true false 30 30 240

cow
false
0
Polygon -7500403 true true 200 193 197 249 179 249 177 196 166 187 140 189 93 191 78 179 72 211 49 209 48 181 37 149 25 120 25 89 45 72 103 84 179 75 198 76 252 64 272 81 293 103 285 121 255 121 242 118 224 167
Polygon -7500403 true true 73 210 86 251 62 249 48 208
Polygon -7500403 true true 25 114 16 195 9 204 23 213 25 200 39 123

cylinder
false
0
Circle -7500403 true true 0 0 300

dot
false
0
Circle -7500403 true true 90 90 120

face happy
false
0
Circle -7500403 true true 8 8 285
Circle -16777216 true false 60 75 60
Circle -16777216 true false 180 75 60
Polygon -16777216 true false 150 255 90 239 62 213 47 191 67 179 90 203 109 218 150 225 192 218 210 203 227 181 251 194 236 217 212 240

face neutral
false
0
Circle -7500403 true true 8 7 285
Circle -16777216 true false 60 75 60
Circle -16777216 true false 180 75 60
Rectangle -16777216 true false 60 195 240 225

face sad
false
0
Circle -7500403 true true 8 8 285
Circle -16777216 true false 60 75 60
Circle -16777216 true false 180 75 60
Polygon -16777216 true false 150 168 90 184 62 210 47 232 67 244 90 220 109 205 150 198 192 205 210 220 227 242 251 229 236 206 212 183

fish
false
0
Polygon -1 true false 44 131 21 87 15 86 0 120 15 150 0 180 13 214 20 212 45 166
Polygon -1 true false 135 195 119 235 95 218 76 210 46 204 60 165
Polygon -1 true false 75 45 83 77 71 103 86 114 166 78 135 60
Polygon -7500403 true true 30 136 151 77 226 81 280 119 292 146 292 160 287 170 270 195 195 210 151 212 30 166
Circle -16777216 true false 215 106 30

flag
false
0
Rectangle -7500403 true true 60 15 75 300
Polygon -7500403 true true 90 150 270 90 90 30
Line -7500403 true 75 135 90 135
Line -7500403 true 75 45 90 45

flower
false
0
Polygon -10899396 true false 135 120 165 165 180 210 180 240 150 300 165 300 195 240 195 195 165 135
Circle -7500403 true true 85 132 38
Circle -7500403 true true 130 147 38
Circle -7500403 true true 192 85 38
Circle -7500403 true true 85 40 38
Circle -7500403 true true 177 40 38
Circle -7500403 true true 177 132 38
Circle -7500403 true true 70 85 38
Circle -7500403 true true 130 25 38
Circle -7500403 true true 96 51 108
Circle -16777216 true false 113 68 74
Polygon -10899396 true false 189 233 219 188 249 173 279 188 234 218
Polygon -10899396 true false 180 255 150 210 105 210 75 240 135 240

house
false
0
Rectangle -7500403 true true 45 120 255 285
Rectangle -16777216 true false 120 210 180 285
Polygon -7500403 true true 15 120 150 15 285 120
Line -16777216 false 30 120 270 120

leaf
false
0
Polygon -7500403 true true 150 210 135 195 120 210 60 210 30 195 60 180 60 165 15 135 30 120 15 105 40 104 45 90 60 90 90 105 105 120 120 120 105 60 120 60 135 30 150 15 165 30 180 60 195 60 180 120 195 120 210 105 240 90 255 90 263 104 285 105 270 120 285 135 240 165 240 180 270 195 240 210 180 210 165 195
Polygon -7500403 true true 135 195 135 240 120 255 105 255 105 285 135 285 165 240 165 195

line
true
0
Line -7500403 true 150 0 150 300

line half
true
0
Line -7500403 true 150 0 150 150

pentagon
false
0
Polygon -7500403 true true 150 15 15 120 60 285 240 285 285 120

person
false
0
Circle -7500403 true true 110 5 80
Polygon -7500403 true true 105 90 120 195 90 285 105 300 135 300 150 225 165 300 195 300 210 285 180 195 195 90
Rectangle -7500403 true true 127 79 172 94
Polygon -7500403 true true 195 90 240 150 225 180 165 105
Polygon -7500403 true true 105 90 60 150 75 180 135 105

plant
false
0
Rectangle -7500403 true true 135 90 165 300
Polygon -7500403 true true 135 255 90 210 45 195 75 255 135 285
Polygon -7500403 true true 165 255 210 210 255 195 225 255 165 285
Polygon -7500403 true true 135 180 90 135 45 120 75 180 135 210
Polygon -7500403 true true 165 180 165 210 225 180 255 120 210 135
Polygon -7500403 true true 135 105 90 60 45 45 75 105 135 135
Polygon -7500403 true true 165 105 165 135 225 105 255 45 210 60
Polygon -7500403 true true 135 90 120 45 150 15 180 45 165 90

square
false
0
Rectangle -7500403 true true 30 30 270 270

square 2
false
0
Rectangle -7500403 true true 30 30 270 270
Rectangle -16777216 true false 60 60 240 240

star
false
0
Polygon -7500403 true true 151 1 185 108 298 108 207 175 242 282 151 216 59 282 94 175 3 108 116 108

target
false
0
Circle -7500403 true true 0 0 300
Circle -16777216 true false 30 30 240
Circle -7500403 true true 60 60 180
Circle -16777216 true false 90 90 120
Circle -7500403 true true 120 120 60

tree
false
0
Circle -7500403 true true 118 3 94
Rectangle -6459832 true false 120 195 180 300
Circle -7500403 true true 65 21 108
Circle -7500403 true true 116 41 127
Circle -7500403 true true 45 90 120
Circle -7500403 true true 104 74 152

triangle
false
0
Polygon -7500403 true true 150 30 15 255 285 255

triangle 2
false
0
Polygon -7500403 true true 150 30 15 255 285 255
Polygon -16777216 true false 151 99 225 223 75 224

truck
false
0
Rectangle -7500403 true true 4 45 195 187
Polygon -7500403 true true 296 193 296 150 259 134 244 104 208 104 207 194
Rectangle -1 true false 195 60 195 105
Polygon -16777216 true false 238 112 252 141 219 141 218 112
Circle -16777216 true false 234 174 42
Rectangle -7500403 true true 181 185 214 194
Circle -16777216 true false 144 174 42
Circle -16777216 true false 24 174 42
Circle -7500403 false true 24 174 42
Circle -7500403 false true 144 174 42
Circle -7500403 false true 234 174 42

turtle
true
0
Polygon -10899396 true false 215 204 240 233 246 254 228 266 215 252 193 210
Polygon -10899396 true false 195 90 225 75 245 75 260 89 269 108 261 124 240 105 225 105 210 105
Polygon -10899396 true false 105 90 75 75 55 75 40 89 31 108 39 124 60 105 75 105 90 105
Polygon -10899396 true false 132 85 134 64 107 51 108 17 150 2 192 18 192 52 169 65 172 87
Polygon -10899396 true false 85 204 60 233 54 254 72 266 85 252 107 210
Polygon -7500403 true true 119 75 179 75 209 101 224 135 220 225 175 261 128 261 81 224 74 135 88 99

wheel
false
0
Circle -7500403 true true 3 3 294
Circle -16777216 true false 30 30 240
Line -7500403 true 150 285 150 15
Line -7500403 true 15 150 285 150
Circle -7500403 true true 120 120 60
Line -7500403 true 216 40 79 269
Line -7500403 true 40 84 269 221
Line -7500403 true 40 216 269 79
Line -7500403 true 84 40 221 269

x
false
0
Polygon -7500403 true true 270 75 225 30 30 225 75 270
Polygon -7500403 true true 30 75 75 30 270 225 225 270
@#$#@#$#@
NetLogo 6.1.0
@#$#@#$#@
@#$#@#$#@
@#$#@#$#@
<experiments>
  <experiment name="ratio 0.5_High_stable" repetitions="30" sequentialRunOrder="false" runMetricsEveryStep="false">
    <setup>setup</setup>
    <go>go</go>
    <timeLimit steps="5400"/>
    <metric>nest-energy</metric>
    <enumeratedValueSet variable="number-of-scouts">
      <value value="50"/>
    </enumeratedValueSet>
    <enumeratedValueSet variable="number-of-recruits">
      <value value="50"/>
    </enumeratedValueSet>
    <enumeratedValueSet variable="nest-size">
      <value value="10"/>
    </enumeratedValueSet>
    <enumeratedValueSet variable="Food-size">
      <value value="1"/>
    </enumeratedValueSet>
    <enumeratedValueSet variable="reward-FS-1">
      <value value="0.75"/>
    </enumeratedValueSet>
    <enumeratedValueSet variable="reward-FS-2">
      <value value="0.75"/>
    </enumeratedValueSet>
    <enumeratedValueSet variable="reward-FS-3">
      <value value="0.75"/>
    </enumeratedValueSet>
    <enumeratedValueSet variable="reward-FS-4">
      <value value="0.75"/>
    </enumeratedValueSet>
    <enumeratedValueSet variable="reward-FS-5">
      <value value="0.75"/>
    </enumeratedValueSet>
    <enumeratedValueSet variable="reward-FS-6">
      <value value="0.75"/>
    </enumeratedValueSet>
    <enumeratedValueSet variable="reward-FS-7">
      <value value="0.75"/>
    </enumeratedValueSet>
    <enumeratedValueSet variable="reward-FS-8">
      <value value="0.75"/>
    </enumeratedValueSet>
    <enumeratedValueSet variable="reward-FS-9">
      <value value="0.75"/>
    </enumeratedValueSet>
    <enumeratedValueSet variable="reward-FS-10">
      <value value="0.75"/>
    </enumeratedValueSet>
    <enumeratedValueSet variable="FS-timer">
      <value value="600"/>
    </enumeratedValueSet>
    <enumeratedValueSet variable="food-energy-high">
      <value value="7500"/>
    </enumeratedValueSet>
    <enumeratedValueSet variable="food-energy-low">
      <value value="750"/>
    </enumeratedValueSet>
    <enumeratedValueSet variable="velocity-nest">
      <value value="0.1"/>
    </enumeratedValueSet>
    <enumeratedValueSet variable="velocity-outside">
      <value value="0.8"/>
    </enumeratedValueSet>
    <enumeratedValueSet variable="FS-searching-time">
      <value value="180"/>
    </enumeratedValueSet>
    <enumeratedValueSet variable="unloading-time">
      <value value="60"/>
    </enumeratedValueSet>
    <enumeratedValueSet variable="recruitment-probability-1">
      <value value="50"/>
    </enumeratedValueSet>
    <enumeratedValueSet variable="recruitment-probability-2">
      <value value="50"/>
    </enumeratedValueSet>
    <enumeratedValueSet variable="recruitment-probability-3">
      <value value="50"/>
    </enumeratedValueSet>
    <enumeratedValueSet variable="recruitment-probability-4">
      <value value="50"/>
    </enumeratedValueSet>
    <enumeratedValueSet variable="recruitment-probability-5">
      <value value="50"/>
    </enumeratedValueSet>
    <enumeratedValueSet variable="recruitment-probability-6">
      <value value="50"/>
    </enumeratedValueSet>
    <enumeratedValueSet variable="recruitment-probability-7">
      <value value="50"/>
    </enumeratedValueSet>
    <enumeratedValueSet variable="recruitment-probability-8">
      <value value="50"/>
    </enumeratedValueSet>
    <enumeratedValueSet variable="recruitment-probability-9">
      <value value="50"/>
    </enumeratedValueSet>
    <enumeratedValueSet variable="recruitment-probability-10">
      <value value="50"/>
    </enumeratedValueSet>
    <enumeratedValueSet variable="TR-breakup-probability">
      <value value="0.005"/>
    </enumeratedValueSet>
    <enumeratedValueSet variable="metabolic-cost">
      <value value="2.44699E-7"/>
    </enumeratedValueSet>
    <enumeratedValueSet variable="happiness-threshold">
      <value value="90"/>
    </enumeratedValueSet>
  </experiment>
  <experiment name="No breakup 1:4 unstable HQ" repetitions="30" runMetricsEveryStep="false">
    <setup>setup</setup>
    <go>go</go>
    <timeLimit steps="5400"/>
    <metric>nest-energy</metric>
    <enumeratedValueSet variable="TR-breakup-probability">
      <value value="0"/>
    </enumeratedValueSet>
    <enumeratedValueSet variable="number-of-scouts">
      <value value="20"/>
    </enumeratedValueSet>
    <enumeratedValueSet variable="number-of-recruits">
      <value value="80"/>
    </enumeratedValueSet>
    <enumeratedValueSet variable="metabolic-cost">
      <value value="2.44699E-7"/>
    </enumeratedValueSet>
    <enumeratedValueSet variable="FS-timer">
      <value value="600"/>
    </enumeratedValueSet>
    <enumeratedValueSet variable="food-energy-low">
      <value value="0.75"/>
    </enumeratedValueSet>
    <enumeratedValueSet variable="velocity-outside">
      <value value="0.8"/>
    </enumeratedValueSet>
    <enumeratedValueSet variable="FS-searching-time">
      <value value="180"/>
    </enumeratedValueSet>
    <enumeratedValueSet variable="reward-FS-1">
      <value value="0.75"/>
    </enumeratedValueSet>
    <enumeratedValueSet variable="reward-FS-2">
      <value value="0.75"/>
    </enumeratedValueSet>
    <enumeratedValueSet variable="reward-FS-3">
      <value value="0.75"/>
    </enumeratedValueSet>
    <enumeratedValueSet variable="reward-FS-4">
      <value value="0.75"/>
    </enumeratedValueSet>
    <enumeratedValueSet variable="reward-FS-5">
      <value value="0.75"/>
    </enumeratedValueSet>
    <enumeratedValueSet variable="reward-FS-6">
      <value value="0.75"/>
    </enumeratedValueSet>
    <enumeratedValueSet variable="reward-FS-7">
      <value value="0.75"/>
    </enumeratedValueSet>
    <enumeratedValueSet variable="reward-FS-8">
      <value value="0.75"/>
    </enumeratedValueSet>
    <enumeratedValueSet variable="reward-FS-9">
      <value value="0.75"/>
    </enumeratedValueSet>
    <enumeratedValueSet variable="reward-FS-10">
      <value value="0.75"/>
    </enumeratedValueSet>
    <enumeratedValueSet variable="unloading-time">
      <value value="60"/>
    </enumeratedValueSet>
    <enumeratedValueSet variable="velocity-nest">
      <value value="0.1"/>
    </enumeratedValueSet>
    <enumeratedValueSet variable="recruitment-probability-1">
      <value value="50"/>
    </enumeratedValueSet>
    <enumeratedValueSet variable="recruitment-probability-2">
      <value value="50"/>
    </enumeratedValueSet>
    <enumeratedValueSet variable="recruitment-probability-3">
      <value value="50"/>
    </enumeratedValueSet>
    <enumeratedValueSet variable="recruitment-probability-4">
      <value value="50"/>
    </enumeratedValueSet>
    <enumeratedValueSet variable="recruitment-probability-5">
      <value value="50"/>
    </enumeratedValueSet>
    <enumeratedValueSet variable="recruitment-probability-6">
      <value value="50"/>
    </enumeratedValueSet>
    <enumeratedValueSet variable="recruitment-probability-7">
      <value value="50"/>
    </enumeratedValueSet>
    <enumeratedValueSet variable="recruitment-probability-8">
      <value value="50"/>
    </enumeratedValueSet>
    <enumeratedValueSet variable="recruitment-probability-9">
      <value value="50"/>
    </enumeratedValueSet>
    <enumeratedValueSet variable="recruitment-probability-10">
      <value value="50"/>
    </enumeratedValueSet>
    <enumeratedValueSet variable="happiness-threshold">
      <value value="90"/>
    </enumeratedValueSet>
    <enumeratedValueSet variable="nest-size">
      <value value="10"/>
    </enumeratedValueSet>
    <enumeratedValueSet variable="food-energy-high">
      <value value="7.5"/>
    </enumeratedValueSet>
    <enumeratedValueSet variable="Food-size">
      <value value="1"/>
    </enumeratedValueSet>
  </experiment>
  <experiment name="No breakup 1:4 stable mixed" repetitions="30" runMetricsEveryStep="false">
    <setup>setup</setup>
    <go>go</go>
    <timeLimit steps="5400"/>
    <metric>nest-energy</metric>
    <enumeratedValueSet variable="TR-breakup-probability">
      <value value="0"/>
    </enumeratedValueSet>
    <enumeratedValueSet variable="number-of-scouts">
      <value value="20"/>
    </enumeratedValueSet>
    <enumeratedValueSet variable="number-of-recruits">
      <value value="80"/>
    </enumeratedValueSet>
    <enumeratedValueSet variable="metabolic-cost">
      <value value="2.44699E-7"/>
    </enumeratedValueSet>
    <enumeratedValueSet variable="FS-timer">
      <value value="600"/>
    </enumeratedValueSet>
    <enumeratedValueSet variable="food-energy-low">
      <value value="750"/>
    </enumeratedValueSet>
    <enumeratedValueSet variable="velocity-outside">
      <value value="0.8"/>
    </enumeratedValueSet>
    <enumeratedValueSet variable="FS-searching-time">
      <value value="180"/>
    </enumeratedValueSet>
    <enumeratedValueSet variable="reward-FS-1">
      <value value="0.75"/>
    </enumeratedValueSet>
    <enumeratedValueSet variable="reward-FS-2">
      <value value="0.075"/>
    </enumeratedValueSet>
    <enumeratedValueSet variable="reward-FS-3">
      <value value="0.75"/>
    </enumeratedValueSet>
    <enumeratedValueSet variable="reward-FS-4">
      <value value="0.075"/>
    </enumeratedValueSet>
    <enumeratedValueSet variable="reward-FS-5">
      <value value="0.75"/>
    </enumeratedValueSet>
    <enumeratedValueSet variable="reward-FS-6">
      <value value="0.075"/>
    </enumeratedValueSet>
    <enumeratedValueSet variable="reward-FS-7">
      <value value="0.75"/>
    </enumeratedValueSet>
    <enumeratedValueSet variable="reward-FS-8">
      <value value="0.075"/>
    </enumeratedValueSet>
    <enumeratedValueSet variable="reward-FS-9">
      <value value="0.75"/>
    </enumeratedValueSet>
    <enumeratedValueSet variable="reward-FS-10">
      <value value="0.075"/>
    </enumeratedValueSet>
    <enumeratedValueSet variable="unloading-time">
      <value value="60"/>
    </enumeratedValueSet>
    <enumeratedValueSet variable="velocity-nest">
      <value value="0.1"/>
    </enumeratedValueSet>
    <enumeratedValueSet variable="recruitment-probability-1">
      <value value="50"/>
    </enumeratedValueSet>
    <enumeratedValueSet variable="recruitment-probability-2">
      <value value="50"/>
    </enumeratedValueSet>
    <enumeratedValueSet variable="recruitment-probability-3">
      <value value="50"/>
    </enumeratedValueSet>
    <enumeratedValueSet variable="recruitment-probability-4">
      <value value="50"/>
    </enumeratedValueSet>
    <enumeratedValueSet variable="recruitment-probability-5">
      <value value="50"/>
    </enumeratedValueSet>
    <enumeratedValueSet variable="recruitment-probability-6">
      <value value="50"/>
    </enumeratedValueSet>
    <enumeratedValueSet variable="recruitment-probability-7">
      <value value="50"/>
    </enumeratedValueSet>
    <enumeratedValueSet variable="recruitment-probability-8">
      <value value="50"/>
    </enumeratedValueSet>
    <enumeratedValueSet variable="recruitment-probability-9">
      <value value="50"/>
    </enumeratedValueSet>
    <enumeratedValueSet variable="recruitment-probability-10">
      <value value="50"/>
    </enumeratedValueSet>
    <enumeratedValueSet variable="happiness-threshold">
      <value value="90"/>
    </enumeratedValueSet>
    <enumeratedValueSet variable="nest-size">
      <value value="10"/>
    </enumeratedValueSet>
    <enumeratedValueSet variable="food-energy-high">
      <value value="7500"/>
    </enumeratedValueSet>
    <enumeratedValueSet variable="Food-size">
      <value value="1"/>
    </enumeratedValueSet>
  </experiment>
  <experiment name="No breakup 1:4 unstable mixed" repetitions="30" runMetricsEveryStep="false">
    <setup>setup</setup>
    <go>go</go>
    <timeLimit steps="5400"/>
    <metric>nest-energy</metric>
    <enumeratedValueSet variable="TR-breakup-probability">
      <value value="0"/>
    </enumeratedValueSet>
    <enumeratedValueSet variable="number-of-scouts">
      <value value="20"/>
    </enumeratedValueSet>
    <enumeratedValueSet variable="number-of-recruits">
      <value value="80"/>
    </enumeratedValueSet>
    <enumeratedValueSet variable="metabolic-cost">
      <value value="2.44699E-7"/>
    </enumeratedValueSet>
    <enumeratedValueSet variable="FS-timer">
      <value value="600"/>
    </enumeratedValueSet>
    <enumeratedValueSet variable="food-energy-low">
      <value value="0.75"/>
    </enumeratedValueSet>
    <enumeratedValueSet variable="velocity-outside">
      <value value="0.8"/>
    </enumeratedValueSet>
    <enumeratedValueSet variable="FS-searching-time">
      <value value="180"/>
    </enumeratedValueSet>
    <enumeratedValueSet variable="reward-FS-1">
      <value value="0.75"/>
    </enumeratedValueSet>
    <enumeratedValueSet variable="reward-FS-2">
      <value value="0.075"/>
    </enumeratedValueSet>
    <enumeratedValueSet variable="reward-FS-3">
      <value value="0.75"/>
    </enumeratedValueSet>
    <enumeratedValueSet variable="reward-FS-4">
      <value value="0.075"/>
    </enumeratedValueSet>
    <enumeratedValueSet variable="reward-FS-5">
      <value value="0.75"/>
    </enumeratedValueSet>
    <enumeratedValueSet variable="reward-FS-6">
      <value value="0.075"/>
    </enumeratedValueSet>
    <enumeratedValueSet variable="reward-FS-7">
      <value value="0.75"/>
    </enumeratedValueSet>
    <enumeratedValueSet variable="reward-FS-8">
      <value value="0.075"/>
    </enumeratedValueSet>
    <enumeratedValueSet variable="reward-FS-9">
      <value value="0.75"/>
    </enumeratedValueSet>
    <enumeratedValueSet variable="reward-FS-10">
      <value value="0.075"/>
    </enumeratedValueSet>
    <enumeratedValueSet variable="unloading-time">
      <value value="60"/>
    </enumeratedValueSet>
    <enumeratedValueSet variable="velocity-nest">
      <value value="0.1"/>
    </enumeratedValueSet>
    <enumeratedValueSet variable="recruitment-probability-1">
      <value value="50"/>
    </enumeratedValueSet>
    <enumeratedValueSet variable="recruitment-probability-2">
      <value value="50"/>
    </enumeratedValueSet>
    <enumeratedValueSet variable="recruitment-probability-3">
      <value value="50"/>
    </enumeratedValueSet>
    <enumeratedValueSet variable="recruitment-probability-4">
      <value value="50"/>
    </enumeratedValueSet>
    <enumeratedValueSet variable="recruitment-probability-5">
      <value value="50"/>
    </enumeratedValueSet>
    <enumeratedValueSet variable="recruitment-probability-6">
      <value value="50"/>
    </enumeratedValueSet>
    <enumeratedValueSet variable="recruitment-probability-7">
      <value value="50"/>
    </enumeratedValueSet>
    <enumeratedValueSet variable="recruitment-probability-8">
      <value value="50"/>
    </enumeratedValueSet>
    <enumeratedValueSet variable="recruitment-probability-9">
      <value value="50"/>
    </enumeratedValueSet>
    <enumeratedValueSet variable="recruitment-probability-10">
      <value value="50"/>
    </enumeratedValueSet>
    <enumeratedValueSet variable="happiness-threshold">
      <value value="90"/>
    </enumeratedValueSet>
    <enumeratedValueSet variable="nest-size">
      <value value="10"/>
    </enumeratedValueSet>
    <enumeratedValueSet variable="food-energy-high">
      <value value="7.5"/>
    </enumeratedValueSet>
    <enumeratedValueSet variable="Food-size">
      <value value="1"/>
    </enumeratedValueSet>
  </experiment>
  <experiment name="No breakup 1:4 mixed long" repetitions="30" runMetricsEveryStep="false">
    <setup>setup</setup>
    <go>go</go>
    <timeLimit steps="21600"/>
    <metric>nest-energy</metric>
    <enumeratedValueSet variable="TR-breakup-probability">
      <value value="0"/>
    </enumeratedValueSet>
    <enumeratedValueSet variable="number-of-scouts">
      <value value="20"/>
    </enumeratedValueSet>
    <enumeratedValueSet variable="number-of-recruits">
      <value value="80"/>
    </enumeratedValueSet>
    <enumeratedValueSet variable="metabolic-cost">
      <value value="2.44699E-7"/>
    </enumeratedValueSet>
    <enumeratedValueSet variable="FS-timer">
      <value value="600"/>
    </enumeratedValueSet>
    <enumeratedValueSet variable="food-energy-low">
      <value value="750"/>
    </enumeratedValueSet>
    <enumeratedValueSet variable="velocity-outside">
      <value value="0.8"/>
    </enumeratedValueSet>
    <enumeratedValueSet variable="FS-searching-time">
      <value value="180"/>
    </enumeratedValueSet>
    <enumeratedValueSet variable="reward-FS-1">
      <value value="0.75"/>
    </enumeratedValueSet>
    <enumeratedValueSet variable="reward-FS-2">
      <value value="0.075"/>
    </enumeratedValueSet>
    <enumeratedValueSet variable="reward-FS-3">
      <value value="0.75"/>
    </enumeratedValueSet>
    <enumeratedValueSet variable="reward-FS-4">
      <value value="0.075"/>
    </enumeratedValueSet>
    <enumeratedValueSet variable="reward-FS-5">
      <value value="0.75"/>
    </enumeratedValueSet>
    <enumeratedValueSet variable="reward-FS-6">
      <value value="0.075"/>
    </enumeratedValueSet>
    <enumeratedValueSet variable="reward-FS-7">
      <value value="0.75"/>
    </enumeratedValueSet>
    <enumeratedValueSet variable="reward-FS-8">
      <value value="0.075"/>
    </enumeratedValueSet>
    <enumeratedValueSet variable="reward-FS-9">
      <value value="0.75"/>
    </enumeratedValueSet>
    <enumeratedValueSet variable="reward-FS-10">
      <value value="0.075"/>
    </enumeratedValueSet>
    <enumeratedValueSet variable="unloading-time">
      <value value="60"/>
    </enumeratedValueSet>
    <enumeratedValueSet variable="velocity-nest">
      <value value="0.1"/>
    </enumeratedValueSet>
    <enumeratedValueSet variable="recruitment-probability-1">
      <value value="50"/>
    </enumeratedValueSet>
    <enumeratedValueSet variable="recruitment-probability-2">
      <value value="50"/>
    </enumeratedValueSet>
    <enumeratedValueSet variable="recruitment-probability-3">
      <value value="50"/>
    </enumeratedValueSet>
    <enumeratedValueSet variable="recruitment-probability-4">
      <value value="50"/>
    </enumeratedValueSet>
    <enumeratedValueSet variable="recruitment-probability-5">
      <value value="50"/>
    </enumeratedValueSet>
    <enumeratedValueSet variable="recruitment-probability-6">
      <value value="50"/>
    </enumeratedValueSet>
    <enumeratedValueSet variable="recruitment-probability-7">
      <value value="50"/>
    </enumeratedValueSet>
    <enumeratedValueSet variable="recruitment-probability-8">
      <value value="50"/>
    </enumeratedValueSet>
    <enumeratedValueSet variable="recruitment-probability-9">
      <value value="50"/>
    </enumeratedValueSet>
    <enumeratedValueSet variable="recruitment-probability-10">
      <value value="50"/>
    </enumeratedValueSet>
    <enumeratedValueSet variable="happiness-threshold">
      <value value="90"/>
    </enumeratedValueSet>
    <enumeratedValueSet variable="nest-size">
      <value value="10"/>
    </enumeratedValueSet>
    <enumeratedValueSet variable="food-energy-high">
      <value value="7500"/>
    </enumeratedValueSet>
    <enumeratedValueSet variable="Food-size">
      <value value="1"/>
    </enumeratedValueSet>
  </experiment>
  <experiment name="No breakup 1:4 stable HQ long" repetitions="30" runMetricsEveryStep="false">
    <setup>setup</setup>
    <go>go</go>
    <timeLimit steps="21600"/>
    <metric>nest-energy</metric>
    <enumeratedValueSet variable="TR-breakup-probability">
      <value value="0"/>
    </enumeratedValueSet>
    <enumeratedValueSet variable="number-of-scouts">
      <value value="20"/>
    </enumeratedValueSet>
    <enumeratedValueSet variable="number-of-recruits">
      <value value="80"/>
    </enumeratedValueSet>
    <enumeratedValueSet variable="metabolic-cost">
      <value value="2.44699E-7"/>
    </enumeratedValueSet>
    <enumeratedValueSet variable="FS-timer">
      <value value="600"/>
    </enumeratedValueSet>
    <enumeratedValueSet variable="food-energy-low">
      <value value="750"/>
    </enumeratedValueSet>
    <enumeratedValueSet variable="velocity-outside">
      <value value="0.8"/>
    </enumeratedValueSet>
    <enumeratedValueSet variable="FS-searching-time">
      <value value="180"/>
    </enumeratedValueSet>
    <enumeratedValueSet variable="reward-FS-1">
      <value value="0.75"/>
    </enumeratedValueSet>
    <enumeratedValueSet variable="reward-FS-2">
      <value value="0.75"/>
    </enumeratedValueSet>
    <enumeratedValueSet variable="reward-FS-3">
      <value value="0.75"/>
    </enumeratedValueSet>
    <enumeratedValueSet variable="reward-FS-4">
      <value value="0.75"/>
    </enumeratedValueSet>
    <enumeratedValueSet variable="reward-FS-5">
      <value value="0.75"/>
    </enumeratedValueSet>
    <enumeratedValueSet variable="reward-FS-6">
      <value value="0.75"/>
    </enumeratedValueSet>
    <enumeratedValueSet variable="reward-FS-7">
      <value value="0.75"/>
    </enumeratedValueSet>
    <enumeratedValueSet variable="reward-FS-8">
      <value value="0.75"/>
    </enumeratedValueSet>
    <enumeratedValueSet variable="reward-FS-9">
      <value value="0.75"/>
    </enumeratedValueSet>
    <enumeratedValueSet variable="reward-FS-10">
      <value value="0.75"/>
    </enumeratedValueSet>
    <enumeratedValueSet variable="unloading-time">
      <value value="60"/>
    </enumeratedValueSet>
    <enumeratedValueSet variable="velocity-nest">
      <value value="0.1"/>
    </enumeratedValueSet>
    <enumeratedValueSet variable="recruitment-probability-1">
      <value value="50"/>
    </enumeratedValueSet>
    <enumeratedValueSet variable="recruitment-probability-2">
      <value value="50"/>
    </enumeratedValueSet>
    <enumeratedValueSet variable="recruitment-probability-3">
      <value value="50"/>
    </enumeratedValueSet>
    <enumeratedValueSet variable="recruitment-probability-4">
      <value value="50"/>
    </enumeratedValueSet>
    <enumeratedValueSet variable="recruitment-probability-5">
      <value value="50"/>
    </enumeratedValueSet>
    <enumeratedValueSet variable="recruitment-probability-6">
      <value value="50"/>
    </enumeratedValueSet>
    <enumeratedValueSet variable="recruitment-probability-7">
      <value value="50"/>
    </enumeratedValueSet>
    <enumeratedValueSet variable="recruitment-probability-8">
      <value value="50"/>
    </enumeratedValueSet>
    <enumeratedValueSet variable="recruitment-probability-9">
      <value value="50"/>
    </enumeratedValueSet>
    <enumeratedValueSet variable="recruitment-probability-10">
      <value value="50"/>
    </enumeratedValueSet>
    <enumeratedValueSet variable="happiness-threshold">
      <value value="90"/>
    </enumeratedValueSet>
    <enumeratedValueSet variable="nest-size">
      <value value="10"/>
    </enumeratedValueSet>
    <enumeratedValueSet variable="food-energy-high">
      <value value="7500"/>
    </enumeratedValueSet>
    <enumeratedValueSet variable="Food-size">
      <value value="1"/>
    </enumeratedValueSet>
  </experiment>
  <experiment name="No breakup 1:4 stable HQ metabolic rate" repetitions="30" runMetricsEveryStep="false">
    <setup>setup</setup>
    <go>go</go>
    <timeLimit steps="5400"/>
    <metric>nest-energy</metric>
    <enumeratedValueSet variable="TR-breakup-probability">
      <value value="0"/>
    </enumeratedValueSet>
    <enumeratedValueSet variable="number-of-scouts">
      <value value="20"/>
    </enumeratedValueSet>
    <enumeratedValueSet variable="number-of-recruits">
      <value value="80"/>
    </enumeratedValueSet>
    <enumeratedValueSet variable="metabolic-cost">
      <value value="2.44699E-6"/>
      <value value="2.44699E-8"/>
    </enumeratedValueSet>
    <enumeratedValueSet variable="FS-timer">
      <value value="600"/>
    </enumeratedValueSet>
    <enumeratedValueSet variable="food-energy-low">
      <value value="7500"/>
    </enumeratedValueSet>
    <enumeratedValueSet variable="velocity-outside">
      <value value="0.8"/>
    </enumeratedValueSet>
    <enumeratedValueSet variable="FS-searching-time">
      <value value="180"/>
    </enumeratedValueSet>
    <enumeratedValueSet variable="reward-FS-1">
      <value value="0.75"/>
    </enumeratedValueSet>
    <enumeratedValueSet variable="reward-FS-2">
      <value value="0.75"/>
    </enumeratedValueSet>
    <enumeratedValueSet variable="reward-FS-3">
      <value value="0.75"/>
    </enumeratedValueSet>
    <enumeratedValueSet variable="reward-FS-4">
      <value value="0.75"/>
    </enumeratedValueSet>
    <enumeratedValueSet variable="reward-FS-5">
      <value value="0.75"/>
    </enumeratedValueSet>
    <enumeratedValueSet variable="reward-FS-6">
      <value value="0.75"/>
    </enumeratedValueSet>
    <enumeratedValueSet variable="reward-FS-7">
      <value value="0.75"/>
    </enumeratedValueSet>
    <enumeratedValueSet variable="reward-FS-8">
      <value value="0.75"/>
    </enumeratedValueSet>
    <enumeratedValueSet variable="reward-FS-9">
      <value value="0.75"/>
    </enumeratedValueSet>
    <enumeratedValueSet variable="reward-FS-10">
      <value value="0.75"/>
    </enumeratedValueSet>
    <enumeratedValueSet variable="unloading-time">
      <value value="60"/>
    </enumeratedValueSet>
    <enumeratedValueSet variable="velocity-nest">
      <value value="0.1"/>
    </enumeratedValueSet>
    <enumeratedValueSet variable="recruitment-probability-1">
      <value value="50"/>
    </enumeratedValueSet>
    <enumeratedValueSet variable="recruitment-probability-2">
      <value value="50"/>
    </enumeratedValueSet>
    <enumeratedValueSet variable="recruitment-probability-3">
      <value value="50"/>
    </enumeratedValueSet>
    <enumeratedValueSet variable="recruitment-probability-4">
      <value value="50"/>
    </enumeratedValueSet>
    <enumeratedValueSet variable="recruitment-probability-5">
      <value value="50"/>
    </enumeratedValueSet>
    <enumeratedValueSet variable="recruitment-probability-6">
      <value value="50"/>
    </enumeratedValueSet>
    <enumeratedValueSet variable="recruitment-probability-7">
      <value value="50"/>
    </enumeratedValueSet>
    <enumeratedValueSet variable="recruitment-probability-8">
      <value value="50"/>
    </enumeratedValueSet>
    <enumeratedValueSet variable="recruitment-probability-9">
      <value value="50"/>
    </enumeratedValueSet>
    <enumeratedValueSet variable="recruitment-probability-10">
      <value value="50"/>
    </enumeratedValueSet>
    <enumeratedValueSet variable="happiness-threshold">
      <value value="90"/>
    </enumeratedValueSet>
    <enumeratedValueSet variable="nest-size">
      <value value="10"/>
    </enumeratedValueSet>
    <enumeratedValueSet variable="food-energy-high">
      <value value="7500"/>
    </enumeratedValueSet>
    <enumeratedValueSet variable="Food-size">
      <value value="1"/>
    </enumeratedValueSet>
  </experiment>
</experiments>
@#$#@#$#@
@#$#@#$#@
default
0.0
-0.2 0 0.0 1.0
0.0 1 1.0 0.0
0.2 0 0.0 1.0
link direction
true
0
Line -7500403 true 150 150 90 180
Line -7500403 true 150 150 210 180
@#$#@#$#@
0
@#$#@#$#@
